# Biophysical basis of cellular multi-specificity encoded in a model molecular switch

**DOI:** 10.1101/2020.01.04.893909

**Authors:** Tina Perica, Christopher J. P. Mathy, Jiewei Xu, Gwendolyn M. Jang, Yang Zhang, Robyn Kaake, Noah Ollikainen, Hannes Braberg, Danielle L. Swaney, David G. Lambright, Mark J. S. Kelly, Nevan J. Krogan, Tanja Kortemme

## Abstract

Molecular switches are central to signal transduction in protein interaction networks. One switch protein can independently regulate distinct cellular processes, but the molecular mechanisms enabling this functional multi-specificity remain unclear. Here we integrate system-scale cellular and biophysical measurements to study how a paradigm switch, the small GTPase Ran/Gsp1, achieves its functional multi-specificity. We make 55 targeted point mutations to individual interactions of Ran/Gsp1 and show through quantitative, systematic genetic and physical interaction mapping that Ran/Gsp1 interface perturbations have widespread cellular consequences that cluster by biological processes but, unexpectedly, not by the targeted interactions. Instead, the cellular consequences of the interface mutations group by their biophysical effects on kinetic parameters of the GTPase switch cycle, and cycle kinetics are allosterically tuned by distal interface mutations. We propose that the functional multi-specificity of Ran/Gsp1 is encoded by a differential sensitivity of biological processes to different kinetic parameters of the Gsp1 switch cycle, and that Gsp1 partners binding to the sites of distal mutations act as allosteric regulators of the switch. Similar mechanisms may underlie biological regulation by other GTPases and biological switches. Finally, our integrative platform to determine the quantitative consequences of cellular perturbations may help explain the effects of disease mutations targeting central switches.

---

Proteins perform their cellular functions within networks of interactions with many partners (1, 2). This complexity raises the fundamental question of functional specificity: How can different functions be controlled individually with the required precision and accuracy, when distinct cellular processes are interconnected and often even share common regulators? Moreover, in highly interconnected networks even a small perturbation targeting individual interactions, introduced by posttranslational modifications, point mutations, or drug binding, could be magnified through the network and have widespread cellular consequences. Protein mutations in disease are enriched in protein-protein interfaces (3, 4), but it is unclear whether the consequences of these mutations can be explained primarily by their effects on individual interactions. Similarly, drug compounds are typically designed against specific targets but could affect cellular functions more broadly. Determining the extent and mechanism by which molecular perturbations affect interconnected biological processes requires an approach that quantifies effects on both the cellular network and on the molecular functions of the targeted protein **(Fig. 1a)**.

**Fig. 1.**
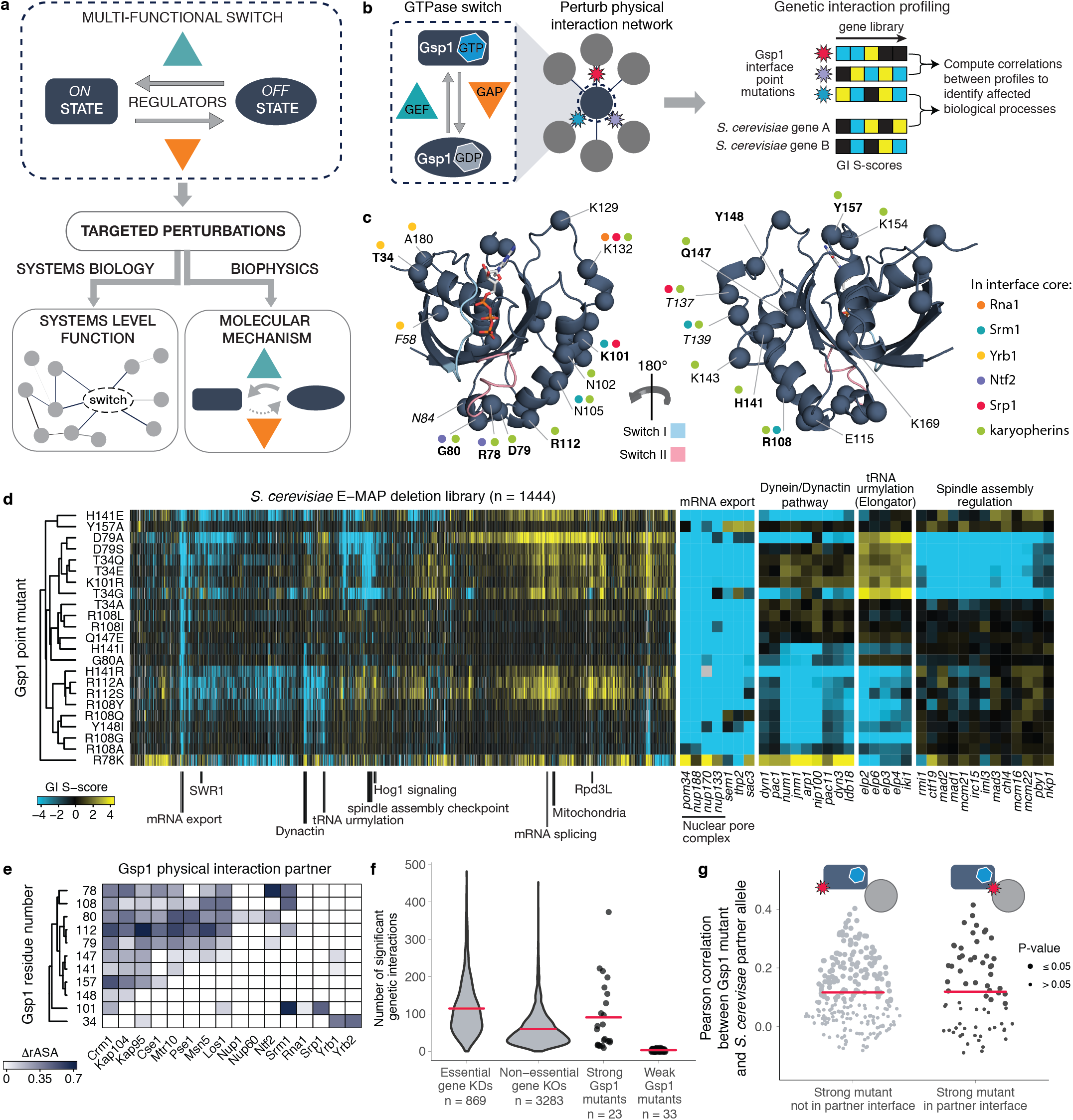
Genetic interaction (GI) profiles of Gsp1 interface point mutants cluster by biological processes but not by targeted interfaces. **a,** Schematic summary of approach combining systems-level and biophysical measurements to characterize functional multi-specificity of a biological switch motif. **b,** Interface point mutations enable probing of the biological functions of the multi-specific GTPase switch Gsp1. **c,** Structure of GTP-bound Gsp1 (navy, PDB ID: 3m1i). The 24 mutated Gsp1 residue positions are shown as Cα atom spheres. Positions of mutations with strong genetic interaction profiles are in bold, and positions that are not conserved in sequence between *S. cerevisiae* Gsp1 and human Ran are in italic. Coloured dots represent the interaction partners for which the residue is in the interface core. Switch I and II regions are shown in light blue and pale pink, respectively. **d,** GI profiles of 23 Gsp1 mutants with nine or more significant GIs. Negative S-score (blue) represents synthetic sick/lethal GIs, positive S-score (yellow) represents suppressive/epistatic GIs. Mutants and genes are hierarchically clustered by Pearson correlation. **e,** Locations of mutated residues in structurally characterized interfaces. ∆rASA is the difference in accessible surface area of a residue upon binding, relative to an empirical maximum for the solvent accessible surface area of each amino acid residue type computed as in (5). **f,** Distributions of significant (see Methods) GIs of Gsp1 point mutants compared to GIs of mutant alleles of essential and non-essential genes. Red bars indicate the mean. **g,** Distributions of Pearson correlations between the GI profiles of Gsp1 interaction partners and Gsp1 mutants if mutation is (right, black) or is not (left, gray) in the interface with that partner. Point size indicates the false discovery rate adjusted one-sided (positive) p-value of the Pearson correlation. Red dots and bars indicate the mean and the upper and lower quartile, respectively.

To develop such an approach, we targeted a central molecular switch, a GTPase. GTPases belong to a class of common biological motifs, where a two-state switch is controlled by regulators with opposing functions (6, 7) (**Fig. 1a**). For GTPases, the two states of the switch are defined by the conformation of the GTPase in either the GTP-or GDP-bound forms, and the interconversion between the two states is catalyzed by guanine nucleotide exchange factors (GEFs) and GTPase-activating proteins (GAPs) (**Fig. 1b**). Other, similar biological switch motifs involve covalent modifications controlled by opposing kinase/phosphatase or acetylase/deacetylase regulators. One striking feature of such motifs is the potential for ultrasensitive response to regulation, where small changes in the activity of the regulators can lead to sharp changes in the state of the switch (8). Moreover, switch motifs are often multi-specific, defined here as regulating several different processes (9). This multi-specificity raises the question of how a single switch motif differentially controls diverse processes at the cellular level.

In this study, we sought to uncover the mechanistic basis of functional multi-specificity in the small GTPase Gsp1 (the *S. cerevisiae* homolog of human Ran, which shares 83% amino acid identity with Gsp1), which is a single molecular switch with one main GEF and one main GAP (10). Gsp1 regulates nucleocytoplasmic transport of proteins (11, 12) and RNA (13, 14), cell cycle progression (15), RNA processing (16) and nuclear envelope assembly (17). Gsp1/Ran forms direct physical interactions with a large number of partners, and high-resolution crystal structures of Gsp1/Ran in complex with 16 different binding partners are known (**Fig. S1, Supplementary File 1 Fig. 1 and Table 1**). We reasoned that by placing defined point mutations in Gsp1 interfaces with these partners, we could differentially perturb subsets of biological processes regulated by Gsp1. We then determined the functional consequences of these Gsp1 mutations on diverse biological processes in *S. cerevisiae* using quantitative genetic interaction mapping, measured changes to the physical interaction network using affinity purification mass spectrometry (AP-MS), and finally quantified molecular changes on the Gsp1 switch motif using biophysical studies *in vitro* (**Fig. 1 a, b**).

## Targeted perturbations to GTPase interaction interfaces

To target each of the 16 known interaction interfaces of Gsp1, we designed 55 *S. cerevisiae* strains with genomically integrated point mutations in the GSP1 gene (**Fig. 1c, Fig. S1, Supplementary File 1 Fig. 1 and Tables 2, 3**). To avoid simultaneously affecting all Gsp1 functions and to create viable mutant strains (as Gsp1 is essential), we excluded mutations in the Gsp1 nucleotide binding site and the switch I and II regions (18). We confirmed by Western blot that the mutant Gsp1 protein levels were close to the endogenous wild-type levels (**Fig. S2**).

## Genetic interactions of Gsp1 mutants

To determine the functional consequences of the Gsp1 interface mutations, we performed a genetic interaction (GI) screen in *S. cerevisiae* using the epistatic mini-array profile (E-MAP) approach (19, 20). We measured growth of each GSP1 point mutant in the context of an array of single gene knockouts, resulting in a quantitative functional profile of up to 1444 GI values for each GSP1 point mutant. Similarity of genetic interaction profiles often indicates shared functions. The 55 GSP1 point mutants fell into two clusters, 23 ‘strong’ mutants with rich GI profiles containing 9-373 significant interactions (**Fig. 1d**), and 32 ‘weak’ mutants with 0-8 significant interactions (**Fig. S3a, Methods and Supplementary File 1 Fig. 2**). The strong mutants covered eleven Gsp1 sequence positions and all 16 structurally characterized Gsp1 protein interaction interfaces (**Fig. 1e**). Remarkably, twelve of the GSP1 interface point mutants had a greater number of significant GIs than an average deletion of a non-essential *S. cerevisiae* gene, and six GSP1 point mutants had more GIs than an average temperature sensitive mutant of an essential gene from a previously published large-scale *S. cerevisiae* GI map (21) (**Fig. 1f, Supplementary File 2**). The GIs of the designed Gsp1 interface mutations spanned diverse biological processes known to be linked to Gsp1, including mRNA transport, the dynein/dynactin pathway, tRNA modification by the elongator complex, and spindle assembly regulation. Furthermore, unbiased hierarchical clustering of *S. cerevisiae* genes solely by their GI profiles with the 55 GSP1 point mutants also grouped many other genes by their biological complex or pathway membership such as members of the Hog1 signaling pathway, SWR1 and Rpd3L complexes, and mitochondrial proteins (**Fig. 1d, Fig. S3b**). Taken together, the GI analysis reveals expansive functional consequences of Gsp1 interface point mutations - similar in magnitude to the effects typically observed for deleting entire genes - that illuminate many of the biological functions of GSP1.

**Fig. 2.**
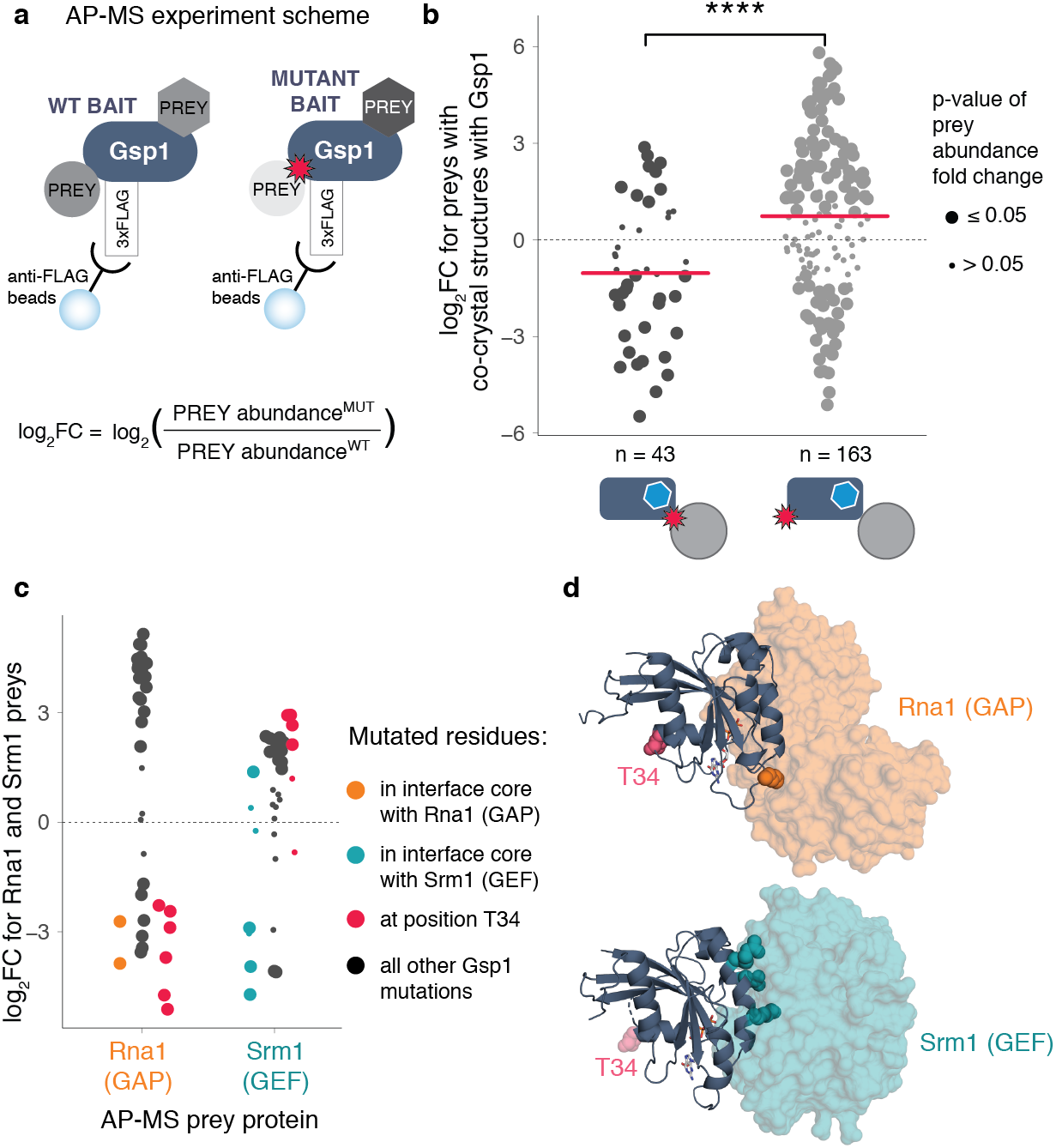
Gsp1 point mutations in the interfaces with protein partners globally rewire the physical interaction network of Gsp1, including changes in interactions with the switch regulators GEF (Srm1) and GAP (Rna1). **a,** The change in abundance of partner proteins pulled down with Gsp1 mutants is represented as log_2_ - transformed fold change (FC) between abundance pulled-down with a Gsp1 mutant versus pulled-down with wild-type Gsp1. **b,** Change in abundance of pulled-down physical interaction partners for which there are also co-complex crystal structures (Rna1, Srm1, Yrb1, Kap95, Pse1, Srp1). Left and right distributions show abundance changes for partners where the mutation is or is not in the interface core with the partner, respectively, with n denoting the number of prey abundance changes quantified in each category. Mean(log_2_ FC) values are −1 and 0.73, respectively (t-test p-value = 1.6×10-5). Point size indicates the p-value of abundance fold change, and pink bars indicate mean values. **c,** Change in abundance of pulled-down Rna1 (GAP) and Srm1 (GEF). Point size is as in b, and points are colored by interface location. **d,** Position of T34 with respect to the interfaces with Rna1 (GAP, PDB id: 1k5d) and Srm1 (GEF, PDB id: 2i1m). As the coordinates for T34 are not resolved in the 2i1m structure, on the right-hand side the pink spheres show the residue location in the aligned 1k5d structure.

In contrast to their clustering of biological processes, the GI profiles of the Gsp1 point mutants did not group based on their location in the Gsp1 partner interfaces (**Fig. 1e**). For example, strains with GSP1 mutations at residues Thr34 (T34E, T34Q) and Asp79 (D79S, D79A) have similar GI profiles (**Fig. 1d**) but these mutations are in different interaction interfaces (**Fig. 1e**) on opposite sides of the Gsp1 structure (**Fig. 1c**). This observation was unexpected and contrary to our initial expectation that Gsp1 achieves its functional specificity by interacting with different partners and, accordingly, targeting different protein interfaces should affect distinct functions of Gsp1. To analyze this finding further and quantify the functional similarities between individual GSP1 mutants across most biological processes in *S. cerevisiae*, we compared the GSP1 mutant GI profiles to profiles from 3358 *S. cerevisiae* genes (21) using Pearson correlations. In this analysis, significant positive correlations signify functional relationships (22) (**Supplementary File 3, Supplementary File 1 Table 4, Fig. S3c**). Strikingly, GI profiles of GSP1 mutants and of Gsp1 physical interaction partners were on average no more similar to each other in instances where the Gsp1 mutation was located in the partner interface than when the mutation was not (**Fig. 1g, Fig. S3d**). These results suggest that the rich functional profiles of GSP1 mutants cannot simply be explained by considering only the interface or partner interaction targeted by the point mutation.

## Physical interactions of Gsp1 mutants

To investigate further why the GI profiles of Gsp1 mutations did not group based on targeted specific physical interactions of Gsp1, we sought to determine how the physical protein interaction network of Gsp1 changes in response to the interface point mutations when all binding partners are present at their endogenous levels. We tagged wild-type Gsp1 and 28 mutants covering all interface residues shown in **Fig. 1e** with an amino-or carboxy-terminal 3xFLAG tag and quantified the abundance of 316 high-confidence ‘prey’ partner proteins in complex with Gsp1 by AP-MS (**Fig. 2a, Fig. S4, Supplementary File 4**). We refer to the prey partner protein abundance in the pulled-down Gsp1 complexes simply as “abundance” below. We quantified the abundance changes of six of the 16 Gsp1 binding partners for which we had structural information and that were robustly observable in the AP-MS data for both Gsp1 wild type and mutants: the two core regulators Rna1 (GAP) and Srm1 (GEF), as well as four effectors Yrb1, Kap95, Pse1 and Srp1 (**Fig. S5**). As expected, the abundance of the prey partner was decreased on average (although not always) when the Gsp1 mutation was in the interface core with the prey partner (**Fig. 2b**, left distribution, **Data Fig. S5a**). However, instead of expected minimal effects, we also found notable changes in prey abundance in cases where the mutation was not directly in the interface (**Fig. 2b** right distribution, **Fig. S5b**). In particular, a wide spread of abundance changes was apparent for the two main GTPase regulators, GAP (Rna1) and GEF (Srm1), even for the mutations that are outside either of the interfaces (**Fig. 2c, Fig. S5, Supplementary File 1 Table 5**). For example, mutations at the position 34 of Gsp1, which is in the core of the interface with Yrb1, increase the abundance of pulled-down GEF, and decrease the abundance of pulled-down GAP, even though the residue is outside either of the interfaces (**Fig. 2d**). In summary, the AP-MS experiments show that the point mutations, in addition to affecting the targeted interactions, also introduce extensive changes to the physical interaction network of Gsp1 that cannot simply be explained by the interface location of the mutations.

## Effect of Gsp1 point mutants on the molecular function of the switch

The AP-MS experiments showed that most Gsp1 interface mutations significantly altered physical interactions with the two principal GTPase regulators, GAP and GEF. This observation prompted the question whether the mutations, rather than acting indirectly in the cellular context (i.e. by altering the competition between physical interaction partners in the cell) affected molecular function of the switch directly. To assess the molecular effects of mutation on the switch function, we recombinantly expressed and purified wild-type and 24 Gsp1 mutants and measured their effects on GAP-mediated GTP hydrolysis and GEF-mediated nucleotide exchange in vitro (**Fig. 3a, b, Fig. S6, Supplementary File 1 Figures 3, 4,** and **Tables 6, 7**). Of the 24 Gsp1 point mutants, 17 (all of which had strong GI profiles except K132H) showed 3-to >200-fold change in k_cat_/K_m_ on either or both of the GAP- or GEF-mediated reactions (**Fig. S6e**). In particular, mutations that are not in the interface core with the GAP both increased (3-fold, R108G mutant) and decreased (3 to 10-fold, T34E/Q/A/G, R78K, D79S/A, R108I, and R112S mutants) the catalytic efficiency k_cat_/K_m_ of GAP-mediated GTP hydrolysis, compared to wild-type Gsp1 (**Fig. 3a**). As expected, mutations in the interface core with the GEF (K101, and R108) decreased the catalytic efficiency of GEF-mediated nucleotide exchange >40-fold. However, other mutations not in the GEF interface core (R78K, R112S, Y157A) also decreased the efficiency notably (3- to 10-fold, **Fig. 3b**). These results show that Gsp1 interface mutations are capable of modulating the GTPase cycle by directly affecting GTP hydrolysis and nucleotide exchange catalyzed by the two core switch regulators, GAP and GEF. Moreover, since nine out of the 17 mutations with larger than 3-fold effects are located outside of the interface cores with either the GAP or the GEF as well as outside the known switch regions, our data suggest considerable, previously unappreciated, allostery in the GTPase switch.

**Fig. 3.**
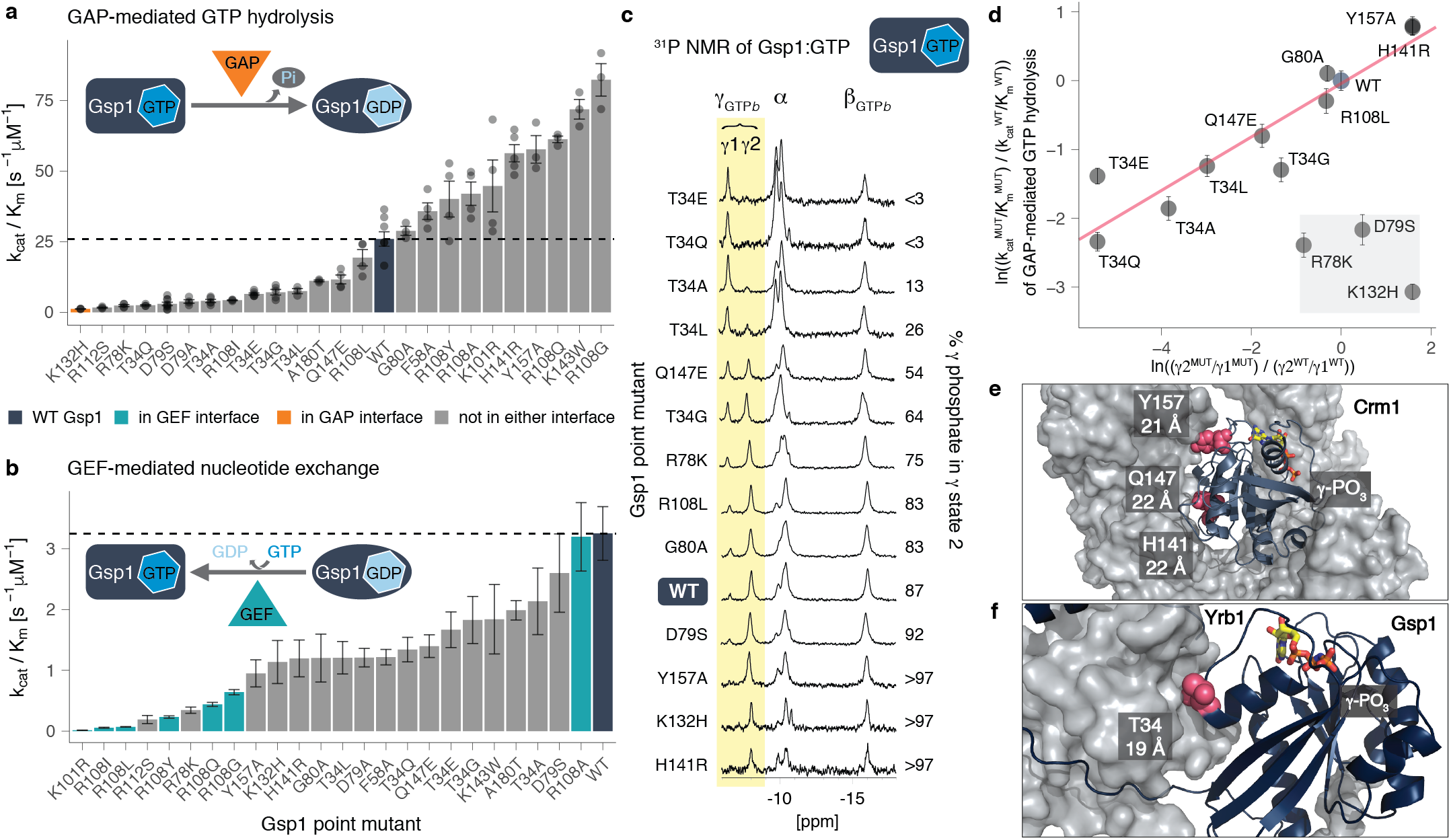
Point mutations in Gsp1 interfaces allosterically modulate GTPase cycle parameters by tuning active site conformational distributions. **a,** Catalytic efficiency (k_cat_/K_m_) of GAP-mediated GTP hydrolysis of Gsp1 mutants. Each individual point represents the ratio of k_cat_ and K_m_ from an individual GAP-mediated GTP hydrolysis experiment fit to an integrated Michaelis-Menten equation (see **Methods**). Error bars represent the value plus/minus standard error of the mean of estimated catalytic efficiency k_cat_/K_m_ from n >= 3 replicates. Dotted line indicates wild-type efficiency. **b,** Catalytic efficiency (k_cat_/K_m_) of GEF-mediated nucleotide exchange of Gsp1 mutants. Error bars represent the value plus/minus the standard error of the mean of the Michaelis-Menten fit to data from n >= 17 measurements at different substrate concentrations. **c,** 31 P NMR of GTP bound Gsp1 point mutants. NMR peak heights are normalized to the β peak of the bound GTP (βGTPb). The two peaks of the γ phosphate of bound GTP are highlighted in yellow. The peak at approximately −7 ppm is defined as γ1 and the peak at approximately −8 ppm is defined as γ2. The percent of γ phosphate in γ2 is defined as a ratio of areas under the curve between the γ2 and the sum of the γ1 and γ2 peaks. **d,** Natural log-transformed ratios MUT/WT of the exchange equilibrium constants K_ex_ = population in γ2 / population in γ1 (assuming a detection limit of 3% for the γ peak estimation by ^31^ P NMR) plotted against the natural log-transformed ratios MUT/WT of the relative catalytic efficiency (k_cat_/K_m_) of GAP-mediated GTP hydrolysis. Error bars represent the mean plus/minus standard error of the mean across at least three replicates of individual GAP-mediated GTP hydrolysis measurements. Pink line shows the least-squares linear fit, excluding the GAP interface mutation K132H and the two mutations adjacent to the switch II, R78K and D79S (gray box). **e,** Residues Y157, H141, and Q147 (pink spheres) are in the interface with Crm1 (gray surface, PDB ID: 3m1i). Gsp1 is represented as a navy cartoon and the GTP nucleotide is in yellow stick representation. **f,** Residue T34 (pink spheres) is in the interface with Yrb1 (gray surface, PDB ID: 1k5d). Distances from the γ phosphate of GTP to the residue α-carbon are indicated below the residue numbers in **e** and **f**.

**Fig. 4.**
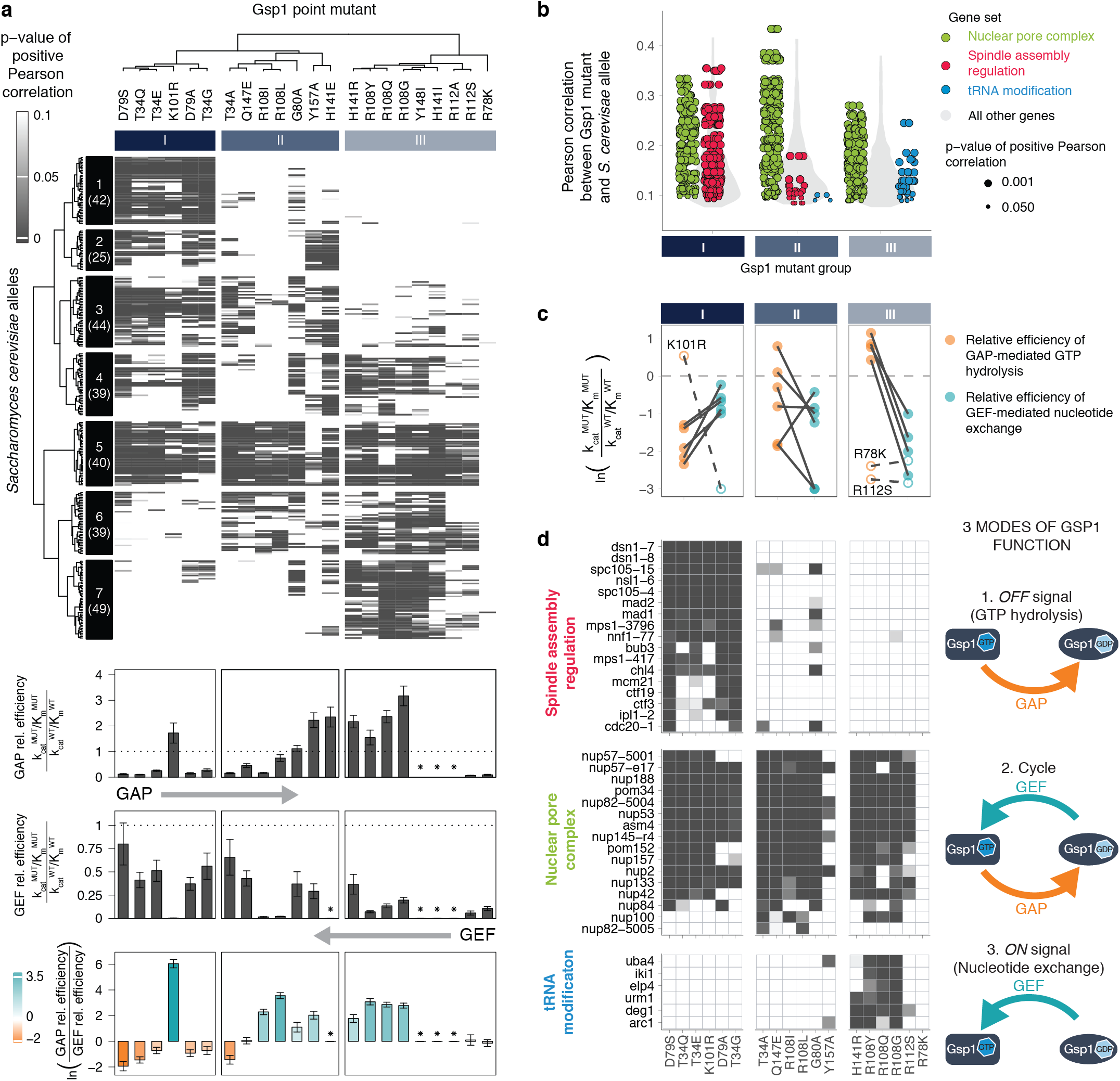
Cellular effects of interface mutations group by their effect on GTPase cycle kinetics. **a,** Clustering of 278 *S. cerevisiae* alleles and 22 strong Gsp1 point mutants by the p-value of Pearson correlations of their GI profiles compared to the relative efficiencies of GAP-mediated GTP hydrolysis and GEF-mediated nucleotide exchange as indicated (asterisks indicate not measured). The p-value is a false discovery rate adjusted one-sided (positive) p-value of the Pearson correlations (represented as gray scale). The number of genes in each of the seven clusters is given in parentheses. **b,** Distributions of Pearson correlations from **a**, separated by groups of Gsp1 point mutants identified by hierarchical clustering (see column dendrogram in **a**). Individual correlations with *S. cerevisiae* genes in three specific gene sets are shown as green, red, or blue points, while the distributions of correlations with all other genes are shown in a gray violin plot. Point size indicates the false discovery rate adjusted one-sided (positive) p-value of the Pearson correlation. Only significant correlations (p-value < 0.05) are included. **c,** Kinetic characteristics of Gsp1 mutant groups I to III. Outliers (empty circles and dashed lines) are at a site of potential posttranslational modification (K101R) or are genetically distinct (R112S and R78K do not correlate with tRNA modification genes unlike the other group III mutants). The log ratio of relative catalytic efficiencies is capped at −3. **d,** Heatmaps of false discovery rate adjusted one-sided (positive) p-value of the Pearson correlation as in **a**, for the three representative gene sets. *S. cerevisiae* genes for each gene set are clustered by p-value. The GTPase cycle schemes next to the heatmaps represent the three modes of Gsp1 function. In **c** and **d** only Gsp1 mutants with kinetics data are shown, ordered and grouped as in **a**.

To probe the mechanism of these allosteric effects, we examined the impact of Gsp1 point mutations on the conformational distribution in the active site of GTP-bound Gsp1 using 1D ^31^P nuclear magnetic resonance (NMR) spectroscopy. Prior ^31^P NMR data on human Ran(23) and Ras (24) showed two distinct peaks for the γ-phosphate of bound GTP arising from differences in the local chemical environment of the γ-phosphate in each of two distinct conformations (termed γ1 and γ2). Our ^31^ NMR spectra of *S. cerevisiae* wild-type Gsp1:GTP showed two distinct peaks for the γ-phosphate of bound GTP with 87% of wild-type Gsp1:GTP in the γ2 state conformation (**Fig. 3c, Fig. S7a**). Strikingly, the relative populations of the γ1 and γ2 states were modulated by our Gsp1 interface mutations, and ranged from close to 0% in the γ2 state for T34E and T34Q, to close to 100% for H141R, Y157A, and K132H (**Fig. 3c**). Furthermore, we observed a linear relationship between the effect of the mutation on the equilibrium between the two γ1 and γ2 conformations (plotted as the natural log-transformed ratio of the equilibrium constant) and the natural log-transformed ratio of the relative catalytic efficiencies of GAP-mediated GTP hydrolysis (**Fig. 3d**) and intrinsic GTP hydrolysis (**Supplementary File 1 Table 8, Fig. S7b and c**). This relationship suggests that the γ2 state represents the active site conformation of Gsp1:GTP competent for GTP hydrolysis. Exceptions to the linear relationship are the K132H mutation, which is in the core of the GAP interface and is hence expected to directly affect the interaction with the GAP, and the D79S and R78K mutations, which are on the edge of the GTPase switch II region (from residues 69 to 77) and could lead to different perturbations of the nucleotide binding site geometry. Remarkably, the mutated residues that tune the population of the γ2 state (T34, H141, Q147, and Y157) are all distal, affecting the chemical environment of the Gsp1-bound GTP γ phosphate from at least 18 Å away (**Fig. 3e, f**). Therefore, our data support an allosteric mechanism where distal mutations at different surface interaction sites of Gsp1 modulate the GTPase switch and in particular the efficiency of GTP hydrolysis, although further studies are required to characterize the conformational changes underlying these effects. Interestingly, neither of the allosteric sites in Gsp1 overlap with the allosteric inhibitor pockets successfully targeted by small molecule inhibitors in Ras(25–27) (**Fig. S7d**). Our *in vitro* analysis showed that most Gsp1 interface mutations affect GEF-catalyzed nucleotide exchange and GAP-catalyzed GTP hydrolysis differentially (**Fig. 3a,b** and **Fig. S6e**). To determine to what extent these effects explain the observed changes in the physical interaction network of Gsp1 (**Fig. 2**), we compared *in vitro* kinetic and our AP-MS data (**Fig. S8**). We found that all Gsp1 mutants that affected the efficiency (k_cat_/K_m_) of GEF-catalyzed nucleotide exchange more than the efficiency of GAP-catalyzed GTP hydrolysis (points above the diagonal in **Fig. S8a**) showed a larger decrease in the abundance of pulled-down GEF compared to pulled-down GAP (teal points), while the opposite was the case for the mutants below the diagonal (orange points in **Fig. S8a**). Similar but weaker relationships were observed for other prey proteins (**Fig. S8d-f**). We conclude that Gsp1 interface mutations allosterically perturb the GTPase cycle, and that the direction of the cycle perturbation is a good predictor of altered physical interactions with the two main cycle regulators, even in the context of many other potentially competing partner proteins.

## Encoding of Gsp1 multi-specificity

We next asked whether the allosteric effects of the mutations on the balance of the GTPase cycle, rather than the interface in which a mutation is made, could better explain the functional effects observed in the cellular GI profiles. We clustered the GI profiles of the Gsp1 mutants based on correlation with the GI profiles of 3358 *S. cerevisiae* alleles(21). We then compared clustering of these GI profile correlations (using the 278 alleles with significant correlations to Gsp1 mutants, **F ig. 4a**) with the biophysical effects of the mutations on the efficiencies of GAP-catalyzed GTP hydrolysis and GEF-catalyzed nucleotide exchange. Remarkably, the Gsp1 mutant GI profile clustering mirrored an approximate ordering by the *in vitro* mutant effects on the GTPase cycle: relative GAP efficiency systematically increased with increasing column number and relative GEF efficiency decreased (**Fig. 4 a**). A clear outlier of this ordering is the K101R mutant, which primarily affects GEF-mediated nucleotide exchange *in vitro* but, by GI profiles, groups with mutations affecting the efficiency of GTP hydrolysis. The lysine at this position was found to be acetylated in both *S. cerevisiae*(28) and human (29) cells. The acetylation at this position in human Ran was shown to reduce the efficiency of nucleotide release from the Ran:GDP:GEF complex (30). We hypothesize that while our K101R mutation affected the interaction with the GEF, it also likely broke a critical mechanism by which the cell reduces GEF activity, phenocopying the mutants with reduced GTP hydrolysis activity. Finally, we asked whether our biophysical measurements of how the different Gsp1 mutants perturb the GTPase cycle regulation could provide insight into Gsp1’s functional multi-specificity, i.e. its ability to distinctly regulate multiple biological processes. Clustering the *S. cerevisiae* genes and Gsp1 mutants based on correlations of their respective GI profiles revealed that genes fall into one of three categories (**Fig. 4a**): (i) genes in cluster 1, but also cluster 2, that correlate primarily with mutants more perturbed in the GTP hydrolysis side of the cycle (orange bars in the bottom barplot in **Fig. 4a**, showing the ratio of relative GAP and GEF efficiency), (ii) genes in cluster 7 that correlate primarily with mutants more perturbed in the nucleotide exchange side of the cycle (teal bars), and (iii) genes that correlate strongly with all or most of the Gsp1 point mutants irrespective of the direction of GTPase cycle perturbation (most strikingly genes in cluster 5, but also clusters 3, 4, and 6). Importantly, clustering of *S. cerevisiae* genes by the correlations of their GI profiles with GI profiles of Gsp1 mutants distinguishes between biological processes, where genes with shared biological functions (gene sets, **Supplementary File 5**) all predominantly fall into one of the three categories defined above. For example, genes involved in spindle assembly regulation have significant GI profile correlations primarily with Gsp1 mutant group I (red points in **Fig. 4b**), genes involved in tRNA modification primarily with Gsp1 mutant group III (blue points in **Fig. 4b**), and genes important for nucleocytoplasmic transport with Gsp1 mutants from all three groups (green points in **Fig. 4b**). Hence, while the effects of the Gsp1 mutations cannot be explained simply by interface location (**Fig. 1g, Fig. 2b**), the three groups of Gsp1 mutants by GI profiles (**Fig. 4a**) show distinct kinetic characteristics: Groups I and III have decreased efficiencies of GTP hydrolysis or nucleotide exchange, respectively, and group II shows intermediate behavior (**Fig. 4c**). Taken together, our analysis suggests that distinct cellular processes regulated by Gsp1, such as spindle assembly regulation, tRNA modification, and nuclear transport (**Fig. 4b, d**), as well as 5’ mRNA capping, transcriptional regulation, cytoplasm-to-vacuole targeting, and actin, tubulin and cell polarity (**Fig. S9**) are differentially sensitive to perturbations of GTPase cycle kinetics (**Fig 4d**, right).

## Discussion

Only five years after the discovery of the small GTPase Ran, Rush et al.(31) proposed that Ran must act by two different mechanisms: one in which the cycling of the GTPase is most important (‘Rab paradigm’), and the other in which the amount of “active” Ran:GTP is most important (‘Ras paradigm’). Our findings lead to a model where Ran/Gsp1 acts by three different paradigms that are defined by the sensitivity of different biological processes to perturbations of different characteristics of the Gsp1 GTPase cycle, i.e. the ability to (i) cycle, (ii) turn off by hydrolyzing to Gsp1:GDP, and (iii) turn on by producing Gsp1:GTP (**Fig. 4d, Fig. S9**). Other effects such as direct perturbations of interactions, binding partner competition, and small changes in expression of Gsp1 or its partners undoubtedly also play a role in modulating the functional effects of our Gsp1 mutations. Nevertheless, our model explains to a remarkable degree how a single molecular switch motif can differentially control subsets of biological processes by using one of the three functional modes. Furthermore, our model is consistent with previous studies of canonical GTPase mutations in Ran, where mutants defective in GTP hydrolysis or nucleotide exchange were exogenously expressed or injected into cells. For example, the *S. cerevisiae* Gsp1 G21V mutant defective in GTP hydrolysis abrogated Mad1 turnover during spindle assembly checkpoint regulation (32) consistent with the functional effects of our Gsp1 mutants affecting hydrolysis (**Fig. 4c-d**). The Xenopus Ran T24N mutant with impaired nucleotide exchange disrupted microtubule assembly (33), consistent with our observation that the effects of ON switch mutants correlate with actin, tubulin, and polarity related genes (**Fig. S9c**). Due to the widespread allostery observed in Gsp1, precisely designing novel mutations to perturb individual Gsp1 functions remains a significant challenge, but our work provides a set of viable mutants with a range of effects on the GTPase cycle that can be used to further address open questions on how switch cycle dynamics differentially impact the cellular functions of Gsp1. The discovery of several allosteric sites (positions 34, 141, 147, and 157) in the model molecular switch Gsp1 both explains the widespread functional consequences we observe for single amino acid point mutations at interaction surfaces of Gsp1 and has important implications for revising our understanding of GTPase switch regulation. We show that mutations in distal interfaces allosterically modulate the switch cycle. This finding demonstrates thermodynamic coupling between interfaces and the classical switch region in the active site and thereby suggests that partners binding to distal sites also regulate the switch by affecting conformational equilibria at the active site. This hypothesis is supported by evidence that the Yrb1 homolog RanBP1 modulates GAP activity (23, 34, 35). Our data provide a mechanistic explanation, where mutations at allosteric sites, including T34 in the Yrb1 binding interface, tune the population of Gsp1 in a hydrolytically-primed conformation. Since the overall switch mechanism is conserved across the small GTPase fold, we propose that thermodynamic coupling between distal interfaces and functional conformational changes may be a more general mechanism to regulate other GTPase switches, and may aid in the development of allosteric inhibitors. Our observation of widespread functional effects of point mutations inducing relatively small perturbations in the GTPase switch kinetics is reminiscent of the zero-order ultrasensitivity achievable in biological motifs with opposing regulators (6). While switch-like ultrasensitivity is typically described for systems controlled by covalent modifications (such as phosphorylation), our results, as well as the observations that cellular levels of small GTPase regulators require tight control (36, 37), corroborate a model of ultra-sensitivity for GTPase conformational switches (38). While we investigated the changes to the GTPase cycle caused by mutations, similar effects on regulation could be exerted by partner binding or posttranslational modification. Finally, deriving a model that explains the cellular multi-specificity of GTPases by differential sensitivity of biological processes to distinct parameters of the switch cycle was enabled by a quantitative analysis that integrated functional genomics, proteomics, and biophysics. Given the prevalence of biological two-state switch motifs controlled by opposing regulators (kinase/phosphatase, acetylase/deacetylase) (39), we envision this approach to be fruitful for other studies of cellular regulation and to be extended to mammalian systems using CRISPR-based approaches to yield mechanistic insights into the drastic consequences of disease mutations targeting central molecular switches.

## Methods

### Point mutations in genomic Gsp1 sequence

We identified all residues in Gsp1 that comprised the interfaces with Gsp1 binding partners for which co-complex crystal structures with Gsp1 were available (**Supplementary File 1 Fig. 1, Fig. S1a, Supplementary File 1 Table 1**). Residues comprising the interface core, the surface exposed rim around the core, and more buried support residues were defined based on per-residue relative solvent accessible surface area (rASA), as previously described(5). rASA is compared to the empirical maximum solvent accessible surface area for each of the 20 amino acids(40). rASA values were calculated for the Gsp1 monomer (rASAmonomer) and for the complex (rASAcomplex) using the bio3d R package(41). The three types of interface residues were defined as: interface core if rASAmonomer > 25%, rASAcomplex < 25% and ΔrASA (change upon complex formation) > 0; rim residues if rASAcomplex > 25% and ΔrASA > 0; and support residues if rASAmonomer < 25% and ΔrASA > 0. All custom code for interface analysis from co-complex crystal structures is provided in the associated code repository in the structure analysis folder. We avoided Gsp1 residues that are within 5 Å of the nucleotide (GDP or GTP) in any of the structures or that are within the canonical small GTPase switch regions (P-loop, switch loop I, and switch loop II). We then mutated residues that are located in interface cores (defined as residues that bury more than 25% of their surface upon complex formation, as previously defined (5), **Supplementary File 1 Table 2, Fig. S1b**) into amino acid residues with a range of properties (differing in size, charge and polarity) and attempted to make stable and viable *S. cerevisiae* strains carrying a genomic Gsp1 point mutation coupled to nourseothricin (clonNAT / nourseothricin, Werner BioAgents GmbH, CAS 96736-11-7) resistance. The list of attempted mutants is provided in **Supplementary File 1 Table 3**. The genomic construct was designed to minimally disrupt the non-coding sequences known at the time, including the 5′ UTR and 3′ UTR, as well as the putative regulatory elements in the downstream gene Sec72 (**Supplementary File 1 Fig. 5**). The *GSP1* genomic region was cloned into a pCR2.1-TOPO vector (Invitrogen) and point mutations in the *GSP1* coding sequence were introduced using the QuikChange™ Site-Directed Mutagenesis (Stratagene, La Jolla) protocol. *S. cerevisiae* strains containing mutant *GSP1* genes were regularly confirmed by sequencing the Gsp1 genomic region.

### *S. cerevisiae* genetics and genetic interaction mapping

#### *S. cerevisiae* transformation

To generate MAT:α strains with Gsp1 point mutations the entire cassette was amplified by PCR using *S. cerevisiae* transformation forward and reverse primers, and *S. cerevisiae* was transformed into the starting SGA MAT:α his3D1; leu2D0; ura3D0; LYS2þ; can1::STE2pr-SpHIS5 (SpHIS5 is the *S. pombe* HIS5 gene); lyp1D::STE3pr-LEU2 strain from (42) as described below. Primers for amplifying the *GSP1* genomic region: *S. cerevisiae* Transformation FWD: GTATGATCAACTTTTCCT-CACCTTTTAAGTTTGTTTCG *S. cerevisiae* Transformation REV: GATTGGAGAAACCAACCCAAATTTTACACCACAA

DNA competent *S. cerevisiae* cells were made using a LiAc protocol. The final transformation mixture contained 10 mM LiAc (Lithium acetate dihydrate, 98%, extra pure, ACROS Organics™, CAS 6108-17-4), 50 µg ssDNA (UltraPure™ Salmon Sperm DNA Solution, Invitrogen, 15632011), 30% sterile-filtered PEG 8000 (Poly(ethylene glycol), BioUltra, 8,000, Sigma-Aldrich, 89510-250G-F). A *S. cerevisiae* pellet of approximately 25 μl was mixed with 15 μl of linear DNA PCR product and 240 μl of the transformation mixture, and heat shocked at 42°C for 40 minutes. Transformed cells were grown on YPD (20 g Bacto™ Peptone (CAT # 211820, BD Diagnostic Systems), 10 g Bacto™ Yeast Extract (CAT # 212720 BD), and 20 g Dextrose (CAT # D16-3, Fisher Chemicals) per 1-liter medium) + clonNAT plates and incubated at 30°C for 3 to 6 days. Many colonies that appeared after 24-48 hours carried the clonNAT cassette but not the *GSP1* point mutation, or the 3×FLAG tag. Cells were therefore sparsely plated and plates were incubated for a longer period of time after which colonies of different sizes were picked and the mutant strains were confirmed by sequencing.

### Epistatic mini-array profiling (E-MAP) of Gsp1 point mutants

Genetic interactions of all viable *GSP1* point mutant (PM-GSP1-clonNAT) strains were identified by epistatic miniarray profile (E-MAP) screens (20, 4 2) using a previously constructed array library of 1,536 KAN-marked (kanamycin) mutant strains assembled from the *S. cerevisiae* deletion collection (43) and the DAmP (decreased abundance by mRNA perturbation) strain collection (44), covering genes involved in a wide variety of cellular processes (19, 45). The E-MAP screen was conducted as previously described in Collins et al. (42), using the HT Colony Grid Analyzer Java program (22) and E-MAP toolbox for MATLAB to extract colony sizes of double mutant strains and a statistical scoring scheme to compute genetic interaction scores. Genetic interaction scores represent the average of 3-5 independent replicate screens. Reproducibility was assessed as previously described (22) by comparing individual scores to the average score for each mutant:gene pair, with the two values showing strong correlation across the dataset (Pearson correlation coefficient = 0.83, **Supplementary File 1 Fig. 6**).

### Hierarchical clustering of E-MAP genetic interaction data

All E-MAP library DAmP strains as well as library strains showing poor reproducibility were discarded, leaving 1444 out of the original 1536 library genes. Averaged S-scores of genetic interactions between wild type and point mutant Gsp1 and the 1444 *S. cerevisiae* genes are provided in **Supplementary File 2**. Hierarchical clustering on the genetic interaction profiles was performed using the average linkage method and the pairwise Pearson correlation coefficient as a distance metric. To identify clusters of functionally related library genes, the hierarchical clustering tree was cut to produce 1200 clusters, resulting in 43 clusters with 3 or more members. Biological function descriptions for genes in these clusters were extracted from the Saccharomyces Genome Database (SGD) (46). Clusters of genes representing common functions (complexes, pathways or biological functions) were selected by manual inspection and represented in the main text **Fig. 1d** and **Fig. S3b**. All of the custom code for E-MAP analysis is provided in the E-MAP analysis GitHub repository. Clustered heatmaps were produced using the ComplexHeatmap package (47).

### Scaling of published genetic interaction data to the E-MAP format

To enable comparison of *GSP1* point mutant genetic interaction profiles to profiles of other *S. cerevisiae* genes, published Synthetic Gene Array (SGA) genetic interaction data (21) from CellMap.org (48) were scaled to the E-MAP format using a published non-linear scaling method (49). Briefly, 75,314 genetic interaction pairs present in both the SGA and a previously described E-MAP dataset used to study chromatin biology (50) were ordered by genetic interaction score and partitioned into 500 equally sized bins separately for each dataset. Bin size (150 pairs per bin) was chosen to provide enough bins for fitting the scaling spline (described below) while still maintaining a large number of pairs per bin such that the mean could be used as a high confidence estimate of the score values in each bin. Scaling factors were computed that scaled the mean of each SGA bin to match the mean of the corresponding E-MAP bin. A non-linear univariate spline was fit through the scaling factors, providing a scaling function that was subsequently applied to each SGA score. The distribution of scores of shared interactions between the scaled SGA and the E-MAP chromatin library was similar to that between replicates in the E-MAP chromatin library, matching what was seen in the previously published scaling of SGA data to E-MAP format (49) (**Supplementary File 1 Fig. 7**). The SGA genetic interaction scores are taken from CellMap.org (48). The scaling code is provided in SGA scaling GitHub repository.

### Significance of genetic interactions

The S-score metric used in scoring genetic interactions measured by the E-MAP method has been previously characterized in terms of confidence that any given averaged S-score represents a significant interaction (22). We fit a spline to data points from Fig. 4c from Collins *et al* (22), allowing us to provide an approximate confidence estimate for each of our measured GSP1 and scaled *S. cerevisiae* SGA genetic interaction scores. The SGA dataset (21) is accompanied by p-values as well as its own recommendations for a threshold at which individual interactions are considered significant. We plotted the SGA score scaled to E-MAP format vs. the associated p-value (negative log-transformed, **Supplementary File 1 Fig. 2a**) and found the distribution to have a similar shape to the confidence function for S-scores (**Supplementary File 1 Fig. 2b**). For example, a 95% confidence threshold is associated with E-MAP S-scores less than −4 or greater than 5, while the median p-value of scaled SGA scores is less than 0.05 for scores less than −5 or greater than 3. We ultimately elected to use a significance cutoff of absolute S-score greater than 3. This threshold corresponds to an estimated confidence value of 0.83 for S-scores less than −3 and 0.65 for S-scores greater than 3. We compared these values to the intermediate significance threshold recommended for the SGA data from Ref. (21), which was p-value < 0.05 and absolute SGA score > 0.08. After scaling to E-MAP format, this threshold corresponds to scaled S-scores less than −2.97 or greater than 2.25, below our chosen threshold of −3 and 3.

### GI profile correlation measurements

Of the 1444 library genes in the *GSP1* point mutant GI profile map, 1129 were present in the SGA dataset from Ref. (21). Pairwise Pearson correlation coefficients were computed between all *GSP1* point mutants and SGA gene profiles, and all profiles trimmed to include only genetic interaction measurements with the 1129 shared library genes. Due to the relative sparsity of genetic interaction profiles, pairwise comparisons are dominated by high numbers of non-significant interactions. Accordingly, we did not consider correlations with *GSP1* point mutants or SGA gene profiles that did not have significant genetic interactions (absolute scaled S-score greater than 3, see above) with at least 10 of the 1129 library genes. This requirement removed all weak Gsp1 point mutants and one strong mutant (R108A) from the correlation analysis (as they had at most nine genetic interactions with absolute score greater than 3), leaving 22 strong mutants and 3383 *S. cerevisiae* SGA genes to be included in the correlation analysis. All Pearson correlations and their p-values between Gsp1 mutants and *S. cerevisiae* genes, including all correlations that did not pass our significance filtering procedures, are provided in **Supplementary File 3**. The subset of Pearson correlations between Gsp1 point mutants and Gsp1 partners with available co-complex X-ray crystal structures, used to make the point plots in **Figs. 1g** and **7c, d**, are also available in **Supplementary File 1 Table 4**. Statistical significance of correlations was computed using both two-sided and one-sided (positive) t-tests adjusted for multiple hypothesis testing using both the Bonferroni method and the FDR method, which controls the false discovery rate (51). All p-values reported in the text and figures are one-sided (positive) and corrected by the FDR method, unless otherwise stated. Custom code for genetic interaction profile correlation calculations and filtering is provided in the accompanying E-MAP correlation GitHub repository.

Significance testing was used to filter out *S. cerevisiae* gene SGA profiles that did not show a significant correlation (one-sided positive, Bonferroni-adjusted) with at least two *GSP1* point mutants profile. In total, 278 *S. cerevisiae* genes from the SGA had a significant profile correlation (one-sided positive, Bonferroni-adjusted) with at least two *GSP1* point mutants and were therefore included in the correlation analysis shown in **Fig. 4b** and **Fig. S8a**. After this filtering step, the one-sided p-values were used to populate a matrix of 22 mutants vs. 278 genes, and hierarchical clustering was performed using Ward’s method. Pearson correlation between correlation vectors was used as a distance metric for the mutant (row) clustering, while Euclidean distance was selected for the gene (column) clustering, due to the column vectors being relatively short (22 mutants per column vs. 278 genes per row) and thus sensitive to outliers when clustered using Pearson correlations as the distance metric.

For the gene set analysis we decreased the stringency of inclusion of *S. cerevisiae* SGA genes to include all genes with a significant profile correlation (one-sided positive, Bonferroni-adjusted) with one or more Gsp1 mutants, which added another 201 genes, resulting in 479 genes. We made the gene sets larger to increase our confidence in connecting the patterns of correlations between *S. cerevisiae* genes and Gsp1 mutants, and GTPase cycle parameters represented in **Fig. 4c** and **Fig. S9**. Manually curated gene sets of *S. cerevisiae* genes with significant correlations with Gsp1 mutants are provided in **Supplementary File 5**.

### Protein expression levels by Western Blot

*S. cerevisiae* strains were grown at 30°C in YPD medium (20 g Bacto™ Peptone (CAT # 211820, BD Diagnostic Systems), 10 g Bacto™ Yeast Extract (CAT # 212720 BD), and 20 g Dextrose (CAT # D16-3, Fisher Chemicals) per 1 L medium) for 1.5 - 2 hours until OD_600_ reached 0.3. Cell culture aliquots of 1 ml were centrifuged for 3 minutes at 21,000 × g and resuspended in 30 μl of phosphate buffered saline (137 mM NaCl, 2.7 mM KCl, 10 mM Na_2_HPO_4_, 1.8 mM KH_2_PO_4_, pH = 7.4) and 10 μl of SDS-PAGE Sample Buffer (CAT # 161-0747, BioRad), to a final SDS concentration of 1%, and 2 mM beta-mercaptoethanol. Lysates were run (3 μl for most, and 6 μl for slow growing mutants with lower OD600) on Stain-Free gels (4-20%, CAT # 4568096, Bio-Rad, Tris/Glycine SDS Buffer (CAT # 161-0732, BioRad)). After electrophoresis, the gel was scanned for total protein quantification and the proteins were subsequently transferred to an Immobilon-FL PVDF membrane (CAT # IPF00010, EMD Millipore). The membrane was probed with Rabbit anti-RAN (CAT # PA 1-5783, ThermoFisher Scientific) primary, and Goat anti-Rabbit-IgG(H+L)-HRP (CAT # 31460, Thermo Fisher) secondary antibodies. The membrane was developed using Super Signal West Femto substrate (CAT # 34096, Thermo Fisher), and scanned and analyzed with Image Lab software on a ChemiDoc MP (BioRad). Each blot had at least one wild-type (WT-GSP1-clonNAT) and at least one MAT:α strain control. The total protein levels (TP^MUT^) for each Gsp1 point mutant lane were then normalized to the wild-type (WT-GSP1-clonNAT) lane of the corresponding blot (TP^WT^), providing an adjustment value to account for differences in loading between lanes

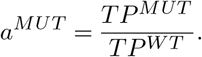

To compute the relative expression of a Gsp1 point mutant, the density (D^MUT^) of the Western blot bands corresponding to the Gsp1 point mutant was divided by the total protein adjustment and finally normalized against the same value for the wild type Gsp1, i.e.

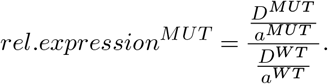

Note that for blots with a single WT lane, a^WT^ = 1. For blots with more than one WT lane included, a^WT^ was computed for each WT lane by normalizing to the average TP across all WT lanes, and the average adjusted WT density (D^WT^/a^WT^) across all WT lanes was used for computing the relative expression of point mutants. Example Western blots are provided in **Supplementary File 1 Fig. 8**, and the final protein expression level data for all the mutants are shown in **Fig. S2**.

### Identifying Gsp1 complexes in *S. cerevisiae* with affinity purification mass spectrometry (AP-MS)

#### *S. cerevisiae* cell lysate preparation

When choosing mutants for AP-MS we sought to cover all Gsp1 sequence positions where mutations had strong GI profiles (**Fig. 1e**), as well as several ‘weak’ mutants. We observed that tagging the endogenous Gsp1 with either an amino-terminal or a carboxy-terminal FLAG tag affects the *S. cerevisiae* growth in culture. We therefore attempted to make each of the mutants intended for AP-MS experiments with both tags, and where both tags were viable, we obtained the AP-MS data for both. We could not make a FLAG-tagged R108Q mutant for AP-MS. *S. cerevisiae* strains for AP-MS were grown in YAPD medium (120 mg adenine hemisulfate salt (CAT # A9126, SIGMA), 10 g Bacto yeast extract (CAT # BD 212720), 20 g Bacto peptone (CAT # BD 211820), 20 g dextrose (D-glucose D16-3 Fisher Chemicals) per 1 L of medium). Each strain was grown at 30°C for 12 to 24 h to OD_600_ of 1-1.5. The cells were harvested by centrifugation at 3000 RCF for 3 minutes and the pellet was washed in 50 ml of ice-cold ddH_2_O, followed by a wash in 50 ml of 2x lysis buffer (200 mM HEPES pH 7.5, 200 mM KCl, 2 mM MgCl_2_, 30 μM GTP (Guanosine 5′-triphosphate sodium salt hydrate, CAT # G8877, Sigma-Aldrich), 1 mM Dithiothreitol (Promega V3151), 0.1% IGEPAL CA-630 (CAT # I8896, Sigma-Aldrich), and 10% glycerol). Each pellet of approximately 500 μl was then resuspended in 500 μl of 2X lysis buffer supplemented with protease inhibitors without EDTA (cOmplete, Mini, EDTA-free Protease Inhibitor Cocktail, CAT # 11836170001, Roche) and dripped through a syringe into liquid nitrogen. The frozen *S. cerevisiae* cell pellets were lysed in liquid nitrogen with a SPEX™ SamplePrep 6870 Freezer/Mill™.

#### FLAG immunoprecipitation

FLAG immunoprecipitations were performed as previously described (52, 53). Details are as follows. For FLAG immunoprecipitations, frozen samples were initially kept at room temperature for 5 minutes and then placed on ice or at 4°C in all subsequent steps, unless indicated otherwise. Following the addition of 1.5 – 3.0 ml Suspension Buffer (0.1 M HEPES pH 7.5, 0.1 M KCl, 1 mM MgCl_2_, 15 μM GTP, and 0.5 mM Dithiothreitol) supplemented with cOmplete mini EDTA-free protease and PhosSTOP phosphatase inhibitor cocktails (Roche), samples were incubated on a rotator for at least 10 minutes and then adjusted to 6 ml total volume with additional Suspension Buffer supplemented with inhibitors before centrifugation at 18,000 rpm for 10 minutes. Anti-FLAG M2 Affinity Gel beads (50 μl slurry; Sigma-Aldrich) were washed twice with 1 ml Suspension Buffer. After reserving 50 μl, the remaining supernatant and anti-FLAG M2 Affinity Gel beads were combined and incubated for >= 2 hours on a tube rotator. Beads were then collected by centrifugation at 300 rpm for 5 minutes and washed three times. For each wash step, beads were alternately suspended in 1 ml Suspension Buffer and collected by centrifugation at 2,000 rpm for 5 minutes. After removing residual wash buffer, proteins were eluted in 42 μl 0.1 mg/ml 3×FLAG peptide, 0.05% RapiGest SF Surfactant (Waters Corporation) in Suspension Buffer by gently agitating beads on a vortex mixer at room temperature for 30 minutes. Immunoprecipitated proteins (4 μl) were resolved on 4-20% Criterion Tris-HCl Precast gels (BioRad) and visualized by silver stain (Pierce Silver Stain Kit; Thermo Scientific) (**Supplementary File 1 Fig. 9**) before submitting 10 μl of each sample for mass spectrometry. At least three independent biological replicates were performed for each FLAG-tagged protein and the untagged negative control.

#### Liquid chromatography with tandem mass spectrometry (LC-MS/MS) analysis

To prepare samples for LC-MS/MS analysis, immunoprecipitated protein (10 μl) was denatured and reduced in 2 M urea, 10 mM NH_4_HCO_3_, and 2 mM Dithiothreitol for 30 minutes at 60°C with constant shaking, alkylated in the dark with 2 mM iodoacetamide for 45 minutes at room temperature and digested overnight at 37°C with 80 ng trypsin (Promega). Following digestion, peptides were acidified with formic acid and desalted using C18 Zip-Tips (Millipore) according to the manufacturer’s specifications. Samples were re-suspended in 4% formic acid, 2% acetonitrile solution, and separated by a 75-minute reversed-phase gradient over a nanoflow C18 column (Dr. Maisch). Peptides were directly injected into a Q-Exactive Plus mass spectrometer (Thermo), with all MS1 and MS2 spectra collected in the orbitrap. Raw MS data were searched against the *S. cerevisiae* proteome (SGD sequences downloaded January 13, 2015) using the default settings in MaxQuant (version 1.5.7.4), with a match-between-runs enabled (54, 55). Peptides and proteins were filtered to 1% false discovery rate in MaxQuant, and identified proteins were then subjected to protein-protein interaction scoring using SAINTexpress (56). Protein were filtered to only those representing high confidence protein-protein interactions (Bayesian false discovery rate from SAINT (SAINT BFDR) < 0.05). Protein abundance values for this filtered list were then subjected to equalized median normalization, label free quantification and statistical analysis were performed using MSstats(57), separately for data from amino- or carboxy-terminally tagged baits. Fold change in abundance of preys for 3×FLAG-tagged Gsp1 point mutants was always calculated compared to the wild type Gsp1 with the corresponding tag. All AP-MS data are available from the PRIDE repository under the PXD016338 identifier. Custom code for analysis of AP-MS data is available from the accompanying AP-MS GitHub repository. Fold change values between prey abundance between the mutant and wild type Gsp1 and the corresponding FDR adjusted p-values are provided in **Supplementary File 4**. The intersection of all prey proteins identified at least once with both the amino- or carboxy-terminal 3xFLAG tag, and their interquartile ranges (IQR) of log_2_(fold change) values across all the Gsp1 mutants, are provided in **Supplementary File 1 Table 5**. Quality of data and reproducibility between replicates was assessed based on correlations of protein abundance between replicates (**Supplementary File 1 Figs. 10, 11**).

### Biochemical and biophysical assays

#### Protein purifications

All proteins were expressed from a pET-28 a (+) vector with a N-terminal 6xHis tag in *E. coli* strain BL21 (DE3) in the presence of 50 mg/L Kanamycin. GEF (Srm1 from *S. cerevisiae*, (Uniprot P21827)) was purified as Δ1-27Srm1 and GAP (Rna1 from *S. pombe*, Uniprot P41391) as a full-length protein. ScΔ1-27Srm1 and SpRna1 were expressed in 2xYT medium (10 g NaCl, 10 g yeast extract (BD BactoTM Yeast Extract #212720), 16 g tryptone (Fisher, BP1421) per 1 L of medium) overnight at 25°C upon addition of 300 μmol/L Isopropyl-β-D-thiogalactoside (IPTG). Gsp1 variants were expressed by autoinduction for 60 hours at 20°C (58). The autoinduction medium consisted of ZY medium (10 g/L tryptone, 5 g/L yeast extract) supplemented with the following stock mixtures: 20xNPS (1M Na_2_HPO_4_, 1 M KH_2_PO_4_, and 0.5 M (NH_4_)_2_SO_4_), 50x 5052 (25% glycerol, 2.5% glucose, and 10% α-lactose monohydrate), 1000x trace metal mixture (50 mM FeCl_3_, 20 mM CaCl_2_, 10 mM each of MnCl_2_ and ZnSO_4_, and 2 mM each of CoCl_2_, CuCl_2_, NiCl_2_, Na_2_MoO_4_, Na_2_SeO_3_, and H_3_BO_3_ in 60 mM HCl). Cells were lysed in 50 mM Tris pH 7.5, 500 mM NaCl, 10 mM imidazole, and 2 mM β-mercaptoethanol using a microfluidizer from Microfluidics. For Gsp1 purifications, the lysis buffer was also supplemented with 10 mM MgCl_2_. The His-tagged proteins were purified on Ni-NTA resin (Thermo Scientific #88222) and washed into a buffer containing 50 mM Tris (pH 7.5) and 100 mM NaCl, with 5 mM MgCl_2_ for Gsp1 proteins. The N-terminal His-tag was digested at room temperature overnight using up to 12 NIH Units per mL of bovine thrombin (Sigma-Aldrich T4648-10KU). Proteins were then purified using size exclusion chromatography (HiLoad 26/600 Superdex 200 pg column from GE Healthcare), and purity was confirmed to be at least 90% by SDS polyacrylamide gel electrophoresis. Samples were concentrated on 10 kDa spin filter columns (Amicon Catalog # UFC901024) into storage buffer (50 mM Tris pH 7.5, 150 mM NaCl, 1 mM Dithiothreitol). Storage buffer for Gsp1 proteins was supplemented with 5 mM MgCl_2_.. Protein concentrations were confirmed by measuring at 10-50x dilution using a Nanodrop (ThermoScientific). The extinction coefficient at 280 nm used for nucleotide (GDP or GTP) bound Gsp1 was 37675 M-1 cm-1, as described in (59). The ratio of absorbance at 260 nm and 280 nm for purified Gsp1 bound to GDP was 0.76. Extinction coefficients for other proteins were estimated based on their primary protein sequence using the ProtParam tool. Concentrated proteins were flash-frozen and stored at −80 °C.

In our hands every attempt to purify the *S. cerevisiae* homologue of GAP (Rna1, Uniprot P11745) from *E. coli* yielded a protein that eluted in the void volume on the Sephadex 200 size exclusion column, indicating that the protein is forming soluble higher-order oligomers. We were, however, successful in purifying the *S. pombe* homologue of GAP (Rna1, Uniprot P41391) as a monomer of high purity as described above. *S. pombe* and *S. cerevisiae* Rna1 proteins have an overall 43% sequence identity and 72% sequence similarity. The X-ray crystal structure of Ran GTPase and its GAP used in our analyses is a co-complex structure of the *S. pombe* homolog of Rna1 (PDB: 15kd), human RAN, and human RANBP1 (**Supplementary File 1 Table 1**). We used the purified *S. pombe* homolog of Rna1 in all of our GTP hydrolysis kinetic experiments. Although we cannot exclude slight differences between the kinetic parameters of *S. pombe* and *S. cerevisiae* Rna1, we do not believe such differences would significantly affect our conclusions for two main reasons: First, residues in the interface with Gsp1 are highly conserved between *S. pombe* and *S. cerevisiae* GAP Rna1, suggesting that mechanism of catalysis and kinetic parameters are also likely to be similar. *S. pombe* and *S. cerevisiae* Rna1 proteins have an overall 39% sequence identity and 53% sequence similarity. Importantly, all but one interface core residues are identical in sequence between *S. cerevisiae* and *S. pombe* homologues (**Supplementary File 1 Fig. 12**). The X-ray crystal structure of Ran GTPase and its GAP used in our analyses is a co-complex structure of the *S. pombe* homolog of Rna1 (PDB: 15kd), human Ran, and human RanBP1 (*Supplementary File 1 Table 1*). Second, we rely only on the relative differences between GAP kinetic parameters of different Gsp1 mutants to group our mutants into three classes. Even in the case of differences between the absolute kinetic parameters between the *S. pombe* and *S. cerevisiae* GAP Rna1, the order of mutants is less likely to be different, and even in the case of some differences, we expect the grouping to be robust to these changes (see **Supplementary File 1 Supplementary Discussion** for more detail).

#### Circular dichroism (CD) spectroscopy of protein thermostability

Samples for CD analysis were prepared at approximately 2 μM Gsp1 in 2 mM HEPES pH 7.5, 5 mM NaCl, 200 μM MgCl_2_, and 50 μM Dithiothreitol. CD spectra were recorded at 25°C using 2 mm cuvettes (Starna, 21-Q-2) in a JASCO J-710 CD-spectrometer (Serial #9079119). The bandwidth was 2 nm, rate of scanning 20 nm/min, data pitch 0.2 nm, and response time 8 s. Each CD spectrum represents the accumulation of 5 scans. Buffer spectra were subtracted from the sample spectra using the Spectra Manager software Version 1.53.01 from JASCO Corporation. Temperature melts were performed from 25°C −95°C, monitoring at 210 nm, using a data pitch of 0.5°C and a temperature slope of 1°C per minute. As all thermal melts of wild type and mutant Gsp1 proteins were irreversible, only apparent Tm was estimated (**Supplementary File 1 Fig. 13**) and is reported in **Supplementary File 1 Table 9**.

#### GTP loading of Gsp1

Gsp1 variants for GTPase assays as well as for ^31^P NMR spectroscopy were first loaded with GTP by incubation in the presence of 20-fold excess GTP (Guanosine 5′-Triphosphate, Disodium Salt, CAT # 371701, Calbiochem) in 50 mM Tris HCl pH 7.5, 100 mM NaCl, 5 mM MgCl_2_. Exchange of GDP for GTP was initiated by the addition of 10 mM EDTA. Reactions were incubated for 3 hours at 4°C and stopped by addition of 1 M MgCl_2_ to a final concentration of 20 mM MgCl_2_ to quench the EDTA. GTP-loaded protein was buffer exchanged into either NMR buffer or the GTPase assay buffer using NAP-5 Sephadex G-25 DNA Grade columns (GE Healthcare # 17085301). We were unable to obtain sufficient material for some mutants (H141E/I, Y148I), for which we collected AP-MS data, since these mutants precipitated during the nucleotide exchange process at the high concentrations required for ^31^P NMR, possibly because of the limited stability of nucleotide-free Ran/Gsp1 generated during exchange, as noted previously(60).

#### Reverse phase high performance liquid chromatography (HPLC)

Analysis of bound nucleotide was performed using reverse-phase chromatography as previously described (59) using a C18 column (HAISIL TS Targa C18, particle size 5 μm, pore size 120 Å, dimensions 150 × 4.6 mm, Higgins Analytical # TS-1546-C185). The column was preceded by a precolumn filter (The Nest Group, Inc, Part # UA318, requires 0.5 μm frits, Part # UA102) and a C18 guard column (HAICart SS Cartridge Column, HAISIL Targa C18, 3.2×20 mm, 5μm, 120 Å Higgins Analytical # TF-0232-C185, requires a Guard Holder Kit, Higgins Analytical # HK-GUARD-FF). To prepare the nucleotide for analysis, a Gsp1 sample was first diluted to a concentration of 25-30 μM and a volume of 40 μl. The protein was denatured by addition of 2.5 μl of 10% perchloric acid (HClO_4_). The pH was raised by addition of 1.75 μl 4 M sodium acetate (CH_3_COONa) pH 4.0. The nucleotide was separated from the precipitated protein before application to the column by spinning at 20,000 × g for 20 minutes. 30 μl of supernatant was withdrawn and mixed 1:1 with reverse-phase buffer (10 mM tetra-n-butylammonium bromide, 100 mM KH_2_PO_4_ / K_2_HPO_4_, pH 6.5, 0.2 mM NaN_3_). 20 μl of sample was injected onto the equilibrated column, and was run isocratically in 92.5% reverse-phase buffer, 7.5% acetonitrile at a flow rate of 1 ml/min for 35 min (20 column volumes). Nucleotide retention was measured by monitoring absorbance at both 254 nm and 280 nm. Example HPLC reverse phase chromatogram of GTP-loaded wild type Gsp1 is shown in **Supplementary File 1 Fig. 14**.

#### NMR Spectroscopy

Gsp1 samples for 31P NMR spectroscopy were first loaded with GTP as described above, and buffer exchanged into NMR Buffer (D_2_O with 50 mM Tris-HCl pH 7.4, 5 mM MgCl_2_, 2 mM Dithiothreitol). Final sample concentrations were between 250 μM and 2 mM, and 400 ul of sample was loaded into 5 mm Shigemi advanced microtubes matched to D_2_O (BMS-005TB; Shigemi Co. Ltd, Tokyo, Japan.). ^31^P NMR experiments were performed on a Bruker Avance III 600 MHz NMR spectrometer with a 5 mm BBFO Z-gradient Probe. Spectra were acquired and processed with the Bruker TopSpin software (version 4.0.3). Indirect chemical shift referencing for ^31^P to DSS (2 mM Sucrose, 0.5 mM DSS, 2 mM NaN_3_ in 90% H_2_O + 10% D_2_O; water-suppression standard) was done using the IUPAC-IUB recommended ratios (61). Spectra were recorded at 25°C using the pulse and acquire program zg (TopSpin 3.6.0), with an acquisition time of 280 milliseconds, a recycle delay of 3.84 seconds, and a 65° hard pulse. 4,096 complex points were acquired over the course of 4,096 scans and a total acquisition time of 4.75 hours. Spectra were zero-filled once and multiplied with an exponential window function (EM) with a line-broadening of 6 Hz (LB = 6) prior to Fourier transformation. Peaks were integrated using the auto-integrate function in TopSpin 4.0.7, and peak areas were referenced to the bound GTP-β peak of each spectrum. Values for the fraction of each variant in state 2 were computed by taking the area of the GTP-γ2 peak and dividing by the sum of the two GTP-γ peak areas.

#### Kinetic measurements of GTP hydrolysis

Kinetic parameters of the GTP hydrolysis reaction were determined using a protocol similar to one previously described (62). Gsp1 samples for GTP hydrolysis kinetic assays were first loaded with GTP as described above. GTP hydrolysis was monitored by measuring fluorescence of the *E. coli* phosphate-binding protein labeled with 7-Diethylamino-3-[N-(2-maleimidoethyl)carbamoyl]coumarin (MDCC) (phosphate sensor, CAT # PV4406, Thermo Fisher) upon binding of the free phosphate GTP hydrolysis product (excitation at 425 nm, emission at 457 nm). All experiments were performed in GTPase assay buffer (40 mM HEPES pH 7.5, 100 mM NaCl, 4 mM MgCl_2_, 1 mM Dithiothreitol) at 30°C in 100 μl reaction volume using a Synergy H1 plate reader from BioTek, using Corning 3881 96-well half-area clear-bottom non-binding surface plates. The phosphate sensor at 20 μM and 50 μM concentrations was calibrated with a range of concentrations of K_2_HPO_4_ using only the data in the linear range to obtain a conversion factor between fluorescence and phosphate concentration. For each individual GAP-mediated GTP hydrolysis experiment, a control experiment with the same concentration of GTP-loaded Gsp1 and the same concentration of sensor, but without added GAP, was run in parallel. The first 100 s of these data were used to determine the baseline fluorescence, and the rest of the data were linearly fit to estimate intrinsic GTP hydrolysis rate (**Supplementary File 1 Table 8**). Although we do estimate the intrinsic hydrolysis rates from the background data, the estimate is only approximate, as well as 10^5^ to 10^6^ lower than the rate of GAP-mediated GTP hydrolysis, which is why we do not use intrinsic hydrolysis rates when fitting the GAP-mediated hydrolysis data. The affinity of Rna1 for GDP-bound Ran is negligible (Kd of 100 μM for Ran:GDP62), which is 250-fold weaker than the estimated K_m_ for GAP-mediated GTP hydrolysis and was not taken into account when fitting the data. As the estimated K_m_ for the GAP-mediated hydrolysis for many of the Gsp1 variants was low (in the 0.1-0.4 μM range, resulting in difficulties to reliably measure hydrolysis at low substrate concentrations), we sought to estimate the kinetic parameters (k_cat_ and K_m_) by directly analysing the full reaction progress curve with an analytical solution of the integrated Michaelis-Menten equation (see section below for details).

#### Estimating the k_cat_ and K_m_ parameters of GAP-mediated hydrolysis using an accurate solution to the integrated Michaelis-Menten equation

Others (e.g. Goudar et al (63)) have shown that both k_cat_ and K_m_ can be estimated with reasonable accuracy/precision from a single time-course with initial [S] > K_m_ by directly analyzing the full reaction progress curve with an analytical solution of the integrated Michaelis-Menten equation based on the Lambert ω function. This analysis is possible because the full reaction progress curve is characterized by an initial linear phase for [S] > K_m_, a final exponential phase for [S], and a transition phase for [S] K_m_. Whereas k_cat_ is sensitive to the slope of the initial linear phase (i.e. the initial velocity), K_m_ is sensitive to the shape of the progress curve, which will have an extended linear phase if K_m_ « initial [S] or no linear phase if K_m_ » initial [S]. Use of the integrated Michaelis-Menten analysis requires the experiment to be set up with the following conditions: (i) [Gsp1:GTP0] > K_m_, (ii) [GAP0] «< [Gsp1:GTP0], and (iii) the reaction time course F(t) is measured to completion (i.e. until it approached equilibrium). Our experiments were all set up to fulfill those conditions, which means that the F(t) sampled a concentration range from [Gsp1:GTP] (at t = 0) > K_m_ to [Gsp1:GTP] (at t = final time) « K_m_. The entire F(t) can then be directly analyzed by a non-linear fit with the analytical solution for the integrated Michaelis-Menten equation. As the initial linear phase of the time course is well measured, k_cat_ can be well determined. As the exponential phase and transition region of the time course are also well measured, the maximum likelihood value of K_m_ can also be determined. Specifically, each time course was fitted to an integrated Michaelis Menten equation:

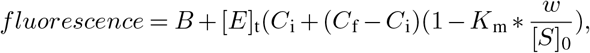

where [E]_t_ is the total enzyme (GAP) concentration, C_i_ is the initial fluorescence, C_f_ is the final fluorescence, [S]_0_ is the initial concentration of the substrate (GTP loaded Gsp1), and B is the baseline slope in fluorescence per second. Exact concentration of loaded Gsp1:GTP [S]_0_ was estimated based on the plateau fluorescence and the sensor calibration parameters to convert the fluorescence to free phosphate concentration. The ω parameter was solved by using the Lambert ω algorithm, as previously described(63), where,

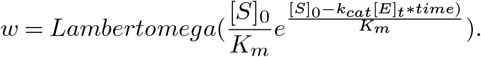

The curves were fit with the custom-made software DELA (64). Examples of full reaction progress curves and their integrated Michaelis-Menten fits are shown in **Supplementary File 1 Fig. 3.** We confirmed that the kinetic value parameters we obtained for wild-type Gsp1 using the phosphate sensor and integrated Michaelis-Menten equation were similar to those estimated using intrinsic tryptophan fluorescence (65). Their values were a K_m_ of 0.45 μM and k_cat_ of 2.1 s^−1^ at 25°C for mammalian Ran hydrolysis activated by *S. pombe* GAP, while our values for wild type *S. cerevisiae* Gsp1 and *S. pombe* GAP at 30°C are K_m_ of 0.38 μM and k_cat_ of 9.2 s^−1^. For most mutants a concentration of 1 nM GAP (SpRna1, Rna1 from *S. pombe*) was used. In order to run the time courses to completion, for mutants with low k_cat_/K_m_ enzyme concentrations of 2-5 nM were used. Initially we collected time course data for all Gsp1 variants at approximately 8 μM concentration of loaded Gsp1:GTP with 1 nM GAP and 20 μM phosphate sensor. If the estimated K_m_ was higher than 1 μM, we repeated the time course kinetic experiments with higher concentration of Gsp1:GTP of approximately tenfold above the K_m_. To quantify the accuracy of parameter (k_cat_, K_m_) estimation for GAP-mediated GTP-hydrolysis by the integrated Michaelis-Menten approach over a range of kinetic parameters and substrate concentrations [Gsp1:GTP] we simulated data covering the range of parameters estimated for all of our Gsp1 point mutants, and estimated the accuracy of parameters determined given the Gaussian noise similar to our experimental data. The largest standard deviations were 3%, 17%, and 18% for k_cat_, K_m_, and k_cat_/K_m_, respectively (**Supplementary File 1 Fig. 15**). In addition, we analysed how the χ^2^ statistic changed as the Michaelis-Menten parameters were systematically varied around the estimated maximum likelihood values (**Supplementary File 1 Fig. 16**). For these analyses, the k_cat_ or K_m_ values were independently fixed and incremented while the remaining parameters were fit to generate χ^2^ surfaces for one degree of freedom. Confidence intervals (CIs) for which χ^2^ increased by 4.0 compared to the maximum likelihood minimum were estimated by linear interpolation after iterative bisection. A χ^2^ increase of 4.0 corresponds to the 95% confidence limit for a normal distribution. The k_cat_/K_m_ ratio and corresponding χ^2^ values were derived from the analyses with systematic variation of either k_cat_ or K_m_. CIs for k_cat_/K_m_ were estimated by linear interpolation without iterative bisection. The χ^2^ surfaces approach a parabolic shape with a well-defined minimum at the maximum likelihood value. The CIs are further consistent with the parameter ranges obtained from the simulations. Thus, both the simulations and χ^2^ surfaces indicate that k_cat_ and K_m_ are estimated with reasonable accuracy over the range of parameter values and experimental conditions used in this study. The Michaelis-Menten k_cat_ and K_m_ parameters and their standard deviations were calculated from at least three technical replicates from two or more independently GTP-loaded Gsp1 samples (**Supplementary File 1 Table 6**).

#### Kinetic measurements of Srm1 mediated nucleotide exchange

Kinetic parameters of GEF mediated nucleotide exchange were determined using a fluorescence resonance energy transfer (FRET) based protocol (65). Each Gsp1 variant was purified as a Gsp1:GDP complex, as confirmed by reverse phase chromatography. Nucleotide exchange from GDP to mant-GTP (2’-(or-3’)-O-(N-Methylanthraniloyl) Guanosine 5′-Triphosphate, CAT # NU-206L, Jena Biosciences) was monitored by measuring a decrease in intrinsic Gsp1 tryptophan fluorescence (295 nm excitation, 335 nm detection) due to FRET upon binding of the mant group. Each time course was measured in GEF assay buffer (40 mM HEPES pH 7.5, 100 mM NaCl, 4 mM MgCl_2_, 1 mM Dithiothreitol) with excess of mant-GTP. The affinity of Ran/Gsp1 is estimated to be 7-11-fold lower for GTP than for GDP (60), and for most variants of Gsp1 we measured time courses at Gsp1:GDP concentrations ranging from 0.25 to 12 μM with an excess mant-GTP concentration of 200 μM. For Gsp1 variants with high Km values that had to be measured at concentrations of up to 200 μM we used an excess of 1000 μM mant-GTP. In addition, we fit the data using a combination of fits following the approach of Klebe (65). For concentrations of substrate (Gsp1:GDP) that were much lower than the excess of mant-nucleotide (200 μM) we used a combination of two exponential decays, and for reactions with high concentrations of Gsp1, where the relative excess of mant-nucleotide was lower, we always estimated the initial rates using linear fits to the very beginning of the reaction, when levels of mant-nucleotide-bound Gsp1 are very low and therefore exchange is overwhelmingly from Gsp1-GDP to Gsp1-mant-nucleotide. All kinetic measurements were done at 30°C in 100 μl reaction volume using 5 nM GEF (Δ1-27Srm1), except for higher concentrations of the mutants with high K_m_ values that were measured at 20 nM GEF. Data were collected in a Synergy H1 plate reader from BioTek, using Corning 3686 96-well half-area non-binding surface plates. For low concentrations of Gsp1:GDP the time course data were fit to a combination of two exponential decays:

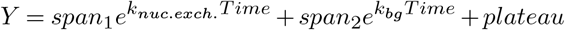

where k_nuc. exch_ is the rate constant of the GDP to mant-GTP exchange, k_bg_ is the rate constant of background decay due to photo-bleaching, and span_1_ and span_2_ are the fluorescence amplitudes for the two processes. For high concentrations of substrate, or for mutants with very low rates, the initial velocity was determined by a linear fit to the initial 10-20% of the data. As the intrinsic exchange rate in the absence of GEF is estimated to be more than 10^4^ lower (60) we do not use the intrinsic rate for fitting the data. The kinetic parameters of the nucleotide exchange were determined by fitting a Michaelis-Menten equation to an average of 38 data points (ranging from 17 to 91) per Gsp1 point mutant for a range of substrate concentrations from [Gsp1:GDP] = 0.25 μM to [Gsp1:GDP] » K_m_. Michaelis-Menten fits are shown in **Supplementary File 1 Fig. 4**. Michaelis-Menten k_cat_ and K_m_ parameters for GEF-mediated nucleotide exchange are provided in **Supplementary File 1 Table 7**. The errors of the k_cat_ and the K_m_ parameters were determined from the standard error of the exponential fit of the Michaelis-Menten equation to the data. The error of the catalytic efficiency (k_cat_/K_m_) was calculated by adding the standard errors of the individual parameters and normalizing it for the values of the parameters

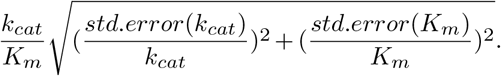

All the custom written code for fitting and analysis of kinetics data is provided in the accompanying kinetics GitHub repository.

## Acknowledgements

We thank Roxana Ordonez for contributions to the design of mutations; Cristina Melero, Deborah Jeon, Shivani Mathur, Raina Danbi Kim, and Kale Kundert for technical help; Maru Jaime Garza for contributions to the conformational analysis by NMR; Colm Ryan for advice on E-MAP analysis; and Dave Agard, Geeta Narlikar, James Fraser and Janet M. Thornton for stimulating discussions; Ricardo Henriques and his lab for the preprint template. This work was supported by a grant from the National Institutes of Health (R01-GM117189) to T.K., and a Sir Henry Wellcome Postdoctoral Fellowship (101614/Z/13/Z) to T.P. C.J.P.M. is a UCSF Discovery Fellow. T.K. is a Chan Zuckerberg Biohub investigator.

## Author Contributions

T.P., C.J.P.M., N.J.K. and T.K. identified and developed the core questions. T.P. and C.J.P.M. performed the bulk of the experiments and data analysis. J.X. and T.P. performed the E-MAP screens. G.M.J. performed the pull-down experiments. D.L.S. and R.K. performed the MS experiments and together with T.P. analyzed the data. N.O. contributed to design of Gsp1 mutants. H.B. contributed to E-MAP analysis. M.J.S.K. suggested the NMR studies. C.J.P.M. and M.J.S.K. performed the NMR experiments and analyzed the data. Y.Z. performed the Western blot experiments. T.P., C.J.P.M., and prepared the proteins. T.P. performed the kinetics experiments. D.G.L. contributed to the analysis of the kinetics data. T.P., C.J.P.M. and T.K. wrote the manuscript with contributions from the other authors. N.J.K. and T.K. oversaw the project.

## Data Availability

The mass spectrometry proteomics data have been deposited to the PRIDE proteomics data repository with the dataset identifier PXD016338 and are available as Supplementary Tables. Raw biophysics data (cycle kinetics, CD, and NMR), and E-MAP S-scores, scaled SGA (48) scores, and their correlations are available from a GitHub repository All other data that support the findings of this study are available within the paper and its Supplementary Files.

## Code Availability

Custom written R and Python scripts are available without restrictions from a a Gsp1 manuscript GitHub repository

## Competing Interests

The authors declare no competing interests.

---

**Fig. S1.**
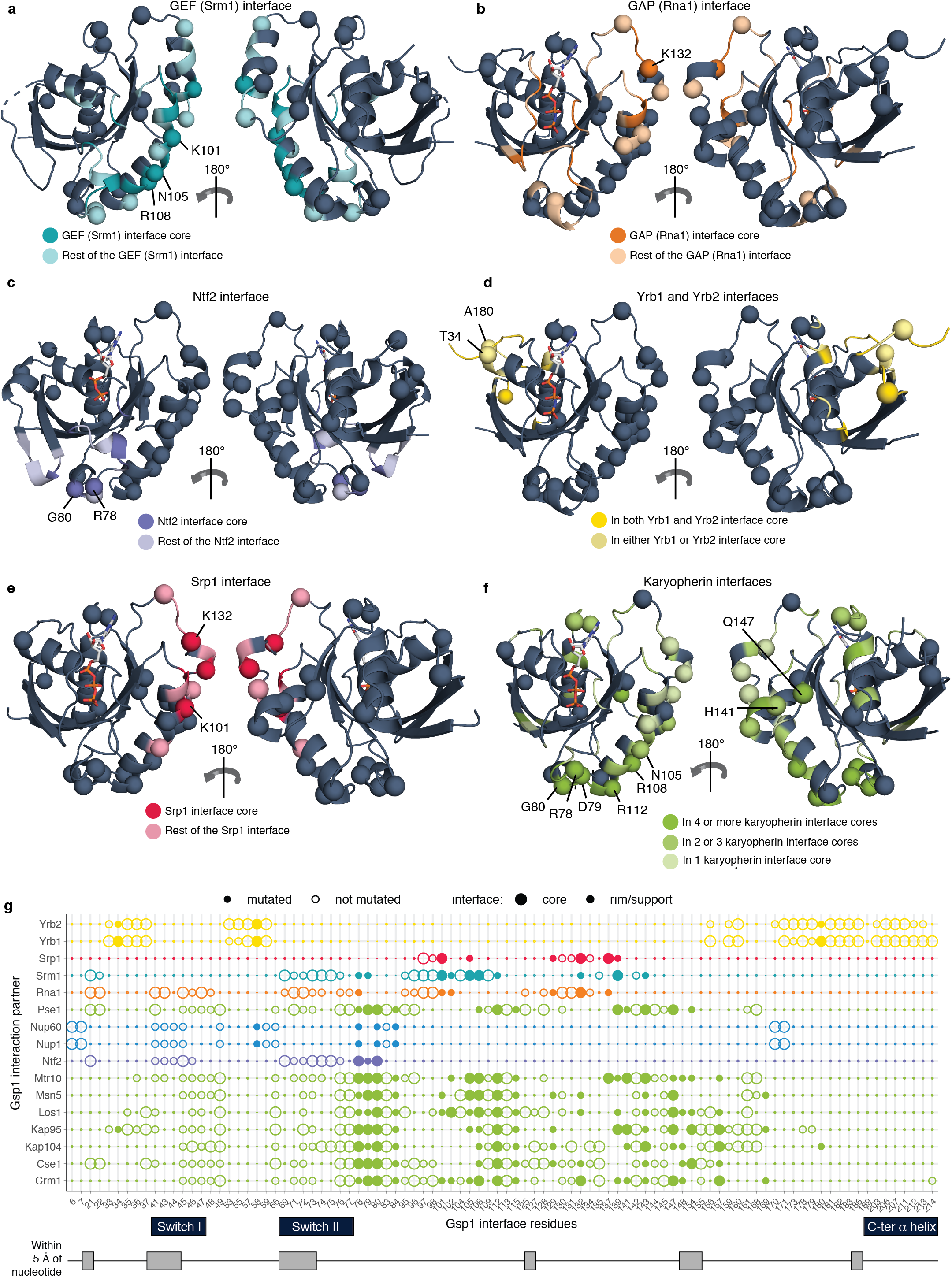
Design of interface point mutations in *S. cerevisiae* Gsp1. Interface core, rim, and support positions are defined as in (5) (see **Methods**) and provided in **Supplementary File 1 Table 2.** **a-f,** Structures of Ran/Gsp1 in different binding conformations. Mutated Gsp1 residues are shown as spheres. Interface residues are coloured by the type of partner protein: **a,** Srm1 (GEF) interface core (dark teal) and interface rim and support (light teal) PDB ID: 1i2m; **b**, Rna1 (GAP) interface core (dark orange) and interface rim and support (light orange) PDB ID: 1k5d; **c,** Ntf2 interface core (dark purple) and interface rim and support (light purple) PDB ID: 1a2k; **d,** Residues that are in both the core of the Yrb1 and Yrb2 interfaces (dark yellow), and in only one of the two interfaces (light yellow) PDB ID: 1k5d; **e,** Srp1 interface core (dark pink) and interface rim and support (light pink) PDB ID: 1wa5; **f,** Residues that are in the core of more than four (dark green), two to three (green) and one (light green) karyopherin interface. Karyopherins are: Kap95, Crm1, Los1, Kap104, Msn5, Cse1, Mtr10. PDB ID: 2bku. **g,** Location of Gsp1 residues in partner interfaces. Residues within 5 Å of the nucleotide, in the canonical P-loop, or in the switch I or II regions are indicated and were not mutated. Chosen Gsp1 point mutation substitutions are provided in **Supplementary File 1 Table 3**.

**Fig. S2.**
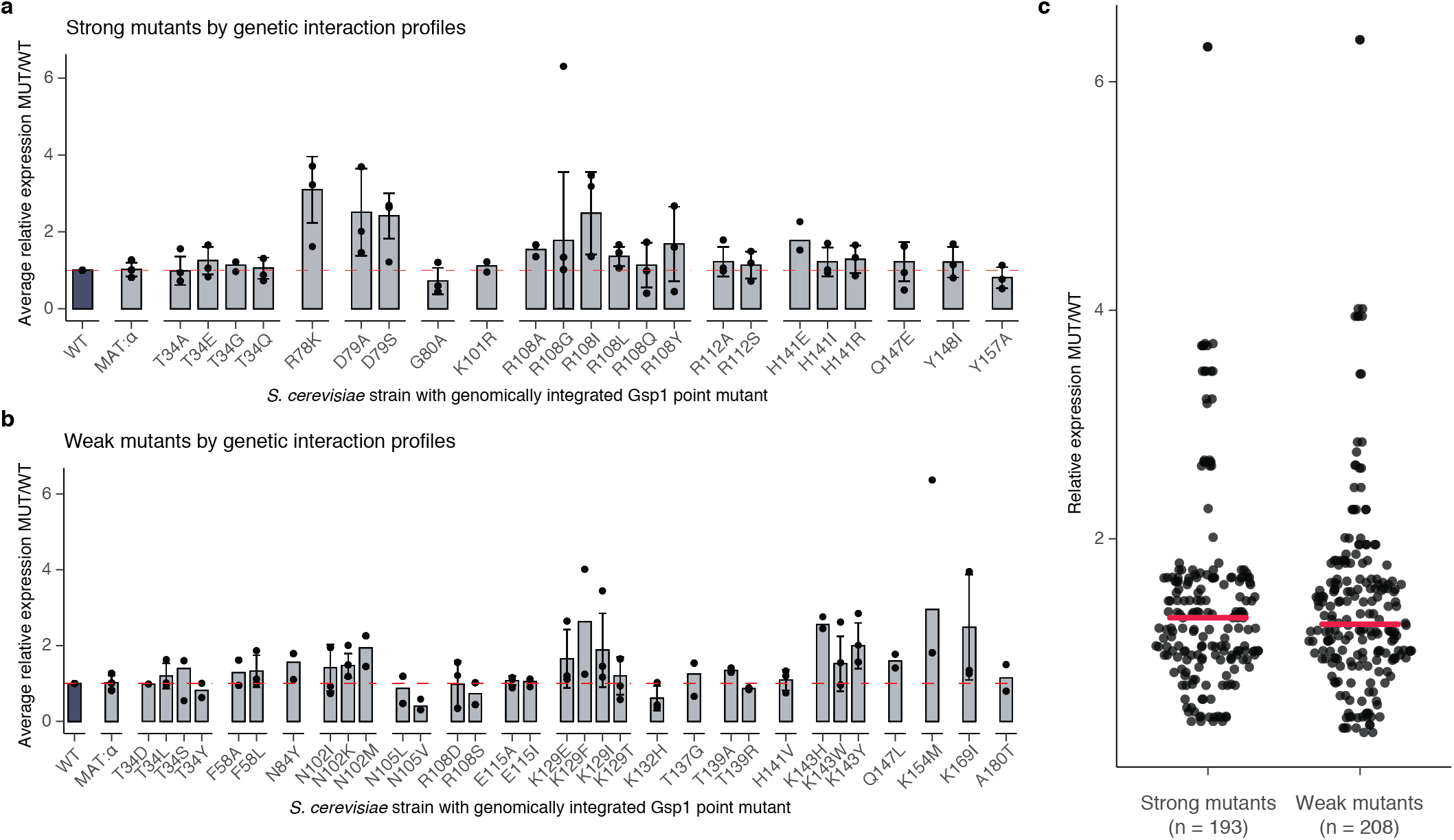
Expression levels of endogenously expressed Gsp1 protein in *S. cerevisiae* strains with genomically integrated Gsp1 point mutations profiled by Western Blot. Expression levels are relative to the expression levels of wild-type Gsp1 protein. **a,** Expression data for strong mutants, defined as mutants with more than nine significant GIs. **b,** Expression data for weak mutants, defined as mutants with fewer than nine significant GIs. Bar heights indicate averages over 2 or more biological replicates (n) with error bars indicating one standard deviation for n >= 3. Overlaid points indicate individual biological replicates (each an average over at least 12 technical replicates per biological replicate for wild-type and MAT-α strains, and between one and six technical replicates per biological replicate for mutant strains). Dashed red line indicates expression at the level of wild-type Gsp1 (fold change of 1). **c,** Distributions of average relative expression levels for strong and weak mutants. Each point is as in **a** and **b**. Horizontal pink bars indicate the mean of the point distributions.

**Fig. S3.**
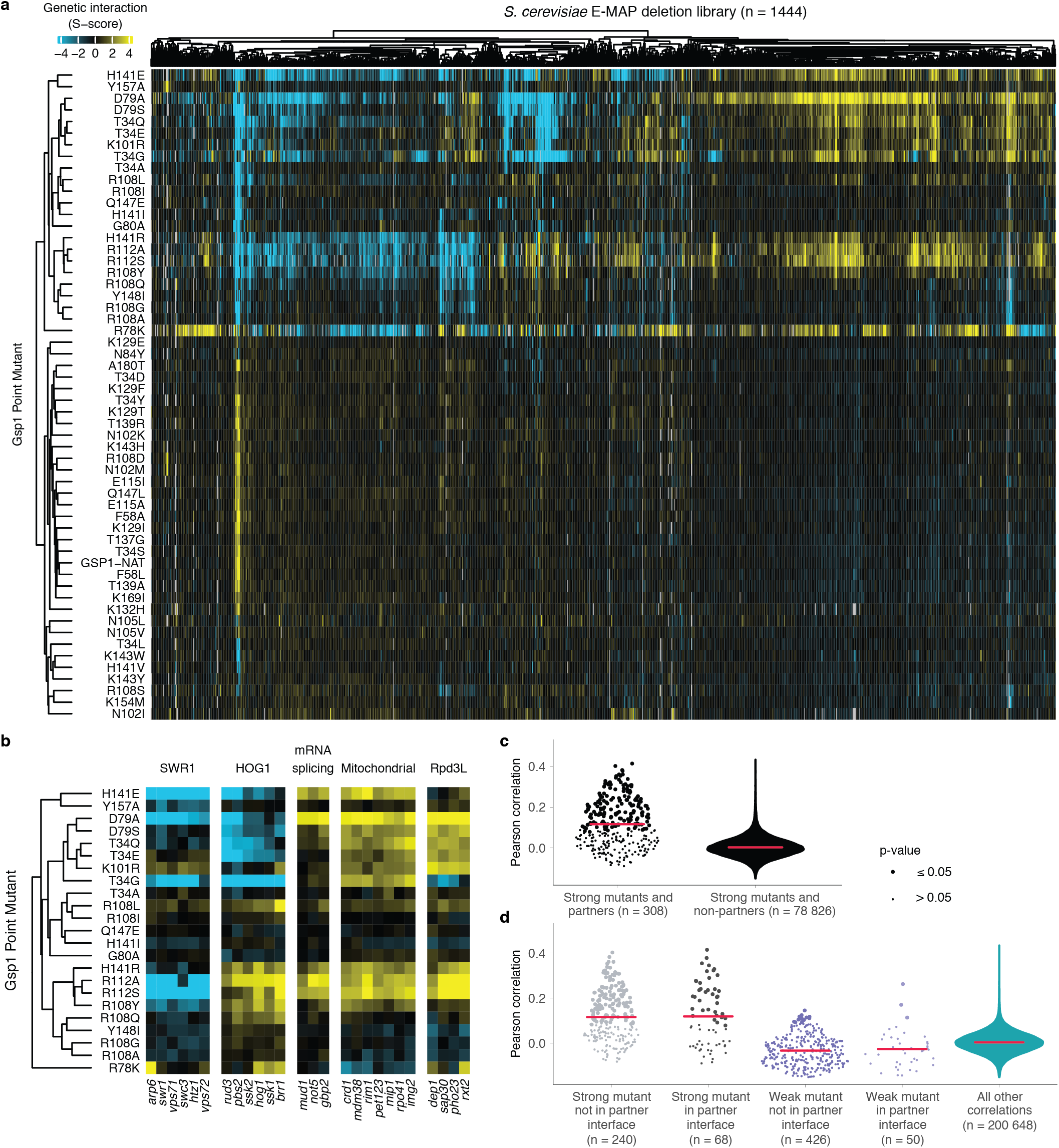
Genetic interaction (GI) profiles of the 55 Gsp1 point mutations. **a,** Complete Gsp1 E-MAP profile. Negative S-score (blue) represents synthetic sick/lethal GIs, positive S-score (yellow) represents suppressive/epistatic GIs; neutral S-scores (no significant GI) are shown in black. Mutants and genes are hierarchically clustered by Pearson correlation. Gsp1 mutants fall into two clusters: a cluster of 23 strong mutants with nine or more significant GIs (blue and yellow S-scores in the heatmap) and 23 weak mutants with fewer than nine significant GIs (mostly black S-scores in the heatmap). **b,** GI profiles of Gsp1 mutants group *S. cerevisiae* genes by biological processes and complexes, such as the SWR1 complex, the Hog1 signaling pathway, mRNA splicing, mitochondrial proteins, and the Rpd3L histone deacetylase complex. **c,** Distributions of Pearson correlations between the GI profiles of strong Gsp1 mutants and alleles of Gsp1 direct interaction partners with available co-complex crystal structures (left, **Fig. 5a**) and strong Gsp1 mutants and all other *S. cerevisiae* genes (right). **d,** Distributions of Pearson correlations between the GI profiles of Gsp1 interaction partners and strong and weak Gsp1 mutants if mutation is (black and light purple) or is not (gray and dark purple) in the interface with that partner. Teal violin plot on the right represents the distribution of all other Pearson correlations between Gsp1 mutants and *S. cerevisiae* genes. In c and d, point size indicates the false discovery rate adjusted one-sided (positive) p-value of Pearson correlation. Red dots and bars indicate the mean and the upper and lower quartile, respectively.

**Fig. S4.**
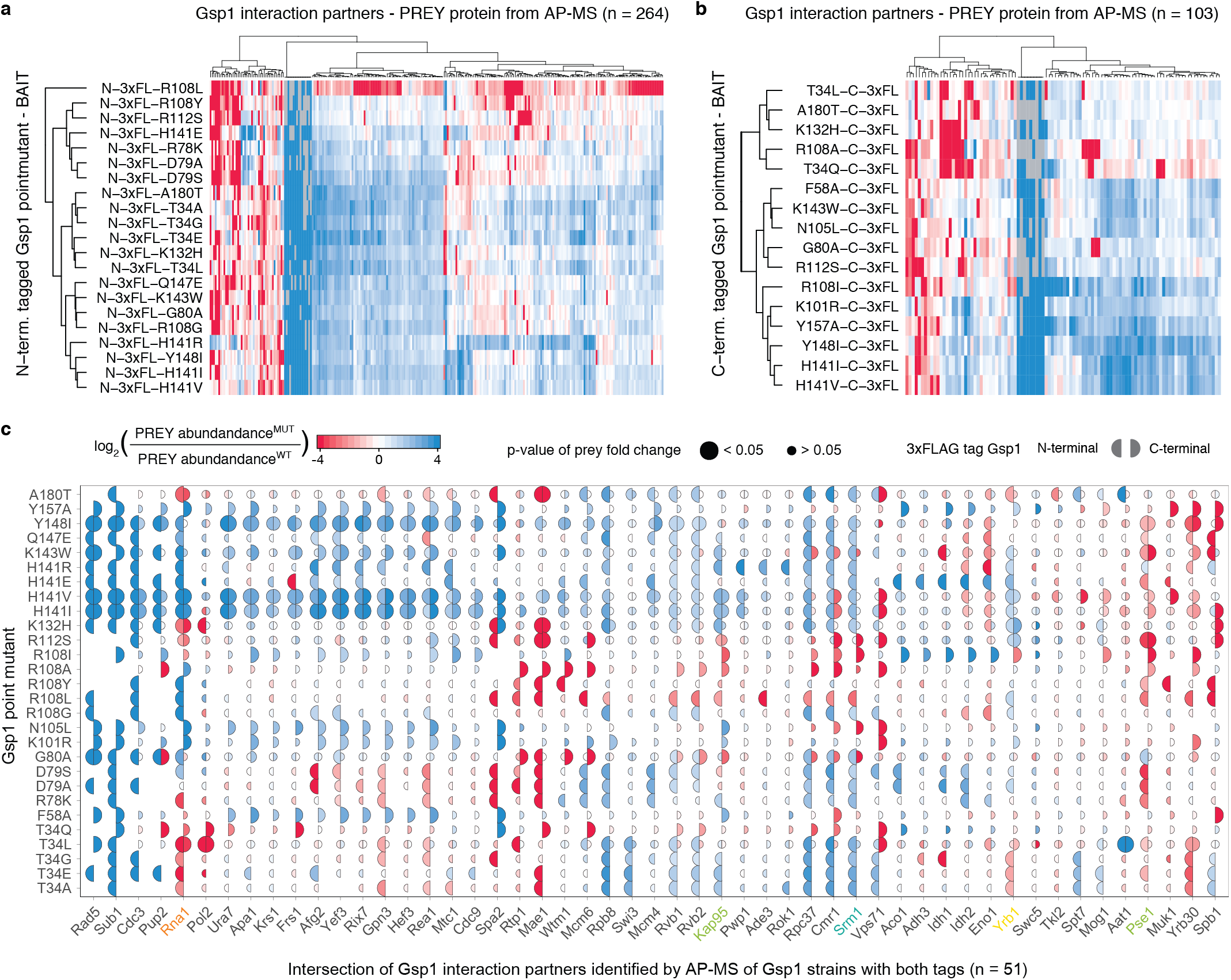
Interface point mutations in Gsp1 rewire its physical interaction network. **a,** Amino- and **b,**-carboxy terminally 3xFLAG-tagged Gsp1 point mutants (rows) and prey proteins identified by AP-MS (columns) hierarchically clustered by the log_2_ - transformed fold change in prey abundance pulled-down with either the mutant or wild-type Gsp1 with the corresponding 3xFLAG-tag (log_2_ (abundance(PREY)^MUT^/abundance(PREY)^WT^)). **c,** Prey proteins pulled down by both amino- and carboxy-terminal tagged constructs. Left semi-circle represents an amino-terminal 3xFLAG-tagged Gsp1 point mutant, and right semi-circle represents carboxy-terminal 3xFLAG-tagged Gsp1 point mutant. Semi-circle size is proportional to the significance of the log_2_ - transformed fold change (false discovery rate adjusted p-value) of the prey abundance in pulled-down complexes with a Gsp1 mutant compared to complexes with the wild-type Gsp1. Overall we identified 316 high-confidence prey partner proteins, with the amino- and carboxy-terminally tagged Gsp1 mutants pulling down 264 and 103 preys, respectively, including 51 overlapping preys. The difference in preys identified by experiments with N- or C-terminal tags illustrates the sensitivity of the interaction network to perturbation of Gsp1. To account for possible tag effects, we always computed the fold change in prey abundance only relative to the wild-type protein with the corresponding tag. In **a, b,** and **c,** decreased abundance compared to pull-down with wild-type Gsp1 is annotated in red and increased abundance in blue. The log_2_ - transformed fold change values are capped at +/− 4.

**Fig. S5.**
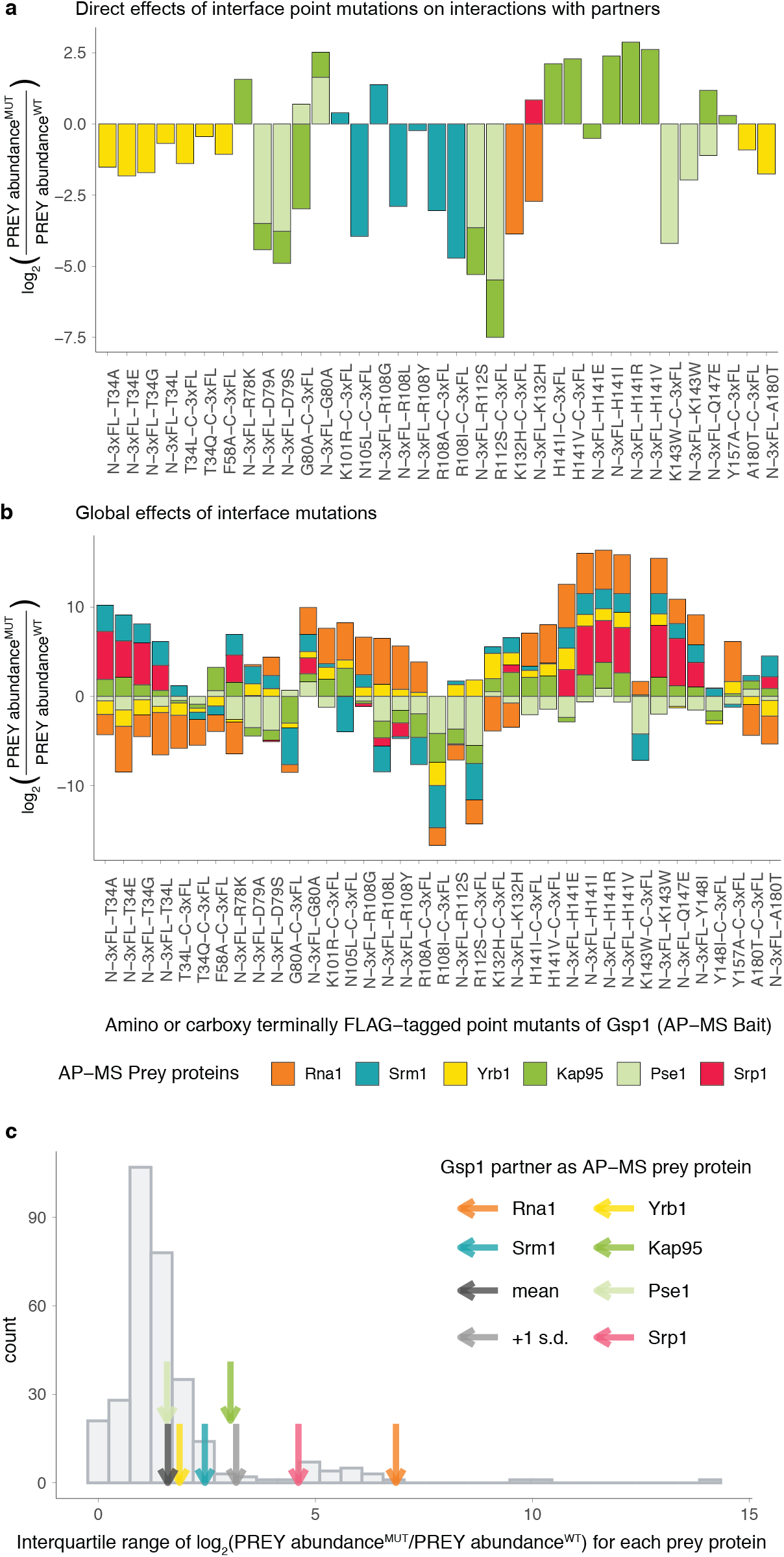
Protein-protein interactions between interface mutants of Gsp1 and Gsp1 partners for which there are co-complex X-ray crystal structures (core regulators Srm1 and Rna1, and effectors Yrb1, Kap95, Pse1, and Srp1). **a, b**, Change in pulled-down prey partner abundance is expressed as log_2_ (PREY abundance^MUT^/PREY abundance^WT^)). N-3xFL and C-3xFL labelled mutants are tagged with an amino- or carboxy-terminal triple FLAG tag, respectively, and partners are coloured as indicated. **a,** Bar plot depicting changes in pulled-down prey partner abundance when the point mutation is in the core of the Gsp1 interface with the prey partner. **b,** Bar plot depicting all changes in pulled-down prey partner abundance for core regulators Srm1 and Rna1, and effectors Yrb1, Kap95, Pse1, and Srp1, regardless whether the mutation is directly in the interface core with the partner or not. **c,** Distribution showing the variation in log_2_ - transformed fold change in abundance of all prey proteins pulled down with the Gsp1 mutants, as defined by interquartile range (IQR) across mutants. Values for core partners shown as arrows (Rna1 orange, Srm1 teal, Yrb1 yellow, Kap95 green, Pse1 light green, Srp1 pink). Mean and +1 standard deviation of IQR values are highlighted with a dark gray and a light gray arrow, respectively. The extent to which the abundance of the two cycle regulators Rna1 and Srm1 changed across the Gsp1 point mutants was significantly larger than the change of an average prey protein. All IQR values are provided in **Supplementary File 1 Table 5.**

**Fig. S6.**
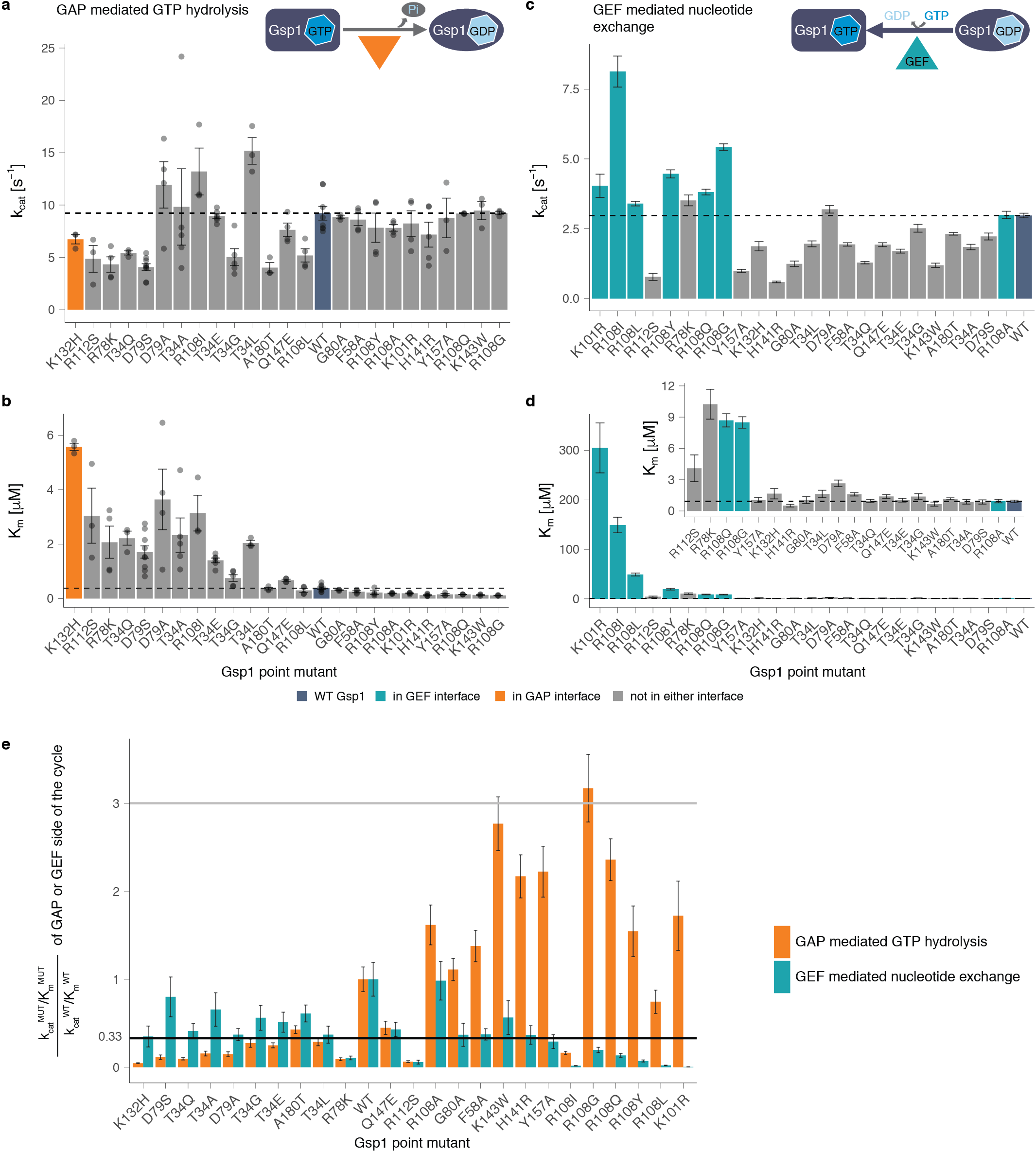
Effect of Gsp1 point mutations on the *in vitro* efficiency of GAP-mediated GTP hydrolysis and GEF-mediated nucleotide exchange. **a**, k_cat_ and **b,** K_m_ values of GAP-mediated GTP hydrolysis of wild-type and point mutant Gsp1. Error bars represent the standard deviation of the k_cat_ and the K_m_ parameters from the integrated Michaelis-Menten fit for n >= 3 replicates. **c**, k_cat_ and **d,** K_m_ of GEF-mediated nucleotide exchange of wild-type and point mutant Gsp1. Inset shows the K_m_ barplot for all but the four mutants with the highest K_m_ (K101R, R108L, R108I, and R108Y). Error bars represent the value plus/minus the standard error of the Michaelis-Menten fit to data from n >= 17 measurements at different substrate concentrations. **a, b, c, d,** Dotted lines indicate the wild-type values. Dark blue bar denotes the wild-type Gsp1, and orange and teal bars highlight the residues that are in the core of the interface with the GAP and GEF, respectively. **e,** Comparison of relative change in catalytic efficiencies of the GAP-mediated GTP hydrolysis (orange bars) and GEF-mediated nucleotide exchange (teal bars) defined as k_cat_^MUT^/K_m_^MUT^/ k_cat_^WT^/K_m_^WT^. Gray line indicates a three-fold increase compared to wild type, black line indicates a three-fold decrease compared to wild type. Error bars represent the added standard error of the mean (for GAP) or standard error of the fit (for GEF) values of the mutant and the wild-type efficiency (k_cat_/K_m_) values.

**Fig. S7.**
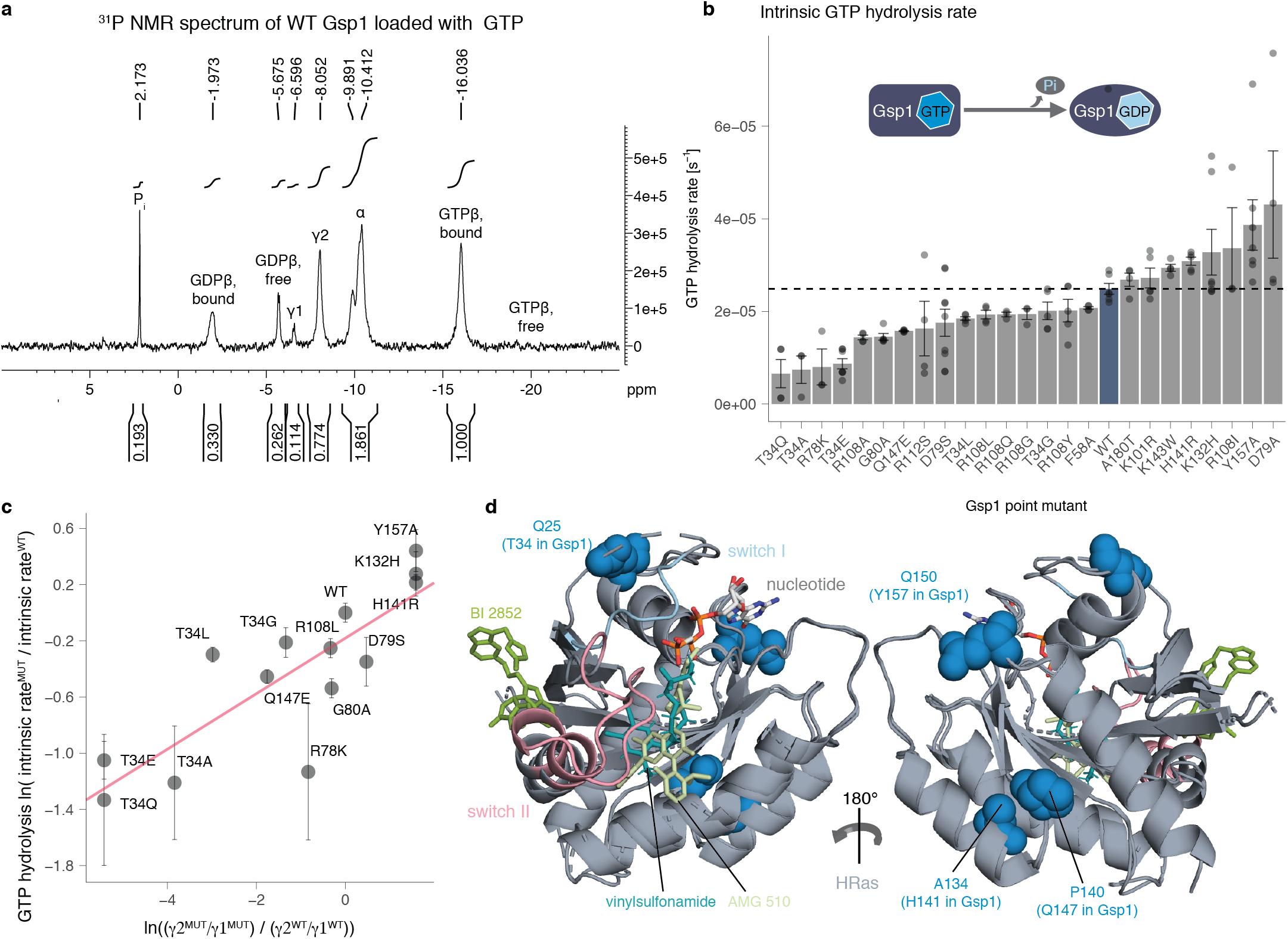
Gsp1 interface mutations act allosterically to modulate the rate of GTP hydrolysis. **a,** Annotated 1D 31 P NMR spectrum of wild-type Gsp1 loaded with GTP. Peak areas are computed over intervals shown and normalized to the GTPβ bound (GTPβbound) peak. The peaks from left to right correspond to: free phosphate (Pi), β phosphate of GDP bound to Gsp1 (GDPβbound), β phosphate of free (unbound) GDP (GDPβfree), γ phosphate of GTP bound to Gsp1 in conformation 1 (γ1), γ phosphate of GTP bound to Gsp1 in conformation 2 (γ2), α phosphate of bound or unbound GDP or GTP, β phosphate of GTP bound to Gsp1 (GTPβbound), β phosphate of free (unbound) GTP (GTPβfree). **b,** Rate of intrinsic GTP hydrolysis of wild-type Gsp1 and mutants. Dotted line indicates wild-type value. Error bars represent the standard deviations from n >= 3 replicates. **c,** Natural log-transformed exchange equilibrium constant between the γ2 and γ1 conformations plotted against the relative rate of intrinsic GTP hydrolysis represented as a natural logarithm of the ratio of the rate for the mutant over the rate of the wild type. The pink line is a linear fit. Error bars represent the standard deviation from n >= 3 replicates of intrinsic GTP hydrolysis measurements. **d,** Structures of HRas (in cartoon representation, gray) bound to inhibitors shown in stick representation: BI 2852 (PDB ID: 6gj8, green), vinylsulfonamide (PDB ID: 4m1w, teal), and AMG 510 (PDB ID: 6oim, light green). Switch I and switch II regions of HRas are in light blue and pink, respectively. Human HRas residues corresponding to Gsp1 allosteric sites (identified from the sequence alignment between Gsp1 and human HRas) are represented as blue spheres. The corresponding Gsp1 residues are in parentheses.

**Fig. S8.**
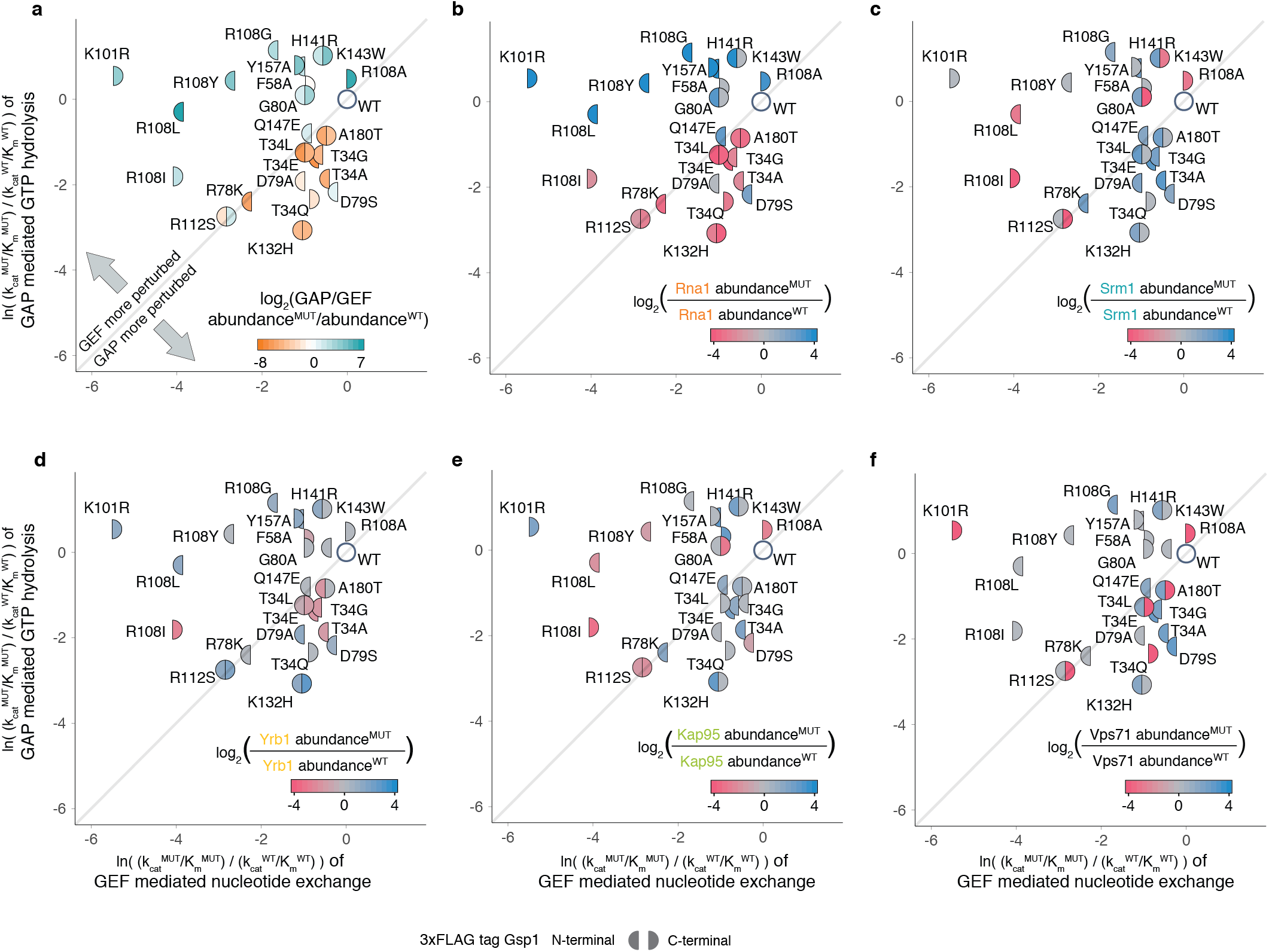
Relative prey protein abundance compared to the wild type with corresponding 3xFLAG tag from the AP-MS proteomics experiment overlaid onto the effects of each mutation on relative *in vitro* efficiencies of GAP-mediated GTP hydrolysis and GEF-mediated nucleotide exchange. Relative GAP-mediated hydrolysis and GEF-mediated exchange efficiencies are plotted as ln(k_cat_^MUT^/K_m_^MUT^/k_cat_^WT^/K_m_^WT^). Left semi-circle represents an amino-terminal 3xFLAG-tagged Gsp1 point mutant, and right semi-circle represents carboxy-terminal 3xFLAG-tagged Gsp1 point mutant, relative to wild-type Gsp1 with the corresponding tag. **a,** Color represents log_2_ - transformed ratio of GAP and GEF abundance fold change for each Gsp1 point mutant compared to wild type defined as log_2_ ((abundance(Rna1)^MUT^/abundance(Rna1)^WT^)/(abundance(Srm1)^MUT^/abundance(Srm1)^WT^)). Orange coloured mutants pull-down relatively less Rna1 (GAP) and teal mutants less Srm1 (GEF). **b-f**, Colour represents the log-transformed ratio of mutant and wild type pulled-down prey protein represented as log_2_ (PREY abundance^MUT^/PREY abundance^WT^). Log-transformed relative abundance values are capped at +/− 4. Decreased prey abundance from AP-MS in pulled-down complexes with a mutant Gsp1 compared to complexes with the wild-type Gsp1 is represented in red and increased abundance in blue. Prey proteins: **b,** Rna1 (GAP); **c,** Srm1 (GEF); **d,** Yrb1; **e,** Kap95, and **f,** Vps71. Yrb1 follows a pattern more similar to that of Rna1 (GAP), while Kap95 and Vps71 are more similar to Srm1 (GEF).

**Fig. S9.**
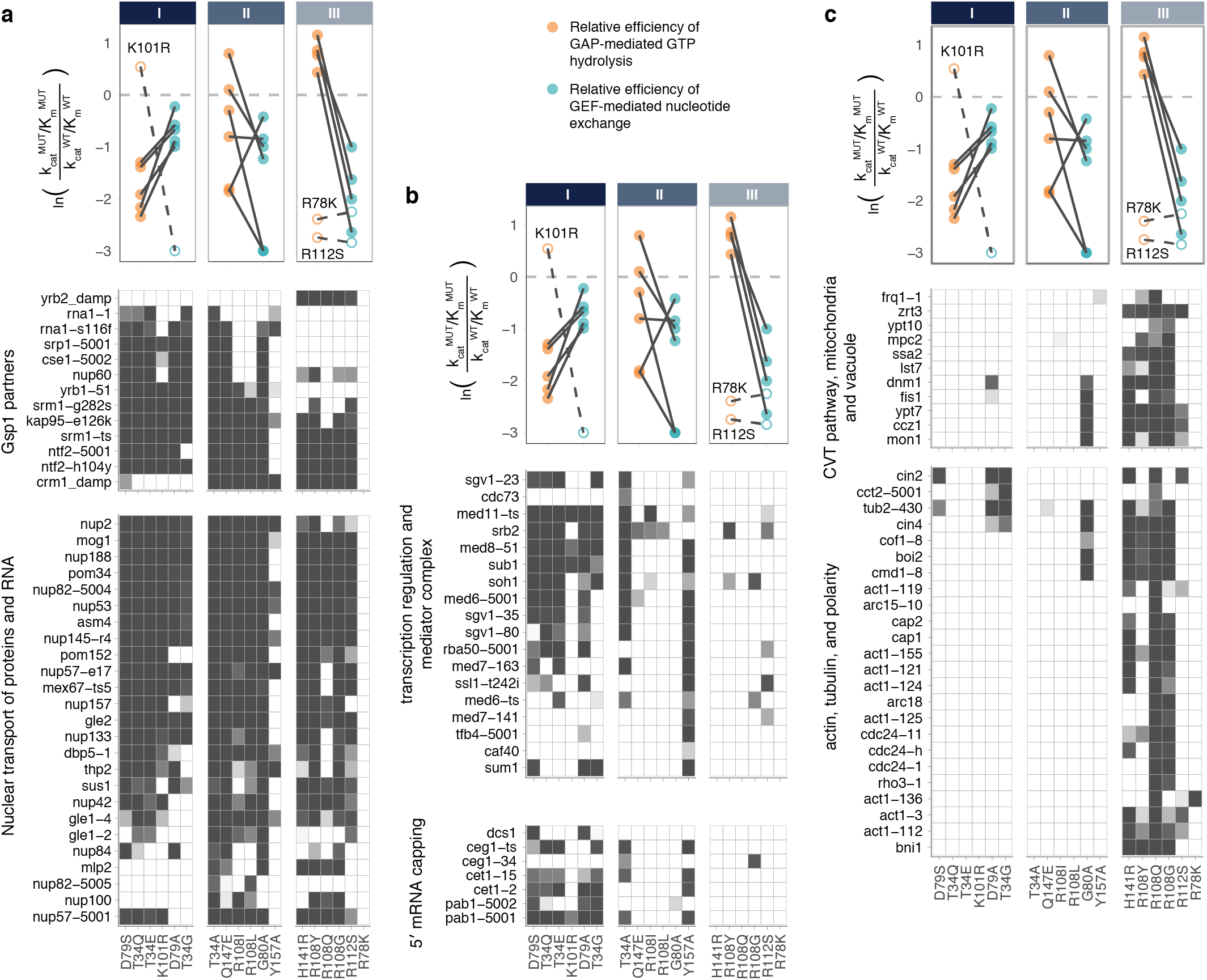
Sets of *S. cerevisiae* genes grouped by biological functions. Heatmaps of the false discovery rate adjusted one-sided (positive) p-values of the Pearson correlations between the GI profiles of 22 strong Gsp1 point mutants and GI profiles of knock-outs or knock-downs of *S. cerevisiae* genes from Ref.(21). The p-value is represented as a white to gray range, with gray being most significant. Genes are organized in gene sets based on their biological function (**Methods**). The line plots above the heatmaps are the same as in **Fig. 4c**. **a,** Gsp1 point mutants and alleles of Gsp1 binding partners with available co-complex X-ray crystal structures, and *S. cerevisiae* genes involved in nuclear transport of RNA and proteins. **b,** Gsp1 point mutants and *S. cerevisiae* genes involved in transcription regulation or 5′ mRNA capping. **c,** Gsp1 point mutants and *S. cerevisiae* genes involved in the cytoplasm-to-vacuole targeting (CVT) pathway, and actin, tubulin, and cell polarity.

## Notes

### Competing Interest Statement

The authors have declared no competing interest.

