## Supplementary File 1 for "Biophysical basis of cellular multi-specificity encoded in a model molecular switch"

### Supplementary Discussion

##### Validity of the Michaelis-Menten formalism for GTPases

Michaelis-Menten formalisms have been used for multiple GTPases including Ran[1], Ras[2], or Rap[3]. Historically there have been many attempts to formalize the conditions under which the Michaelis-Menten equation to describe enzyme kinetics are valid (as reviewed by Schnell[4]). These conditions have converged on the steady-state approximation or more generally, on the reactant stationary assumption. The formal condition for steady-state approximation is that $t_{[ES]}$ (the time it takes for the steady-state levels of [ES] complex to accumulate) is substantially shorter than $t_{[S]}$ (the time where [S] changes significantly). The formal condition for reactant stationary assumption is that $[S] \approx [S_0]$ during initial build-up of [ES].

The formal condition for validity of the Michaelis-Menten equation can be expressed as:

$$\frac{[E_0]}{K_m + [S_0]} \ll \left(1 + \frac{K}{K_S}\right)\left(1 + \frac{[S_0]}{K_m}\right),$$

where $K = \frac{k_{cat}}{k_{on}}$ and $K_S = \frac{k_{off}}{k_{on}}$, and $k_{off}$ and $k_{on}$ are the rates of [ES] complex formation[5].

The measured dissociation constant, $K_S = \frac{k_{off}}{k_{on}}$, for the formation of the Ran:GDP:RCC1 complex from Ran:GDP and RCC1, where RCC1 is the human RanGEF, is 0.9 µM[6], which is approximately the same as the Km value obtained for the GEF-mediated nucleotide exchange for both *S. cerevisiae* Gsp1 and human Ran. That means than $K \ll K_S$, which means the condition for validity of the Michaelis-Menten equation can be approximated as $\frac{[E_0]}{K_m + [S_0]} \ll \left(1 + \frac{[S_0]}{K_m}\right)$, and since in all of our GEF experiments both $[E_0]$ = 5-20 nM << $K_m$ and $[E_0]$ << $[S_0]$, the conditions holds true for the entire range of $[S_0]$ values, both below and above the $K_m$.

As $\frac{K}{K_S}$ can also be expressed as $\frac{k_{cat}}{k_{off}}$, and the measured $k_{off}$ of human Ran:GTP and RanGAP from *S. pombe* is estimated to be around 150 $s^{-1}$, while our measured $k_{cat}$ values range from 1 to 10 $s^{-1}$, as above, $\frac{K}{K_S} \ll 1$ the assumption of steady-state holds true as long as $[E_0]$ << $K_m$ and $[E_0]$ << $[S_0]$, which is the case as we used 1-5 nM GAP in all of our experiments.

##### Potential caveats associated with using the GAP (Rna1) from *S. pombe*

All of our GAP-mediated GTP hydrolysis kinetics experiments used the wild type and mutant Gsp1 from *S. cerevisiae*, but Rna1 GAP from *S. pombe*. We chose to use the Rna1 ortholog from *S. pombe* as *S. cerevisiae* Rna1 formed soluble aggregates after purification, and *S. pombe* Rna1 was the only RanGAP for which there was a structure in complex with Ran (PDB IDs: 1k5d and 1k5g). While there could be slight differences between the kinetic parameters of *S. pombe* and *S. cerevisiae* GAP Rna1 acting on Gsp1, we do not believe these differences would significantly affect our conclusions, based on the following considerations:

1.) **Sequence conservation between *S. cerevisiae* and *S. pombe* Rna1.** A sequence alignment between *S. cerevisiae*, *S. pombe,* and human GAP proteins shows that all but one interface core residue in the PDB file 1k5d is conserved in sequence between *S. cerevisiae* and *S. pombe* (**Supplementary File 1 Supplementary Fig. 12**). Overall, out of the 1290 Å$^2$ buried by *S. pombe* Rna1 upon interface formation with Ran (PDB ID: 1k5d), 997 Å$^2$ (77%) are buried by residues that are conserved in sequence between *S. pombe* and *S. cerevisiae*, and the sequence identity of the Rna1 interface with Ran/Gsp1 (including all residues that change solvent accessible surface are upon complex formation) overall is 71% (**Supplementary File 1 Table 1**).

2.) **Comparable kinetic parameters to the human Ran/RanGAP1 pair.** The kinetic parameters for our *S. cerevisiae* Gsp1 and *S. pombe* Rna1 GAP are comparable to the kinetic parameters for the human Ran and human RanGAP1 reported by Klebe *et al.*[6]. They estimate a $K_m$ of 0.45 μM and $k_{cat}$ of 2.1 $s^{-1}$ for Ran/RanGAP1 at 25˚C, while our values for the wild type *S. cerevisiae* Gsp1 and *S. pombe* Rna1 at 30˚C are a $K_m$ of 0.38 μM and $k_{cat}$ of 9.2 $s^{-1}$. In addition, it was shown that Rna1 from *S. pombe* can activate the hydrolysis in both human and *S. cerevisiae* Ran/Gsp1 with very similar observed rates of hydrolysis (Fig. 4a in Becker *et al*[7]).

3.) **Our conclusions are based on relative values between the wild-type Gsp1 and its point mutants.** Although we report the absolute values of the kinetics parameters, when we compare the kinetics parameters with the results from genetic interaction profile and AP-MS, we always use the relative parameters as compared to the wild type. Based on the sequence conservation and comparable kinetics described above, we expect the relative ordering of mutants to be similar as well. Importantly, we use the relative kinetic data to group our mutants into three classes. Even in the case of small quantitative differences caused by using the *S. pombe* instead of the *S. cerevisae* Rna1 GAP, we make the assumption that these differences would not significantly affect this grouping.

#### Supplementary Figures

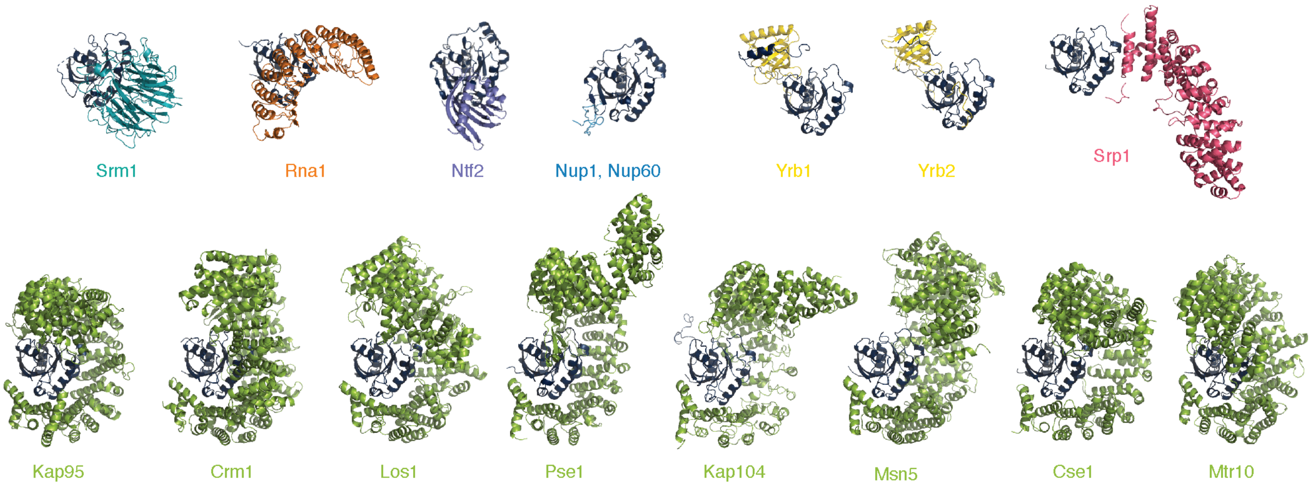


**Supplementary Figure 1 Cartoon representation of co-complex structures of *S. cerevisiae* Gsp1 (dark navy) with indicated partners (or homologs)**. Srm1 (PDB: 1i2m), Rna1 (PDB: 1k5d), Ntf2 (PDB: 1a2k), Nup1/Nup60 (PDB: 3ch5), Yrb1 (PDB: 3m1i), Yrb2 (PDB: 3wyf), Srp1 (PDB: 1wa5), Kap95 (PDB: 2bku), Crm1 (PDB: 3m1i), Los1 (PDB: 3icq), Pse1(PDB: 3w3z), Kap104 (PDB: 1qbk), Msn5 (PDB: 3a6p), Cse1 (PDB: 1wa5), Mtr10 (PDB: 4ol0). Species and sequence identity to *S. cerevisiae* homologs for these structures are provided in **Supplementary File 1 Table 1**

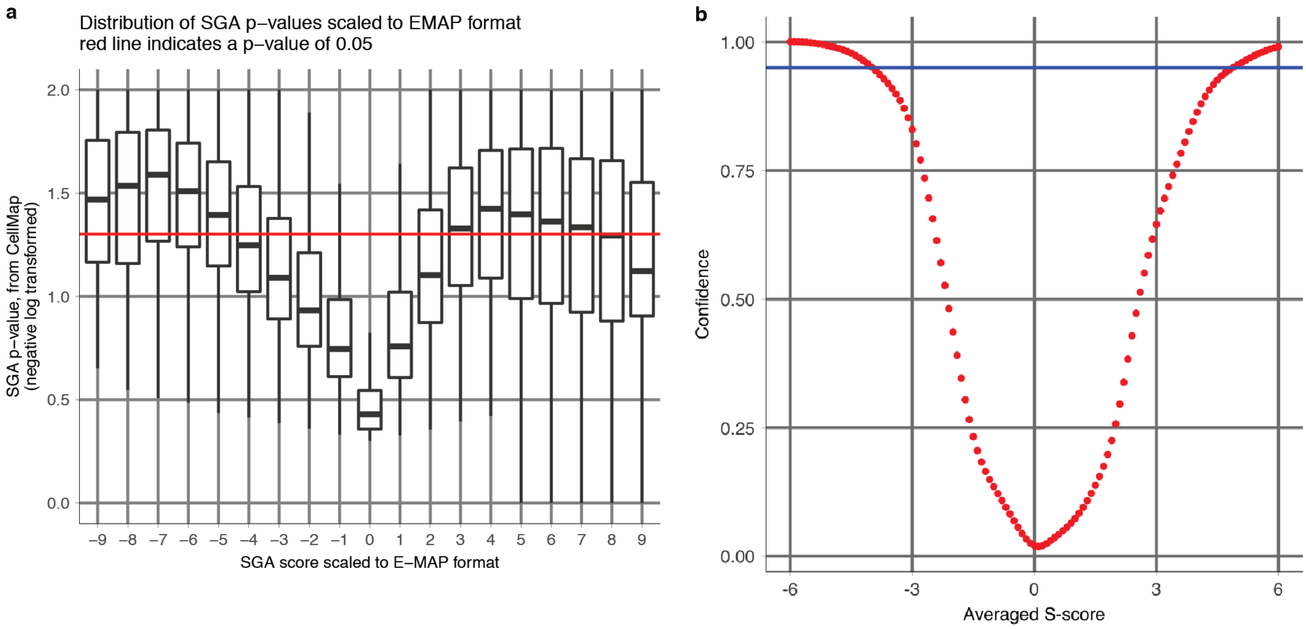


**Supplementary Figure 2 Comparison of definitions of high confidence S-scores used in our analysis**. **a,** Distribution of the SGA scores scaled to the E-MAP S-scores versus their corresponding published p-values from the CellMap[8]. **b,** Distribution of the E-MAP S-score averaged from all the individual replicates versus the confidence of the functional genetic interaction reproduced from Collins *et al*[9].

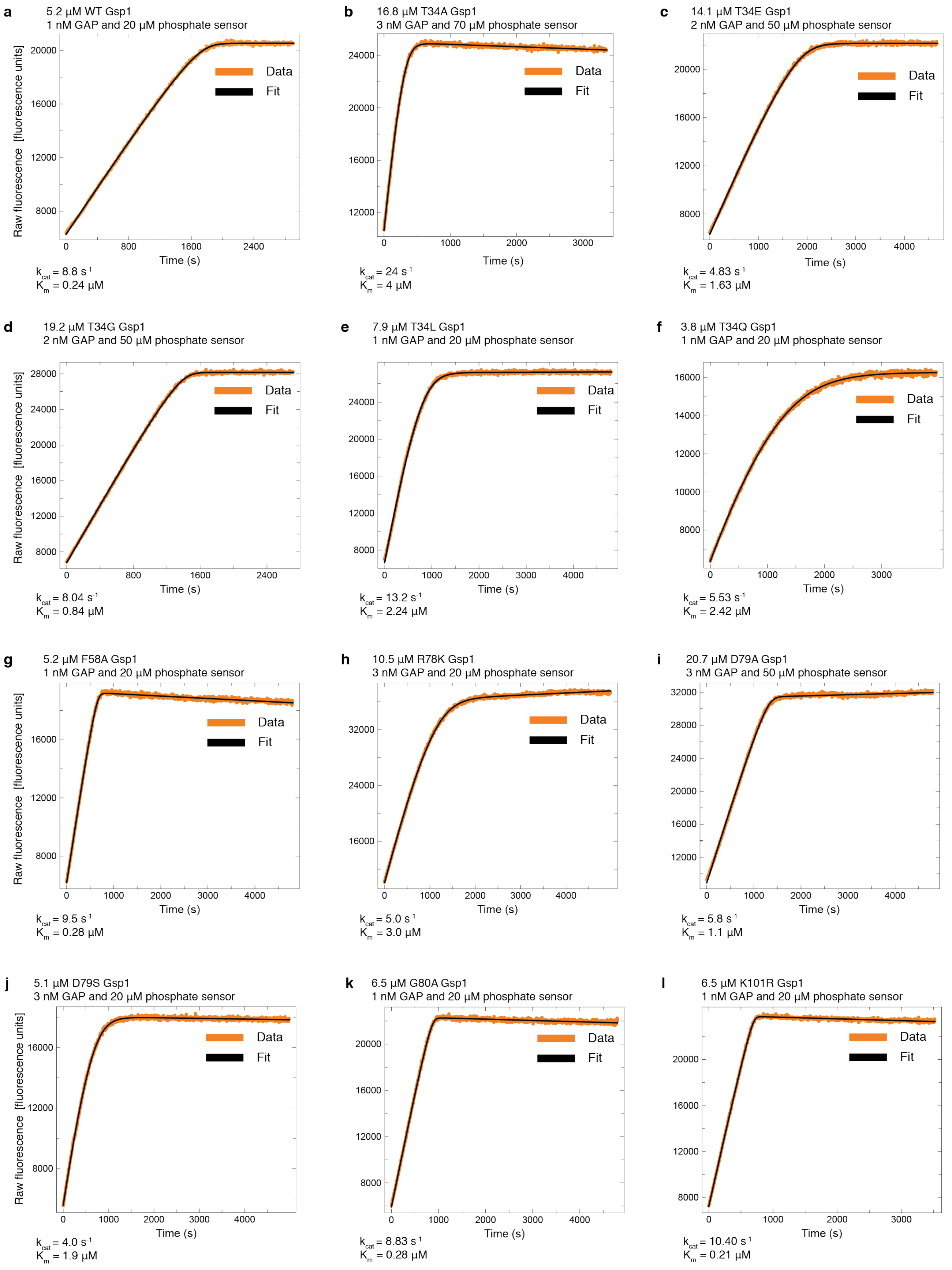
a
5.2 µM WT Gsp1
1 nM GAP and 20 µM phosphate sensor
Data
Fit
Raw fluorescence [fluorescence units]
Time (s)
$k_{cat}$ = 8.8 $s^{-1}$
$K_m$ = 0.24 µM
b
16.8 µM T34A Gsp1
3 nM GAP and 70 µM phosphate sensor
Data
Fit
Time (s)
$k_{cat}$ = 24 $s^{-1}$
$K_m$ = 4 µM
c
14.1 µM T34E Gsp1
2 nM GAP and 50 µM phosphate sensor
Data
Fit
Time (s)
$k_{cat}$ = 4.83 $s^{-1}$
$K_m$ = 1.63 µM
d
19.2 µM T34G Gsp1
2 nM GAP and 50 µM phosphate sensor
Data
Fit
Raw fluorescence [fluorescence units]
Time (s)
$k_{cat}$ = 8.04 $s^{-1}$
$K_m$ = 0.84 µM
e
7.9 µM T34L Gsp1
1 nM GAP and 20 µM phosphate sensor
Data
Fit
Time (s)
$k_{cat}$ = 13.2 $s^{-1}$
$K_m$ = 2.24 µM
f
3.8 µM T34Q Gsp1
1 nM GAP and 20 µM phosphate sensor
Data
Fit
Time (s)
$k_{cat}$ = 5.53 $s^{-1}$
$K_m$ = 2.42 µM
g
5.2 µM F58A Gsp1
1 nM GAP and 20 µM phosphate sensor
Data
Fit
Raw fluorescence [fluorescence units]
Time (s)
$k_{cat}$ = 9.5 $s^{-1}$
$K_m$ = 0.28 µM
h
10.5 µM R78K Gsp1
3 nM GAP and 20 µM phosphate sensor
Data
Fit
Time (s)
$k_{cat}$ = 5.0 $s^{-1}$
$K_m$ = 3.0 µM
i
20.7 µM D79A Gsp1
3 nM GAP and 50 µM phosphate sensor
Data
Fit
Time (s)
$k_{cat}$ = 5.8 $s^{-1}$
$K_m$ = 1.1 µM
j
5.1 µM D79S Gsp1
3 nM GAP and 20 µM phosphate sensor
Data
Fit
Raw fluorescence [fluorescence units]
Time (s)
$k_{cat}$ = 4.0 $s^{-1}$
$K_m$ = 1.9 µM
k
6.5 µM G80A Gsp1
1 nM GAP and 20 µM phosphate sensor
Data
Fit
Time (s)
$k_{cat}$ = 8.83 $s^{-1}$
$K_m$ = 0.28 µM
l
6.5 µM K101R Gsp1
1 nM GAP and 20 µM phosphate sensor
Data
Fit
Time (s)
$k_{cat}$ = 10.40 $s^{-1}$
$K_m$ = 0.21 µM

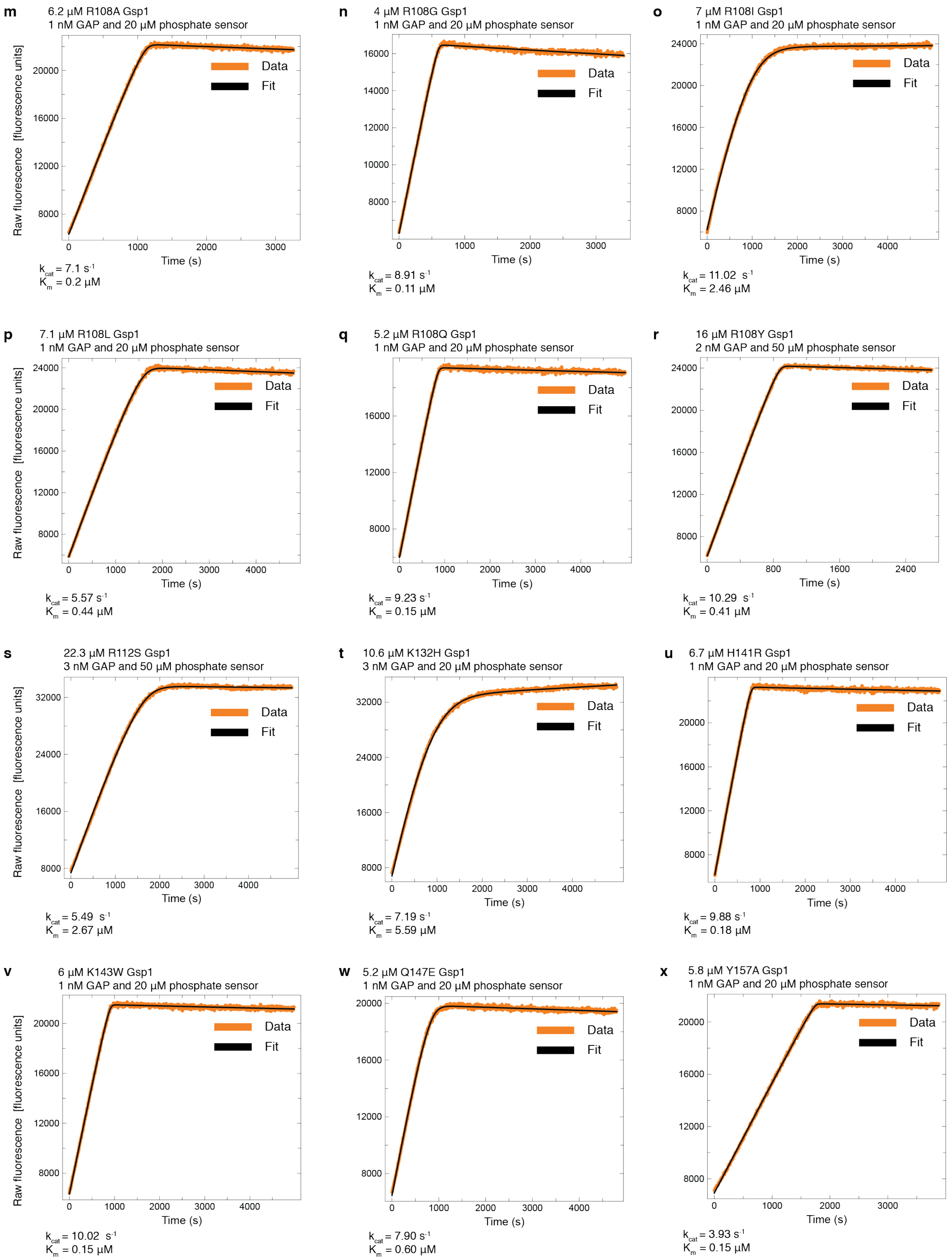
m
6.2 μM R108A Gsp1
1 nM GAP and 20 μM phosphate sensor
Raw fluorescence [fluorescence units]
8000 12000 16000 20000
0 1000 2000 3000
Time (s)
Data
Fit
$k_{cat}$ = 7.1 $s^{-1}$
$K_m$ = 0.2 μM
n
4 μM R108G Gsp1
1 nM GAP and 20 μM phosphate sensor
8000 10000 12000 14000 16000
0 1000 2000 3000
Time (s)
Data
Fit
$k_{cat}$ = 8.91 $s^{-1}$
$K_m$ = 0.11 μM
o
7 μM R108I Gsp1
1 nM GAP and 20 μM phosphate sensor
8000 12000 16000 20000 24000
0 1000 2000 3000 4000
Time (s)
Data
Fit
$k_{cat}$ = 11.02 $s^{-1}$
$K_m$ = 2.46 μM
p
7.1 μM R108L Gsp1
1 nM GAP and 20 μM phosphate sensor
Raw fluorescence [fluorescence units]
8000 12000 16000 20000 24000
0 1000 2000 3000 4000
Time (s)
Data
Fit
$k_{cat}$ = 5.57 $s^{-1}$
$K_m$ = 0.44 μM
q
5.2 μM R108Q Gsp1
1 nM GAP and 20 μM phosphate sensor
8000 12000 16000
0 1000 2000 3000 4000
Time (s)
Data
Fit
$k_{cat}$ = 9.23 $s^{-1}$
$K_m$ = 0.15 μM
r
16 μM R108Y Gsp1
2 nM GAP and 50 μM phosphate sensor
8000 12000 16000 20000 24000
0 800 1600 2400
Time (s)
Data
Fit
$k_{cat}$ = 10.29 $s^{-1}$
$K_m$ = 0.41 μM
s
22.3 μM R112S Gsp1
3 nM GAP and 50 μM phosphate sensor
Raw fluorescence [fluorescence units]
8000 16000 24000 32000
0 1000 2000 3000 4000
Time (s)
Data
Fit
$k_{cat}$ = 5.49 $s^{-1}$
$K_m$ = 2.67 μM
t
10.6 μM K132H Gsp1
3 nM GAP and 20 μM phosphate sensor
8000 16000 24000 32000
0 1000 2000 3000 4000
Time (s)
Data
Fit
$k_{cat}$ = 7.19 $s^{-1}$
$K_m$ = 5.59 μM
u
6.7 μM H141R Gsp1
1 nM GAP and 20 μM phosphate sensor
8000 12000 16000 20000
0 1000 2000 3000 4000
Time (s)
Data
Fit
$k_{cat}$ = 9.88 $s^{-1}$
$K_m$ = 0.18 μM
v
6 μM K143W Gsp1
1 nM GAP and 20 μM phosphate sensor
Raw fluorescence [fluorescence units]
8000 12000 16000 20000
0 1000 2000 3000 4000
Time (s)
Data
Fit
$k_{cat}$ = 10.02 $s^{-1}$
$K_m$ = 0.15 μM
w
5.2 μM Q147E Gsp1
1 nM GAP and 20 μM phosphate sensor
8000 12000 16000 20000
0 1000 2000 3000 4000
Time (s)
Data
Fit
$k_{cat}$ = 7.90 $s^{-1}$
$K_m$ = 0.60 μM
x
5.8 μM Y157A Gsp1
1 nM GAP and 20 μM phosphate sensor
8000 12000 16000 20000
0 1000 2000 3000
Time (s)
Data
Fit
$k_{cat}$ = 3.93 $s^{-1}$
$K_m$ = 0.15 μM

**y**

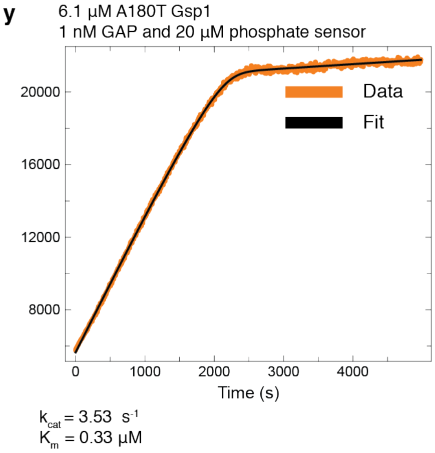


**Supplementary Figure 3 GAP-mediated GTP hydrolysis monitored as fluorescence increase upon binding of released free phosphate to a fluorescent phosphate sensor.** Curves were fit with the integrated Michaelis-Menten equation using the DELA software. Final Michaelis-Menten kinetic parameters ($k_{cat}$ and $K_m$) for each Gsp1 mutant were calculated from three to nine individually fit curves as the ones shown in this figure. **a,** Wild type Gsp1, **b-y,** Gsp1 point mutants.

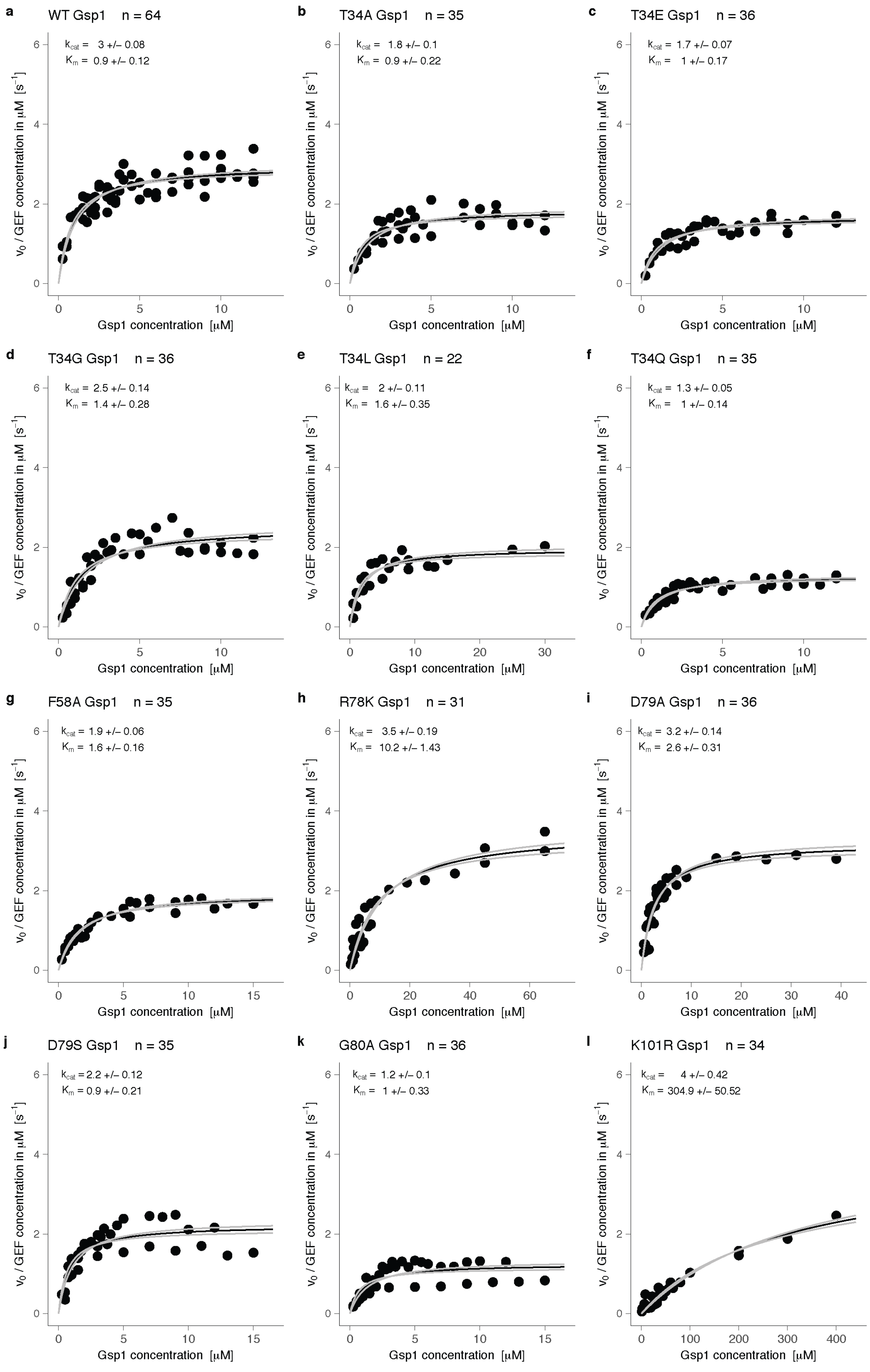
a
WT Gsp1 n = 64
kcat = 3 +/– 0.08
Km = 0.9 +/– 0.12
b
T34A Gsp1 n = 35
kcat = 1.8 +/– 0.1
Km = 0.9 +/– 0.22
c
T34E Gsp1 n = 36
kcat = 1.7 +/– 0.07
Km = 1 +/– 0.17
d
T34G Gsp1 n = 36
kcat = 2.5 +/– 0.14
Km = 1.4 +/– 0.28
e
T34L Gsp1 n = 22
kcat = 2 +/– 0.11
Km = 1.6 +/– 0.35
f
T34Q Gsp1 n = 35
kcat = 1.3 +/– 0.05
Km = 1 +/– 0.14
g
F58A Gsp1 n = 35
kcat = 1.9 +/– 0.06
Km = 1.6 +/– 0.16
h
R78K Gsp1 n = 31
kcat = 3.5 +/– 0.19
Km = 10.2 +/– 1.43
i
D79A Gsp1 n = 36
kcat = 3.2 +/– 0.14
Km = 2.6 +/– 0.31
j
D79S Gsp1 n = 35
kcat = 2.2 +/– 0.12
Km = 0.9 +/– 0.21
k
G80A Gsp1 n = 36
kcat = 1.2 +/– 0.1
Km = 1 +/– 0.33
l
K101R Gsp1 n = 34
kcat = 4 +/– 0.42
Km = 304.9 +/– 50.52
v0 / GEF concentration in μM [s−1]
Gsp1 concentration [μM]

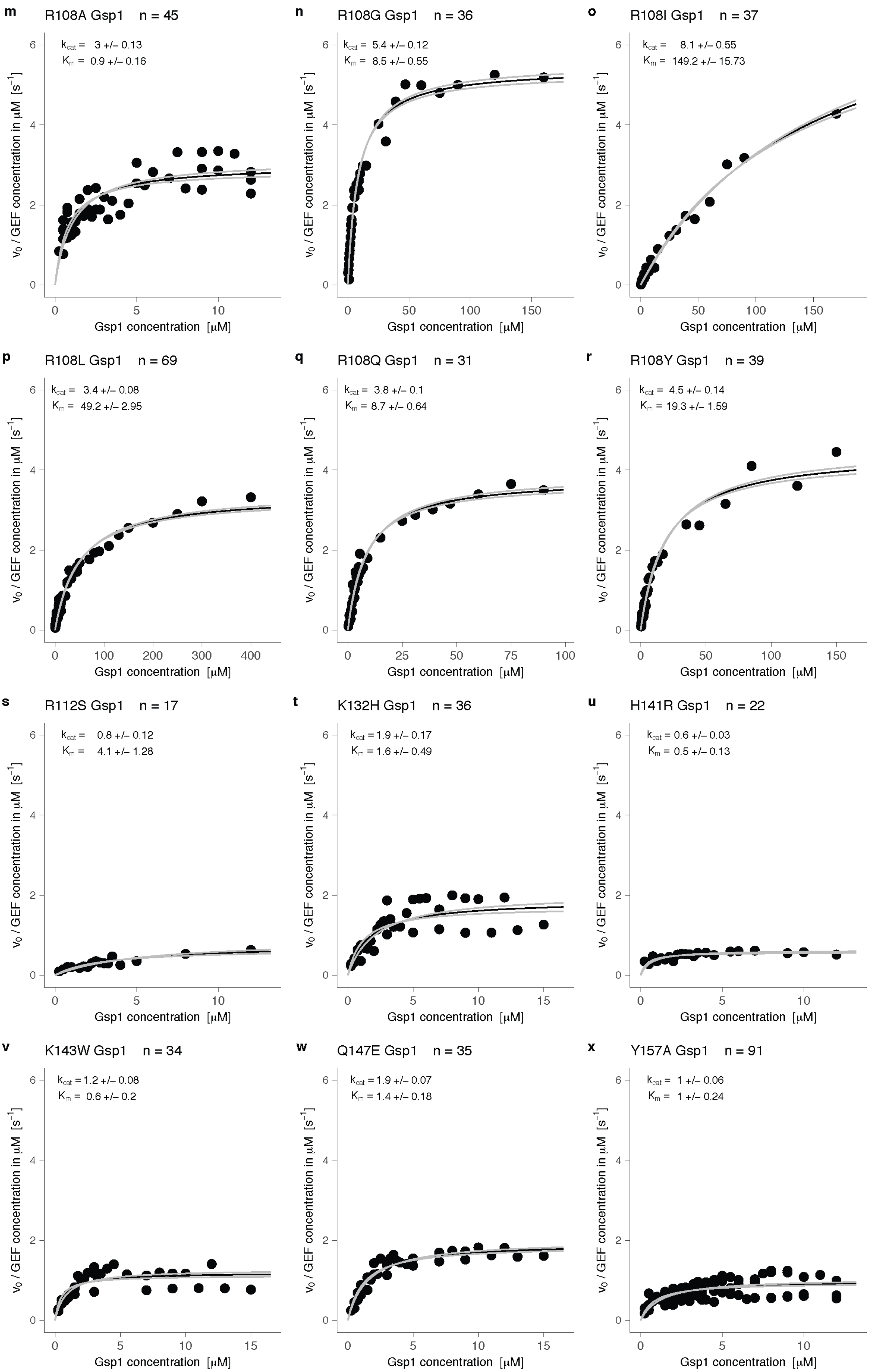

m
R108A Gsp1 n = 45
$k_{cat}$ = 3 +/– 0.13
$K_m$ = 0.9 +/– 0.16
n
R108G Gsp1 n = 36
$k_{cat}$ = 5.4 +/– 0.12
$K_m$ = 8.5 +/– 0.55
o
R108I Gsp1 n = 37
$k_{cat}$ = 8.1 +/– 0.55
$K_m$ = 149.2 +/– 15.73
p
R108L Gsp1 n = 69
$k_{cat}$ = 3.4 +/– 0.08
$K_m$ = 49.2 +/– 2.95
q
R108Q Gsp1 n = 31
$k_{cat}$ = 3.8 +/– 0.1
$K_m$ = 8.7 +/– 0.64
r
R108Y Gsp1 n = 39
$k_{cat}$ = 4.5 +/– 0.14
$K_m$ = 19.3 +/– 1.59
s
R112S Gsp1 n = 17
$k_{cat}$ = 0.8 +/– 0.12
$K_m$ = 4.1 +/– 1.28
t
K132H Gsp1 n = 36
$k_{cat}$ = 1.9 +/– 0.17
$K_m$ = 1.6 +/– 0.49
u
H141R Gsp1 n = 22
$k_{cat}$ = 0.6 +/– 0.03
$K_m$ = 0.5 +/– 0.13
v
K143W Gsp1 n = 34
$k_{cat}$ = 1.2 +/– 0.08
$K_m$ = 0.6 +/– 0.2
w
Q147E Gsp1 n = 35
$k_{cat}$ = 1.9 +/– 0.07
$K_m$ = 1.4 +/– 0.18
x
Y157A Gsp1 n = 91
$k_{cat}$ = 1 +/– 0.06
$K_m$ = 1 +/– 0.24
$v_0$ / GEF concentration in μM [$s^{-1}$]
Gsp1 concentration [μM]

**y**

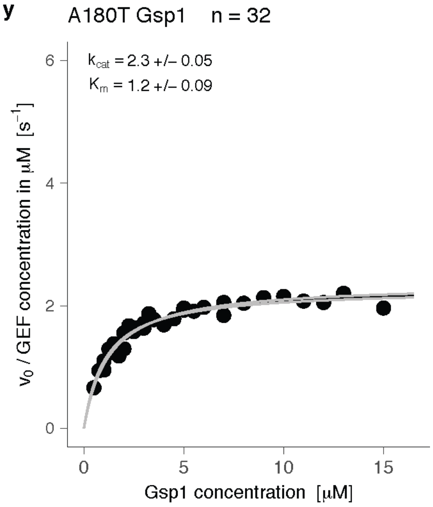


**Supplementary Figure 4 Michaelis-Menten plots for GEF-mediated nucleotide exchange**. Black line represents the Michaelis-Menten fit, and the gray lines represent the plus and minus one standard error of the fit. **a,** Wild type Gsp1. **b-y**, Gsp1 point mutants.

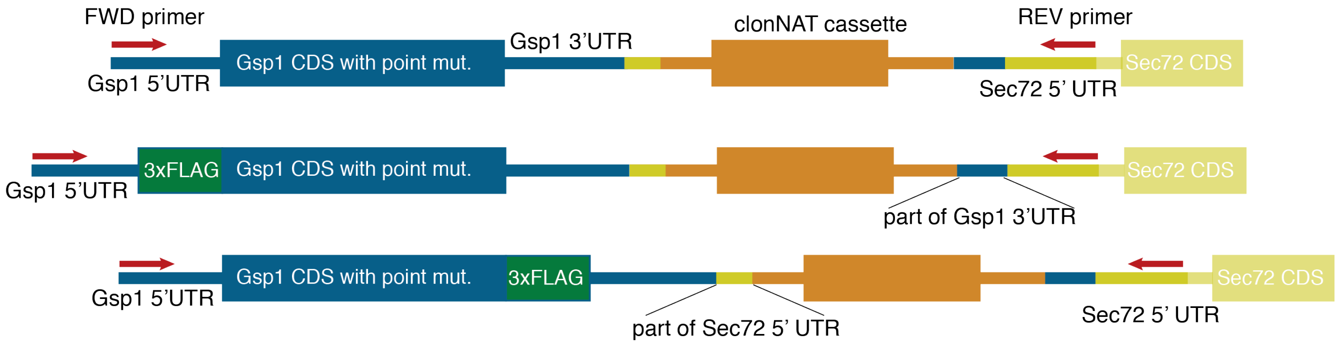


**Supplementary Figure 5 Schematic of genomically integrated *GSP1* constructs**. For E-MAP experiments, wild type or mutant GSP1 cassettes including the clonNAT resistance cassette were integrated into the MAT:α strain. For AP-MS the constructs also included either an amino- (N) terminal or a carboxy- (C) terminal 3xFLAG tag (M**DYKDHDGDYKDHDIDYKDDDDK**GGGGA and GGGGA**DYKDHDGDYKDHDIDYKDDDDK**, respectively).

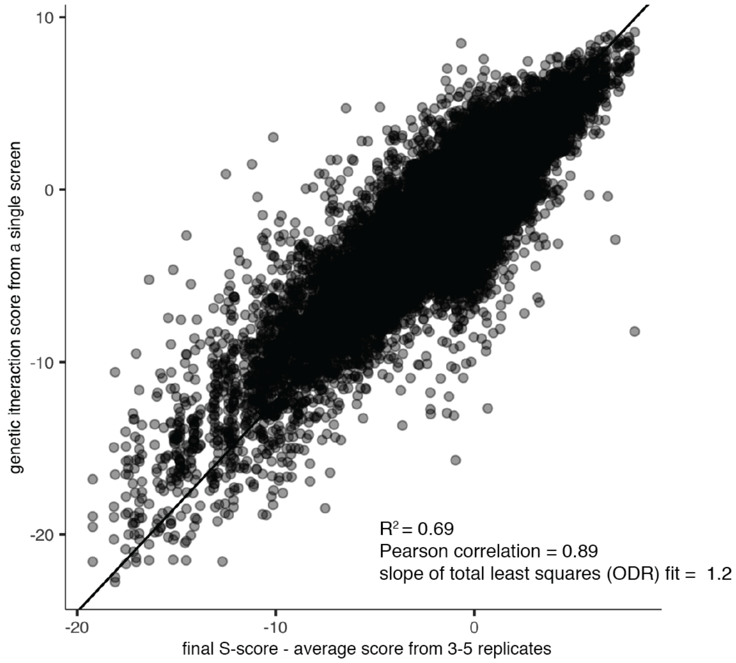


**Supplementary Figure 6 Reproducibility of *GSP1* point mutant E-MAP screens**, represented as a linear relationship between the genetic interaction S-score from a single E-MAP experiment and the final average S-score based on three or more replicates. The linear fit was calculated using the odregress function from the pracma R package.

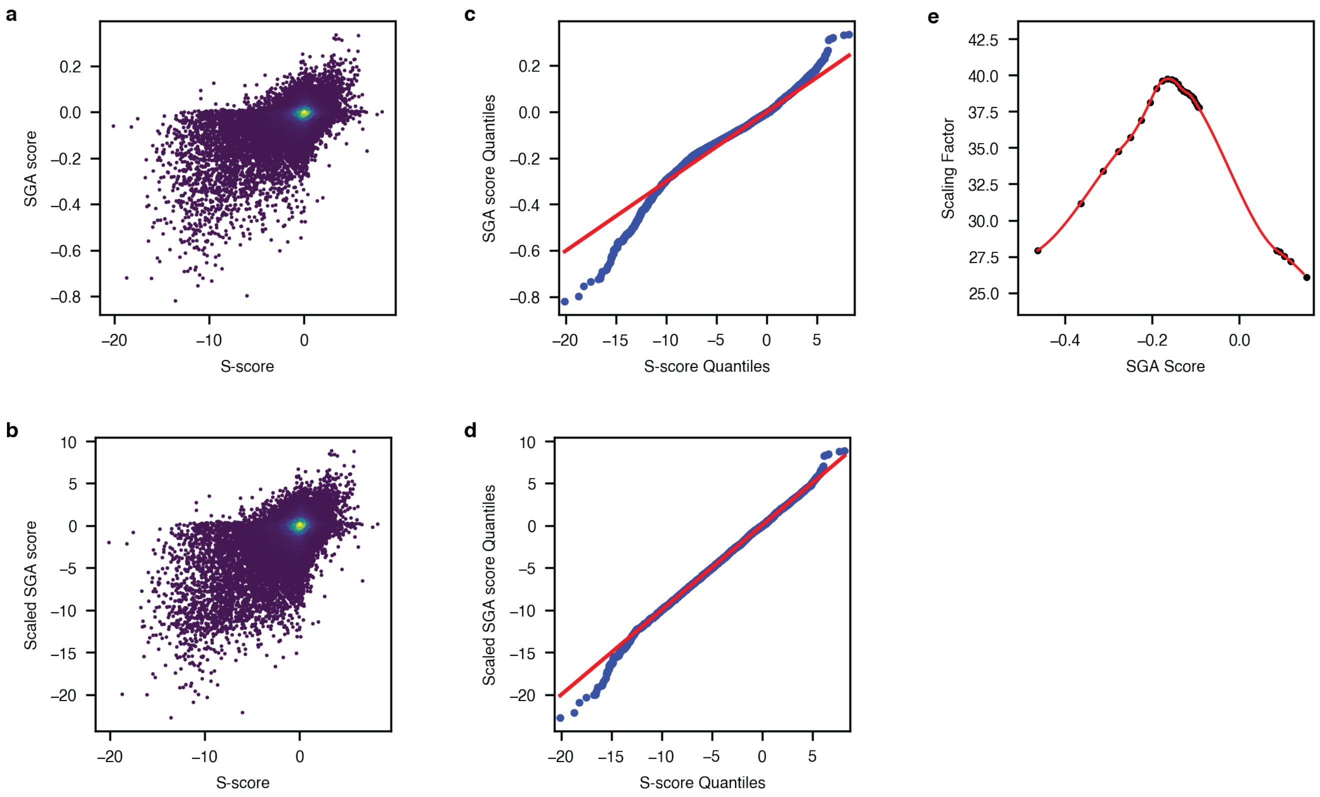


**Supplementary Figure 7 Non-linear scaling of SGA data from the Cell Map[8] to E-MAP format. a,** Distribution of S-scores from the chromatin biology E-MAP dataset[10] and the SGA score from the CellMap dataset. **b,** Distribution of S-scores from the chromatin biology E-MAP dataset and the *scaled* SGA score from the CellMap dataset. **c,** Quantile-quantile plot showing the distribution of genetic interaction scores from the CellMap and E-MAP chromatin biology datasets. **d,** Quantile-quantile plot showing the distribution of genetic interaction scores from the CellMap and E-MAP chromatin biology datasets after the CellMap dataset was scaled. **e,** The scaling function applied to the CellMap data. Red curve is the fitted spline of the scaling factors between the E-MAP S-scores and the SGA scores. Black dots represent the individual bins.

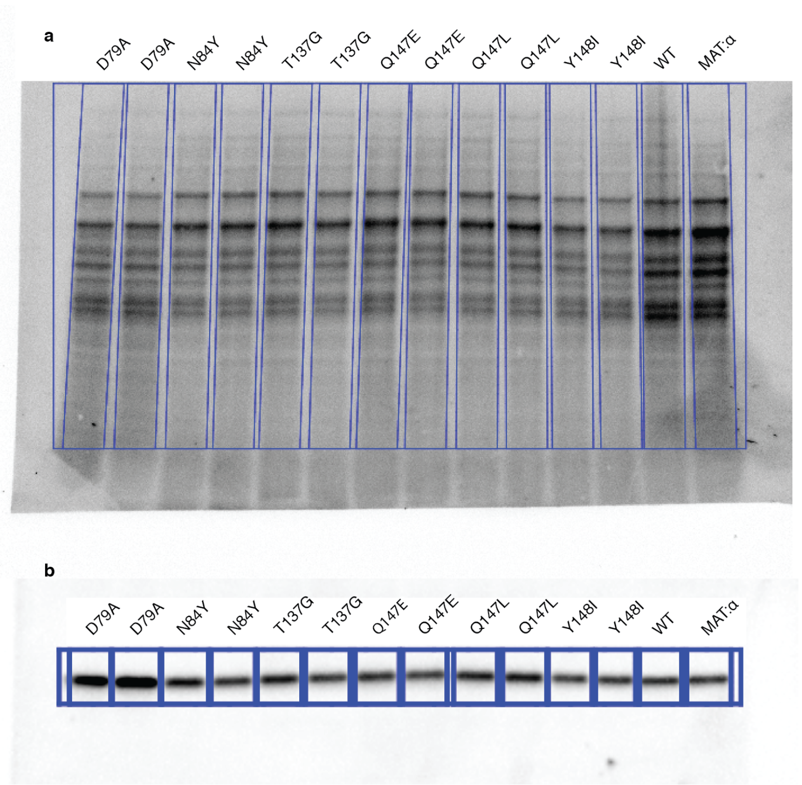


**Supplementary Figure 8 Example data for Gsp1 protein expression estimation by Western blot. a**, Total protein staining. **b**, Western blot of starting *S. cerevisiae* strain (MAT:α, see **Methods** for full strain description), wild type Gsp1 with clonNAT resistance marker (WT), and its mutants with anti-Ran antibody.

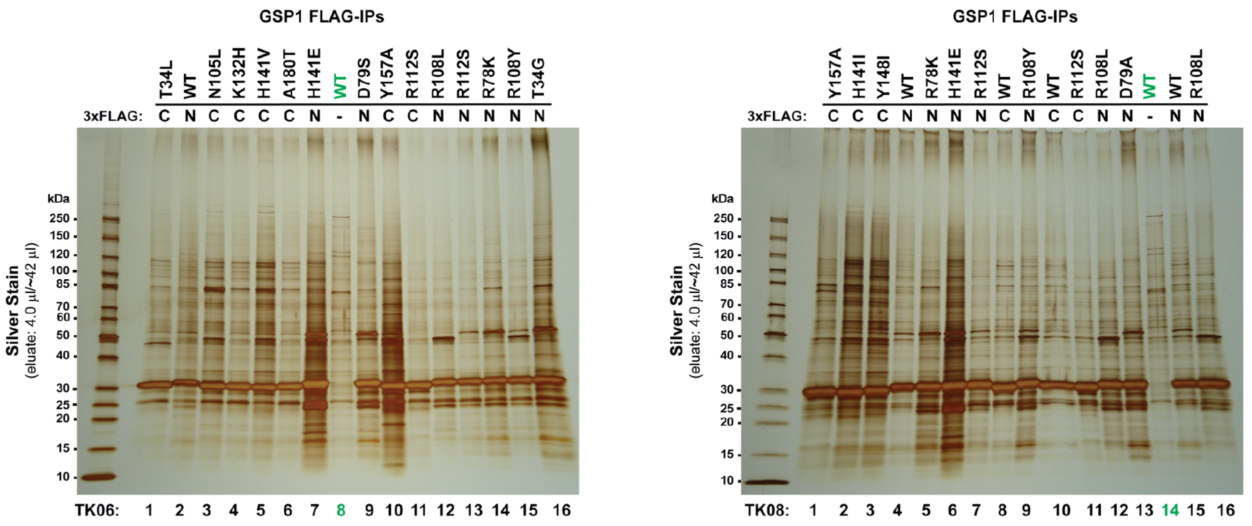


**Supplementary Figure 9 Silver stain gels after FLAG immunoprecipitation of amino- (N) or carboxy- (C) terminally 3xFLAG tagged genomically integrated Gsp1.** The strongest band at approximatelly 30 kDa corresponds to tagged Gsp1. Untagged wild type Gsp1 (lanes 8 and 14 in the left and right gel, respectively) were used as negative control for mass spectrometry analysis.

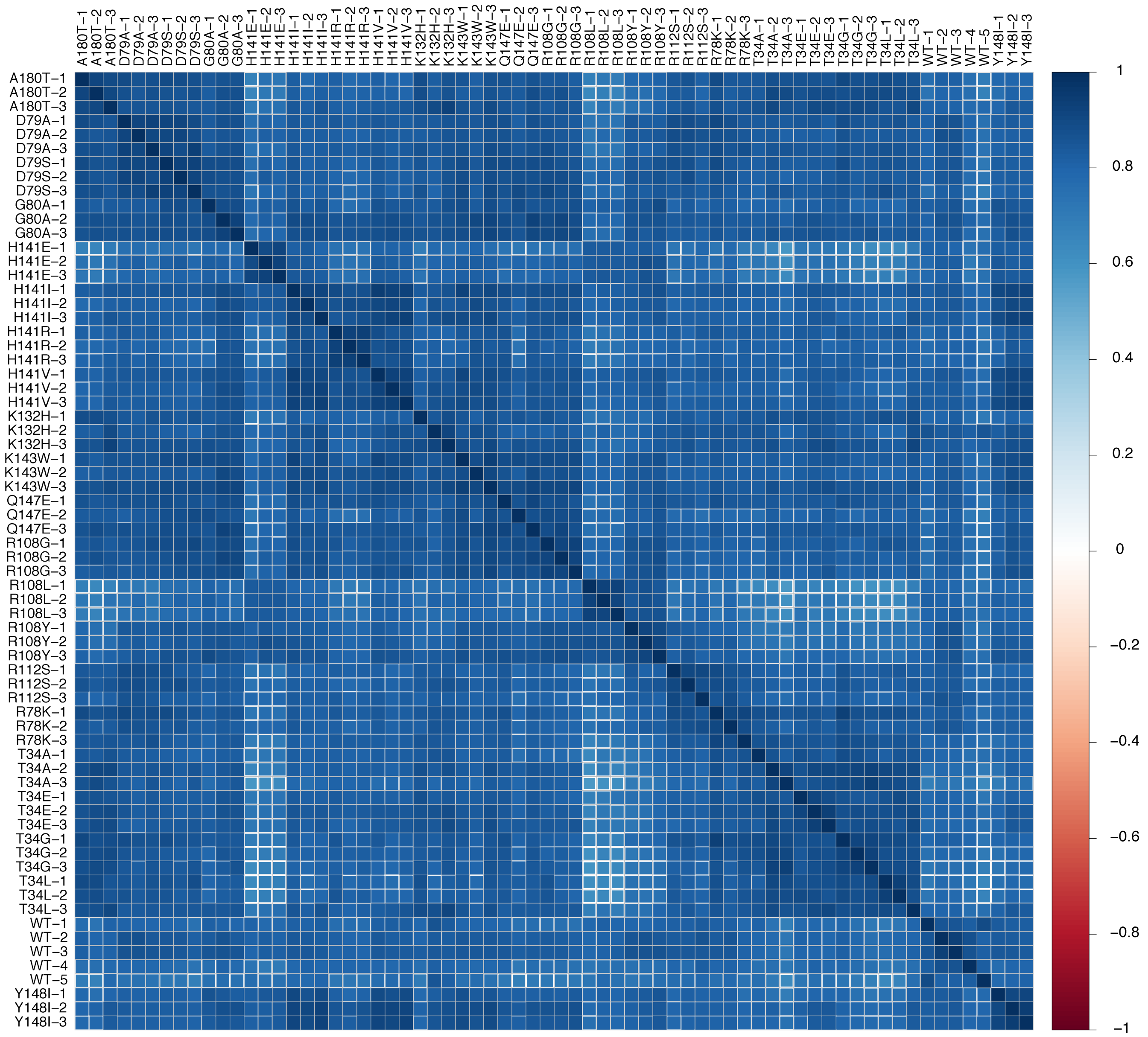


**Supplementary Figure 10 Clustering of individual AP-MS replicates based on correlations between protein abundance before the final scoring.** Data shown are for amino-terminally FLAG tagged wild type (WT) and Gsp1 mutants.

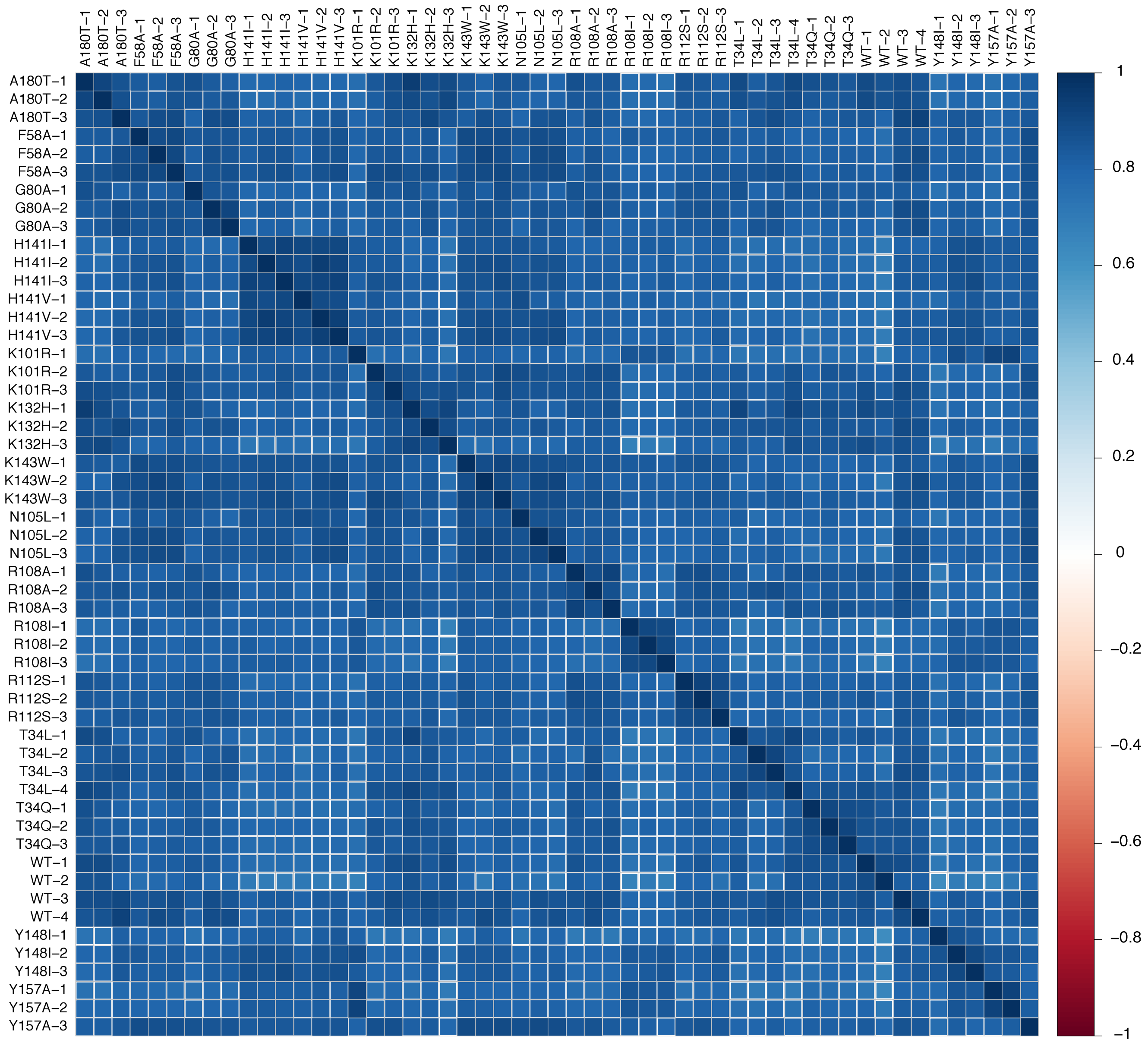


**Supplementary Figure 11 Clustering of individual AP-MS replicates based on correlations between protein abundance before the final scoring.** Data show are for carboxy-terminally FLAG tagged wild type (WT) and Gsp1 mutants.

```
Rna1_YEAST    82 ADLYTSRLVDEVVDSLKFLLPVLLK-CPHLEIVNLSDNAFGLRTI 125
Rna1_SCHPO    68 SDIFTGRVKDEIPEALRLLLQALLK-CPKLHTVRLSDNAFGPTAQ 111
RAGP1_HUMAN   85 SDMFTGRLRTEIPPALISLGEGLITAGAQLVELDLSDNAFGPDGV 129

Rna1_YEAST   126 ELLEDYIAH--AVNIKHLILSNNGMGPFAGERIGKALFHLAQNKK 168
Rna1_SCHPO   112 EPLIDFLSK--HTPLEHLYLHNNGLGPQAGAKIARALQELAVNKK 154
RAGP1_HUMAN  130 QGFEALLKSSACFTLQELKLNNCGMGIGGGKILAAALTECHRKSS 174

Rna1_YEAST   169 AASKP-FLETFICGRNRLENGSAVYLALGLKSHSEGLKVVKLYQN 212
Rna1_SCHPO   155 AKNAP-PLRSIICGRNRLENGSMKEWAKTFQSHR-LLHTVKMVQN 197
RAGP1_HUMAN  175 AQGKPLALKVFVAGRNRLENDGATALAEAFRVIG-TLEEVHMPQN 218

Rna1_YEAST   213 GIRPKGVATLIHYGLQYLKNLEILDLQDNTFTKHASLILAKALPT 257
Rna1_SCHPO   198 GIRPEGIEHLLLEGLAYCQELKVLDLQDNTFTHLGSSALAIALKS 242
RAGP1_HUMAN  219 GINHPGITALA-QAFAVNPLLRVINLNDNTFTEKGAVAMAETLKT 262

Rna1_YEAST   258 WKDSLFELNLNDCLLKTAGSDEVFKVFTEVKFPNLHVLKFEYNEM 302
Rna1_SCHPO   243 WPN-LRELGLNDCLLSARGAAAVVDAFSKLENIGLQTLRLQYNEI 286
RAGP1_HUMAN  263 LRQ-VEVINFGDCLVRSKGAVAIADAIR-GGLPKLKELNLSFCEI 305
```

**Supplementary Figure 12 Multiple sequence alignment between Rna1 from *S. cerevisiae* (*Rna1_YEAST*) and *S. pombe* (*Rna1_SCHPO*), as well as human RanGAP (*RAGP1_HUMAN*,** excluding the C-terminal SUMO conjugation domain which is absent in Fungi). Overall sequence identity between *S. cerevisiae* and *S. pombe* Rna1 is 39%, with 53% sequence similarity. Interface core residues (based on the X-ray crystal structure between *S. pombe* Rna1 and mammalian Ran, PDB ID: 1k5d) are highlighted in orange. All interface core residues except Pro108 in *S. pombe* Rna1, which corresponds to Leu122 in *S. cerevisiae* Rna1, are conserved in sequence between *S. cerevisiae* and *S. pombe* Rna1.

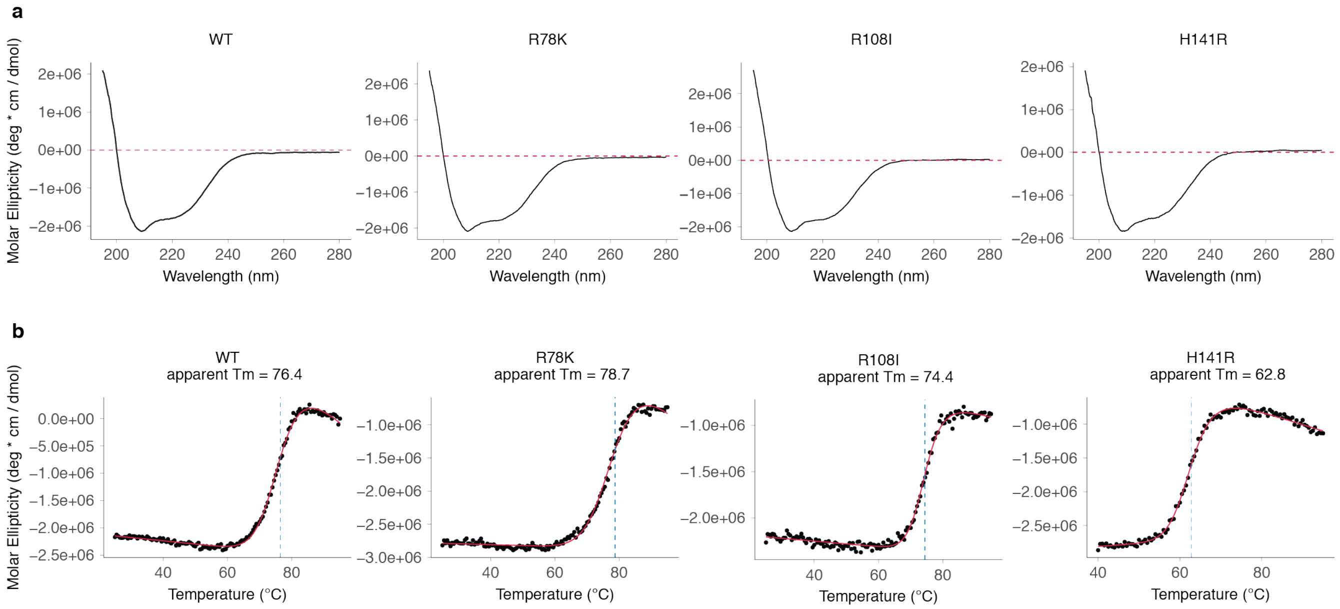


**Supplementary Figure 13 Circular dichroism (CD) data** for wild type (WT) Gsp1 and select mutants. **a,** CD spectra. **b,** Irreversible temperature melts.

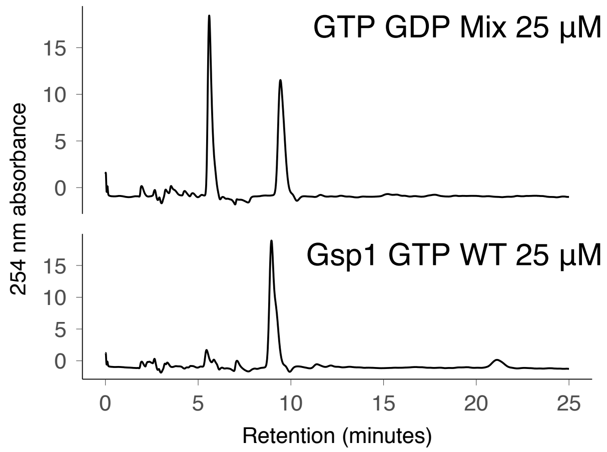


**Supplementary Figure 14 HPLC reverse phase chromatograms of a GTP/GDP mix (top) and that of a purified and GTP loaded wild type Gsp1 (bottom).**

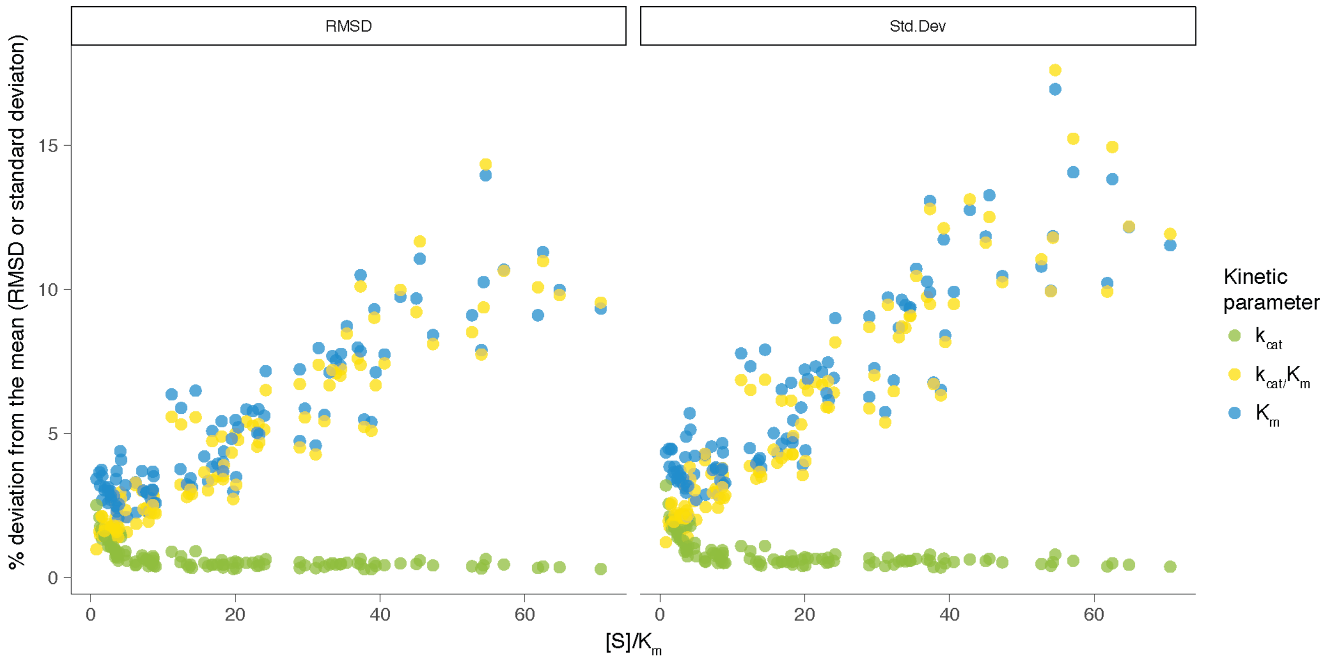


**Supplementary Figure 15 Accuracy estimation for determining the kinetic parameters of GAP-mediated GTP hydrolysis from individual time courses spanning [S] > $K_m$ to [S] << $K_m$ fit with an accurate solution of the integrated Michaelis Menten (IMM) equation.** Each time course was simulated using the experimentally determined parameters determined from the fitted IMM model, with added Gaussian noise similar to the experimental fluorescence signal noise. The deviation from the mean is plotted against a ratio of initial substrate (Gsp1:GTP) concentration [S] and the experimentally determined $K_m$. Deviation from the mean is reported either as standard deviation or $RMSD = \sqrt{\frac{\sum(\text{simulation\_param} - \text{experimental\_param})^2}{N}}$, where N = 100 simulations, and simulation_param and experimental_param are experimental and simulated $k_{cat}$, $K_m$, and $k_{cat}/K_m$, respectively. Here, simulated refers to the average of the fitted values for the simulated data sets.

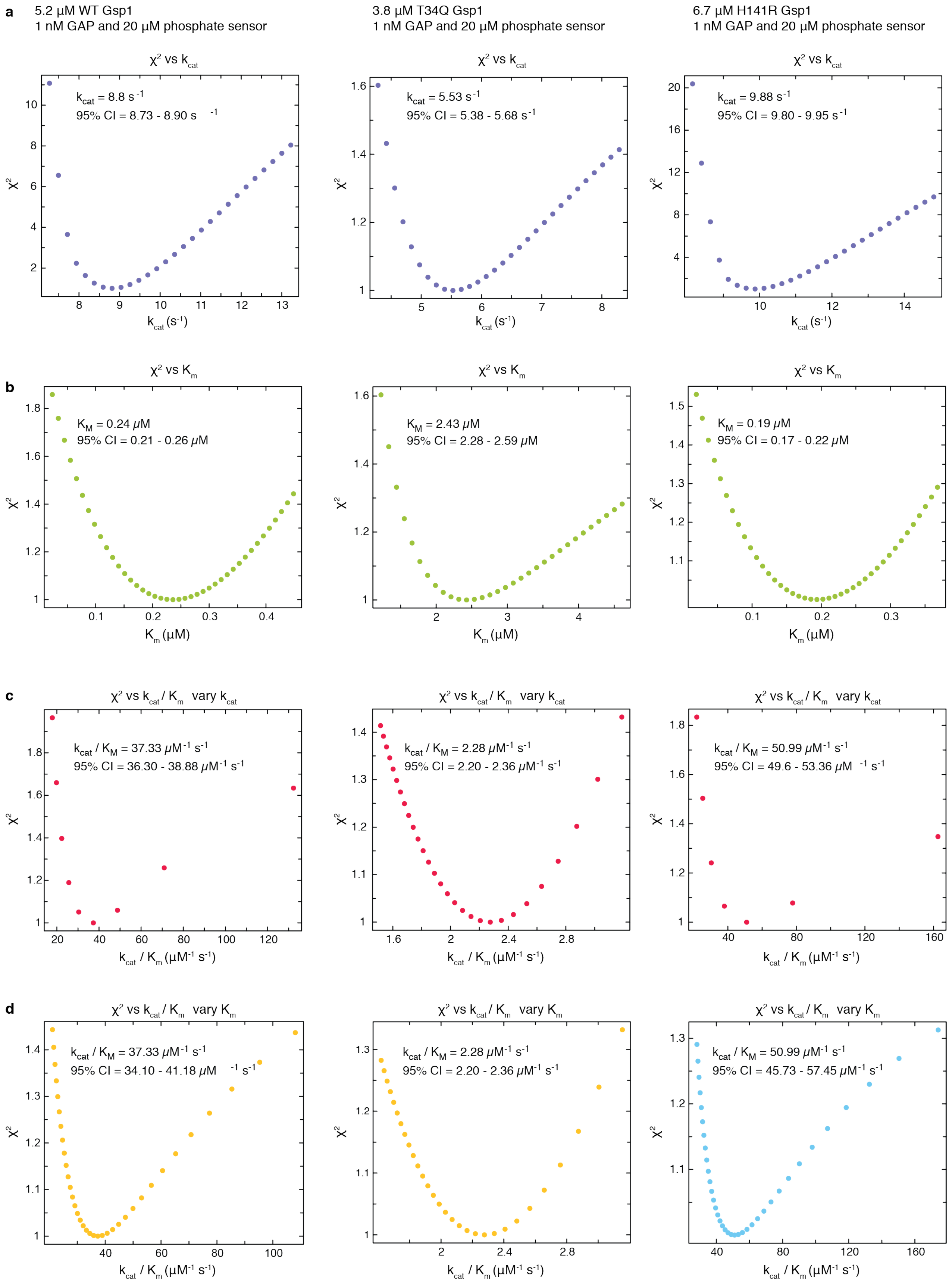


**Supplementary Figure 16 Estimated error around the maximum likelihood estimated values of the Michaelis-Menten parameters.** Plotted is the change in $\chi^2$ statistics as each of the parameters was fixed in gradual increments around the maximum likelihood value. The $\chi^2$ values are relative to the maximum likelihood values.

Error estimate analysis is shown for three of the Gsp1 variants: wild type Gsp1, the low efficiency Gsp1 T34Q mutant, and the high efficiency Gsp1 H141R mutant. 95% CI is the estimated 95% confidence interval for each value, based on the $\chi^2$ surface. **a**, Change of $\chi^2$ statistics as the $k_{cat}$ value is varied around the maximum likelihood value. **b**, Change of $\chi^2$ statistics as the $K_m$ value is varied around the maximum likelihood value. **c**, Change of $\chi^2$ statistics as the $k_{cat}/K_m$ value is varied around the maximum likelihood value and the $K_m$ is kept fixed at the maximum likelihood value ($k_{cat}$ is varied). **d**, Change of $\chi^2$ statistics as the $k_{cat}/K_m$ value is varied around the maximum likelihood value and the $k_{cat}$ is kept fixed at the maximum likelihood value ($K_m$ is varied).

### Supplementary Tables

**Supplementary File 1 Table 1** Co-complex X-ray crystal structures of Ran or Gsp1 with its partners.

| Ran/Gsp1 binding **partner** | | | | Ran/Gsp1 source species | Overall sequence identity to *S. cerevisiae* homolog | | Interface sequence identity to *S. cerevisiae* homolog | |
|---|---|---|---|---|---|---|---|---|
| Gene name | Partner protein name / function | PDB ID | source species | | Gsp1 [%] | partner [%] | Gsp1 [%] | partner [%] |
| Srm1 | Guanine nucleotide exchange factor of Gsp1 (GEF) | 1i2m | *H. sapiens* | *H. sapiens* | 83 | 25 | 94 | 42 |
| Rna1 | Ran GTPase-activating protein 1 of Gsp1 (GAP) | 1k5d | *S. pombe* | *H. sapiens* | 83 | 39 | 84 | 71 |
| Ntf2 | Nuclear transport factor 2 | 1a2k | *R. norvegicus* | *C. lupus* | 83 | 40 | 89 | 44 |
| Nup1 | FG-repeat nucleoporin | 3ch5 | *R. norvegicus* | *H. sapiens* | 83 | 13 | 67 | 37 |
| Nup60 | FG-repeat nucleoporin | 3ch5 | *R. norvegicus* | *H. sapiens* | 83 | 8 | 67 | 37 |
| Yrb1 | Ran-specific GTPase-activating protein 1 | 3m1i | *S. cerevisiae* | *S. cerevisiae* | 100 | 100 | 100 | 100 |
| Yrb2 | Ran-specific GTPase-activating protein 2 | 3wyf | *S. cerevisiae* | *S. cerevisiae* | 100 | 100 | 100 | 100 |
| Srp1 | Importin subunit alpha - receptor for simple and bipartite NLS | 1wa5 | *S. cerevisiae* | *C. lupus* | 83 | 100 | 67 | 100 |
| Kap95 | Importin subunit beta-1 - receptor for cNLS | 2bku | *S. cerevisiae* | *C. lupus* | 83 | 100 | 94 | 100 |
| Crm1 | Exportin-1 - Receptor for the leucine-rich nuclear export signal (NES) | 3m1i | *S. cerevisiae* | *S. cerevisiae* | 100 | 100 | 100 | 100 |
| Los1 | Exportin-T - tRNA nucleus export | 3icq | *S. pombe* | *S. cerevisiae* | 100 | 22 | 100 | 30 |
| Pse1 | Importin subunit beta-3 - receptor for cNLS and rg-NLS | 3w3z | *S. cerevisiae* | *C. lupus* | 83 | 100 | 89 | 100 |
| Kap104 | Importin subunit beta-2 - receptor for rg-NLS and PY-NLS | 1qbk | *H. sapiens* | *H. sapiens* | 83 | 31 | 92 | 43 |
| Msn5 | Exportin and importin of proteins and tRNA | 3a6p | *H. sapiens* | *C. lupus* | 83 | 18 | 89 | 29 |
| Cse1 | Importin alpha re-exporter - export receptor for Srp1 | 1wa5 | *S. cerevisiae* | *C. lupus* | 83 | 100 | 89 | 100 |
| Mtr10 | mRNA transport regulator | 4ol0 | *H. sapiens* | *H. sapiens* | 83 | 21 | 88 | 36 |

**Supplementary Table 2** Mutated residues in Gsp1 and their interface position and ΔrASA. CellMap alleles are annotated in parentheses.

| Gsp1 residue number | Crm1 | Cse1 (cse1-5002) | Kap104 | Kap95 (kap95-e126k) | Los1 (los1) | Msn5 (msn5) |
|---|---|---|---|---|---|---|
| 34 | | | | rim / 0.1 | | |
| 58 | | | | | | |
| 78 | rim / 0.1 | core / 0.34 | core / 0.44 | core / 0.2 | rim / 0.33 | rim / 0.18 |
| 79 | core / 0.3 | core / 0.29 | support / 0.12 | core / 0.37 | support / 0.16 | core / 0.28 |
| 80 | core / 0.31 | core / 0.27 | core / 0.42 | core / 0.29 | core / 0.37 | core / 0.32 |
| 84 | rim / 0.3 | rim / 0.21 | rim / 0.3 | rim / 0.41 | rim / 0.09 | rim / 0.09 |
| 101 | rim / 0.17 | | | | rim / 0.13 | rim / 0.02 |
| 102 | support / 0.01 | | | | core / 0.08 | |
| 105 | rim / 0.06 | | | rim / 0.03 | rim / 0.16 | core / 0.25 |
| 108 | core / 0.26 | rim / 0.1 | rim / 0.11 | rim / 0.12 | core / 0.43 | core / 0.42 |
| 112 | core / 0.55 | core / 0.45 | core / 0.44 | core / 0.56 | core / 0.4 | core / 0.58 |
| 115 | rim / 0.25 | rim / 0.2 | rim / 0.27 | rim / 0.07 | rim / 0.34 | rim / 0.34 |
| 129 | | rim / 0.61 | rim / 0.59 | rim / 0.19 | | rim / 0.23 |
| 132 | core / 0.12 | rim / 0 | | | rim / 0.03 | rim / 0.12 |
| 137 | | | | | | |
| 139 | rim / 0.01 | | | core / 0.04 | rim / 0.02 | |
| 141 | support / 0.14 | | support / 0.19 | core / 0.15 | | |
| 143 | rim / 0.48 | rim / 0.15 | core / 0.35 | rim / 0.27 | rim / 0.09 | rim / 0.01 |
| 147 | core / 0.23 | | support / 0.23 | core / 0.25 | core / 0.07 | core / 0.14 |
| 148 | support / 0.11 | support / 0.03 | support / 0.13 | | support / 0.01 | |
| 154 | | core / 0.28 | | core / 0.38 | rim / 0.13 | |
| 157 | core / 0.38 | rim / 0.13 | core / 0.39 | core / 0.29 | rim / 0.05 | |
| 169 | rim / 0.21 | | | rim / 0.02 | | rim / 0.17 |
| 180 | | | rim / 0.01 | | | |
| **Gsp1 residue number** | **Mtr10** | **Ntf2 (ntf2-h104y, ntf2-5001)** | **Nup1** | **Nup60** | **Pse1** | **Rna1 (rna1-1, rna1-s116f)** |
| 34 | | | | | | |
| 58 | | | rim / 0.28 | rim / 0.28 | | |
| 78 | core / 0.25 | core / 0.57 | rim / 0 | rim / 0 | support / 0.03 | support / 0.02 |
| 79 | core / 0.36 | rim / 0.1 | | | core / 0.26 | |
| 80 | core / 0.51 | core / 0.27 | support / 0.13 | support / 0.13 | core / 0.51 | |
| 84 | rim / 0.36 | | rim / 0.25 | rim / 0.25 | rim / 0.2 | |
| 101 | | | | | | rim / 0.01 |
| 102 | support / 0.1 | | | | | support / 0.07 |
| 105 | core / 0.21 | | | | | |
| 108 | core / 0.18 | | | | rim / 0.12 | |
| 112 | core / 0.57 | | | | core / 0.46 | |
| 115 | rim / 0.12 | | | | rim / 0.37 | |
| 129 | | | | | | rim / 0 |
| 132 | | | | | | core / 0.44 |
| 137 | core / 0.08 | | | | | rim / 0.01 |
| 139 | rim / 0.15 | | | | core / 0.18 | |
| 141 | core / 0.14 | | | | support / 0 | |
| 143 | core / 0.44 | | | | core / 0.52 | |
| 147 | support / 0.09 | | | | core / 0.09 | |
| 148 | support / 0.01 | | | | | |
| 154 | rim / 0.03 | | | | rim / 0.05 | |
| 157 | rim / 0.04 | | | | | |
| 169 | | | | | | |
| 180 | | | | | | |

| Gsp1 residue number | Srm1 (srm1-g282s, srm1-ts) | Srp1 (srp1-5001 | Yrb1 (yrb1-51) | Yrb2 |
|---|---|---|---|---|
| 34 | | | core / 0.4 | rim / 0.24 |
| 58 | | | core / 0.4 | core / 0.39 |
| 78 | rim / 0.46 | | | |
| 79 | rim / 0.01 | | | |
| 80 | | | | |
| 84 | | | | |
| 101 | core / 0.67 | core / 0.47 | | |
| 102 | support / 0.15 | | | |
| 105 | core / 0.44 | rim / 0.03 | | |
| 108 | core / 0.47 | | | |
| 112 | rim / 0.24 | | | |
| 115 | | | | |
| 129 | | rim / 0.1 | | |
| 132 | rim / 0.16 | core / 0.22 | | |
| 137 | | core / 0.2 | | |
| 139 | core / 0.26 | rim / 0.01 | | |
| 141 | | | | |
| 143 | rim / 0.07 | | | |
| 147 | | | | |
| 148 | | | | |
| 154 | | | | |
| 157 | | | | |
| 169 | | | | |
| 180 | | | core / 0.64 | rim / 0.5 |

**Supplementary Table 3** - Gsp1 mutants and attempted yeast constructs

| construct name | Gsp1 residue number | Gsp1 point mutation | yeast strain successfully made |
|---|---|---|---|
| C-terminal 3xFLAG GSP1 T34L | 34 | T34L | yes |
| C-terminal 3xFLAG GSP1 T34Q | 34 | T34Q | yes |
| GSP1 T34A | 34 | T34A | yes |
| GSP1 T34D | 34 | T34D | yes |
| GSP1 T34E | 34 | T34E | yes |
| GSP1 T34G | 34 | T34G | yes |
| GSP1 T34L | 34 | T34L | yes |
| GSP1 T34Q | 34 | T34Q | yes |
| GSP1 T34S | 34 | T34S | yes |
| GSP1 T34Y | 34 | T34Y | yes |
| N-terminal 3xFLAG GSP1 T34A | 34 | T34A | yes |
| N-terminal 3xFLAG GSP1 T34E | 34 | T34E | yes |
| N-terminal 3xFLAG GSP1 T34G | 34 | T34G | yes |
| N-terminal 3xFLAG GSP1 T34L | 34 | T34L | yes |
| C-terminal 3xFLAG GSP1 F58A | 58 | F58A | yes |
| GSP1 F58A | 58 | F58A | yes |
| GSP1 F58L | 58 | F58L | yes |
| GSP1 R78K | 78 | R78K | yes |
| N-terminal 3xFLAG GSP1 R78K | 78 | R78K | yes |
| C-terminal 3xFLAG GSP1 D79A | 79 | D79A | yes |
| GSP1 D79A | 79 | D79A | yes |
| GSP1 D79S | 79 | D79S | yes |
| N-terminal 3xFLAG GSP1 D79A | 79 | D79A | yes |
| N-terminal 3xFLAG GSP1 D79S | 79 | D79S | yes |
| C-terminal 3xFLAG GSP1 G80A | 80 | G80A | yes |
| GSP1 G80A | 80 | G80A | yes |
| N-terminal 3xFLAG GSP1 G80A | 80 | G80A | yes |
| GSP1 N84Y | 84 | N84Y | yes |
| C-terminal 3xFLAG GSP1 K101R | 101 | K101R | yes |
| GSP1 K101R | 101 | K101R | yes |
| GSP1 N102I | 102 | N102I | yes |
| GSP1 N102K | 102 | N102K | yes |
| GSP1 N102M | 102 | N102M | yes |
| C-terminal 3xFLAG GSP1 N105L | 105 | N105L | yes |
| GSP1 N105L | 105 | N105L | yes |
| GSP1 N105V | 105 | N105V | yes |
| C-terminal 3xFLAG GSP1 R108A | 108 | R108A | yes |
| C-terminal 3xFLAG GSP1 R108I | 108 | R108I | yes |
| C-terminal 3xFLAG GSP1 R108Y | 108 | R108Y | yes |
| GSP1 R108A | 108 | R108A | yes |
| GSP1 R108D | 108 | R108D | yes |
| GSP1 R108G | 108 | R108G | yes |
| GSP1 R108I | 108 | R108I | yes |
| GSP1 R108L | 108 | R108L | yes |
| GSP1 R108Q | 108 | R108Q | yes |
| GSP1 R108S | 108 | R108S | yes |
| GSP1 R108Y | 108 | R108Y | yes |

| construct name | Gsp1 residue number | Gsp1 point mutation | yeast strain successfully made |
|---|---|---|---|
| N-terminal 3xFLAG GSP1 R108G | 108 | R108G | yes |
| N-terminal 3xFLAG GSP1 R108Y | 108 | R108Y | yes |
| C-terminal 3xFLAG GSP1 R112S | 112 | R112S | yes |
| GSP1 R112A | 112 | R112A | yes |
| GSP1 R112S | 112 | R112S | yes |
| N-terminal 3xFLAG GSP1 R112S | 112 | R112S | yes |
| GSP1 E115A | 115 | E115A | yes |
| GSP1 E115I | 115 | E115I | yes |
| GSP1 K129E | 129 | K129E | yes |
| GSP1 K129F | 129 | K129F | yes |
| GSP1 K129I | 129 | K129I | yes |
| GSP1 K129T | 129 | K129T | yes |
| C-terminal 3xFLAG GSP1 K132H | 132 | K132H | yes |
| GSP1 K132H | 132 | K132H | yes |
| N-terminal 3xFLAG GSP1 K132H | 132 | K132H | yes |
| GSP1 T137G | 137 | T137G | yes |
| GSP1 T139A | 139 | T139A | yes |
| GSP1 T139R | 139 | T139R | yes |
| C-terminal 3xFLAG GSP1 H141I | 141 | H141I | yes |
| C-terminal 3xFLAG GSP1 H141V | 141 | H141V | yes |
| GSP1 H141E | 141 | H141E | yes |
| GSP1 H141I | 141 | H141I | yes |
| GSP1 H141R | 141 | H141R | yes |
| GSP1 H141V | 141 | H141V | yes |
| N-terminal 3xFLAG GSP1 H141E | 141 | H141E | yes |
| N-terminal 3xFLAG GSP1 H141I | 141 | H141I | yes |
| N-terminal 3xFLAG GSP1 H141R | 141 | H141R | yes |
| N-terminal 3xFLAG GSP1 H141V | 141 | H141V | yes |
| C-terminal 3xFLAG GSP1 K143W | 143 | K143W | yes |
| GSP1 K143H | 143 | K143H | yes |
| GSP1 K143W | 143 | K143W | yes |
| GSP1 K143Y | 143 | K143Y | yes |
| N-terminal 3xFLAG GSP1 K143W | 143 | K143W | yes |
| GSP1 Q147E | 147 | Q147E | yes |
| GSP1 Q147L | 147 | Q147L | yes |
| N-terminal 3xFLAG GSP1 Q147E | 147 | Q147E | yes |
| C-terminal 3xFLAG GSP1 Y148I | 148 | Y148I | yes |
| GSP1 Y148I | 148 | Y148I | yes |
| N-terminal 3xFLAG GSP1 Y148I | 148 | Y148I | yes |
| GSP1 K154M | 154 | K154M | yes |
| C-terminal 3xFLAG GSP1 Y157A | 157 | Y157A | yes |
| GSP1 Y157A | 157 | Y157A | yes |
| GSP1 K169I | 169 | K169I | yes |
| C-terminal 3xFLAG GSP1 A180T | 180 | A180T | yes |
| GSP1 A180T | 180 | A180T | yes |
| N-terminal 3xFLAG GSP1 A180T | 180 | A180T | yes |
| T34A Cter3xFL | 34 | T34A | no |

| construct name | Gsp1 residue number | Gsp1 point mutation | yeast strain successfully made |
|---|---|---|---|
| T34E Cter3xFL | 34 | T34E | no |
| T34G Cter3xFL | 34 | T34G | no |
| T34Q Nter3xFL | 34 | T34Q | no |
| K39M | 39 | K39M | no |
| Y41A | 41 | Y41A | no |
| V49D | 49 | V49D | no |
| F58A Nter3xFL | 58 | F58A | no |
| G70N | 70 | G70N | no |
| Q71E | 71 | Q71E | no |
| K73Q | 73 | K73Q | no |
| G75N | 75 | G75N | no |
| R78K Cter3xFL | 78 | R78K | no |
| D79K | 79 | D79K | no |
| D79S Cter3xFL | 79 | D79S | no |
| G80N | 80 | G80N | no |
| G80S | 80 | G80S | no |
| I98F | 98 | I98F | no |
| K101R Nter3xFL | 101 | K101R | no |
| R108G Cter3xFL | 108 | R108G | no |
| R108I Nter3xFL | 108 | R108I | no |
| R108L Nter3xFL | 108 | R108L | no |
| R108Q Cter3xFL | 108 | R108Q | no |
| R108S Cter3xFL | 108 | R108S | no |
| K132M | 132 | K132M | no |
| K132Y | 132 | K132Y | no |
| T137E | 137 | T137E | no |
| H141E Cter3xFL | 141 | H141E | no |
| H141R Cter3xFL | 141 | H141R | no |
| Q147E Cter3xFL | 147 | Q147E | no |
| Y157K | 157 | Y157K | no |

**Supplementary Table 4** Pearson correlations between Gsp1 mutants and the alleles of their direct interaction partners from the SGA CellMap. Ordered by correlation value.

| GSP1 mutant | Partner strain name | Pearson correlation coefficient | Residue in interface core |
|---|---|---|---|
| D79S | kap95-e126k | 0.4146 | TRUE |
| Y148I | crm1_damp | 0.4027 | FALSE |
| R108L | ntf2-h104y | 0.3827 | FALSE |
| R108G | crm1_damp | 0.3783 | TRUE |
| R108L | ntf2-5001 | 0.3612 | FALSE |
| R108Y | ntf2-h104y | 0.3612 | FALSE |
| G80A | kap95-e126k | 0.3545 | TRUE |
| R112A | ntf2-h104y | 0.3533 | FALSE |
| R108Y | crm1_damp | 0.3453 | TRUE |
| K101R | ntf2-h104y | 0.3389 | FALSE |
| R112S | ntf2-h104y | 0.3353 | FALSE |
| R108Q | crm1_damp | 0.3291 | TRUE |
| T34A | ntf2-5001 | 0.3231 | FALSE |
| Q147E | kap95-e126k | 0.323 | TRUE |
| H141R | crm1_damp | 0.3199 | FALSE |
| K101R | srm1-ts | 0.3197 | TRUE |
| T34E | srm1-ts | 0.3155 | FALSE |
| T34Q | ntf2-h104y | 0.3135 | FALSE |
| R112A | ntf2-5001 | 0.3117 | FALSE |
| T34E | ntf2-h104y | 0.3116 | FALSE |
| R108Y | ntf2-5001 | 0.3091 | FALSE |
| D79S | ntf2-h104y | 0.309 | FALSE |
| D79S | srm1-ts | 0.3085 | FALSE |
| R112S | ntf2-5001 | 0.3043 | FALSE |
| D79S | cse1-5002 | 0.3022 | TRUE |
| T34Q | srm1-ts | 0.3015 | FALSE |
| T34A | ntf2-h104y | 0.2946 | FALSE |
| H141R | ntf2-h104y | 0.2929 | FALSE |
| T34E | yrb1-51 | 0.2898 | TRUE |
| T34A | srm1-ts | 0.2881 | FALSE |
| T34E | kap95-e126k | 0.2813 | FALSE |
| T34A | kap95-e126k | 0.2791 | FALSE |
| R108L | srm1-ts | 0.2773 | TRUE |
| D79A | kap95-e126k | 0.2754 | TRUE |
| H141I | crm1_damp | 0.2706 | FALSE |
| T34Q | kap95-e126k | 0.2681 | FALSE |
| T34A | yrb1-51 | 0.2676 | TRUE |
| T34Q | ntf2-5001 | 0.2588 | FALSE |
| Y148I | ntf2-5001 | 0.2555 | FALSE |
| K101R | ntf2-5001 | 0.255 | FALSE |
| R108I | ntf2-5001 | 0.2544 | FALSE |
| D79A | srm1-ts | 0.251 | FALSE |
| D79S | yrb1-51 | 0.2502 | FALSE |
| H141I | ntf2-5001 | 0.2501 | FALSE |
| T34G | cse1-5002 | 0.2467 | FALSE |
| D79A | ntf2-h104y | 0.2459 | FALSE |
| K101R | kap95-e126k | 0.2449 | FALSE |
| T34Q | yrb1-51 | 0.241 | TRUE |
| G80A | ntf2-h104y | 0.2402 | TRUE |
| T34G | kap95-e126k | 0.2365 | FALSE |
| K101R | srm1-g282s | 0.2359 | TRUE |
| G80A | cse1-5002 | 0.2357 | TRUE |
| Y148I | ntf2-h104y | 0.2355 | FALSE |
| T34E | ntf2-5001 | 0.2354 | FALSE |
| G80A | yrb1-51 | 0.2354 | FALSE |
| D79S | srp1-5001 | 0.2343 | FALSE |
| T34G | srm1-ts | 0.2321 | FALSE |
| G80A | crm1_damp | 0.2317 | TRUE |
| H141R | ntf2-5001 | 0.2296 | FALSE |
| T34A | crm1_damp | 0.2291 | FALSE |
| R108I | ntf2-h104y | 0.2275 | FALSE |
| G80A | ntf2-5001 | 0.2275 | TRUE |
| T34E | srm1-g282s | 0.2249 | FALSE |
| R108Q | ntf2-h104y | 0.2245 | FALSE |
| G80A | srm1-ts | 0.2188 | FALSE |
| H141E | rna1-s116f | 0.2185 | FALSE |
| R108L | crm1_damp | 0.2176 | TRUE |
| D79S | ntf2-5001 | 0.2171 | FALSE |
| R108G | ntf2-h104y | 0.2171 | FALSE |
| Q147E | srm1-ts | 0.2142 | FALSE |
| R108G | ntf2-5001 | 0.2131 | FALSE |
| T34A | cse1-5002 | 0.2104 | FALSE |
| Q147E | ntf2-h104y | 0.2101 | FALSE |
| R108Y | srm1-ts | 0.208 | TRUE |
| R112A | crm1_damp | 0.2075 | TRUE |
| H141E | kap95-e126k | 0.2073 | FALSE |
| R108I | srm1-ts | 0.2059 | TRUE |
| D79A | cse1-5002 | 0.2047 | TRUE |
| Q147E | ntf2-5001 | 0.2044 | FALSE |
| T34Q | srm1-g282s | 0.2038 | FALSE |
| H141I | ntf2-h104y | 0.2025 | FALSE |
| T34E | cse1-5002 | 0.1994 | FALSE |
| R112S | srm1-ts | 0.1945 | FALSE |
| Q147E | yrb1-51 | 0.1942 | FALSE |
| R108Q | ntf2-5001 | 0.1936 | FALSE |
| H141I | srm1-ts | 0.1934 | FALSE |
| R112A | srm1-ts | 0.1929 | FALSE |
| K101R | yrb1-51 | 0.1912 | FALSE |
| R108Q | yrb2_damp | 0.1895 | FALSE |
| R112S | crm1_damp | 0.1894 | TRUE |
| H141I | kap95-e126k | 0.188 | FALSE |
| H141E | cse1-5002 | 0.1817 | FALSE |
| T34E | srp1-5001 | 0.1805 | FALSE |
| T34G | yrb1-51 | 0.1781 | TRUE |
| T34G | srp1-5001 | 0.1779 | FALSE |
| G80A | srp1-5001 | 0.1775 | FALSE |
| H141E | ntf2-5001 | 0.1762 | FALSE |
| R108L | kap95-e126k | 0.1753 | FALSE |
| T34Q | cse1-5002 | 0.1738 | FALSE |
| T34A | srm1-g282s | 0.1729 | FALSE |

| GSP1 mutant | Partner strain name | Pearson correlation coefficient | Residue in interface core |
|---|---|---|---|
| R108I | kap95-e126k | 0.1719 | FALSE |
| H141R | yrb2_damp | 0.1717 | FALSE |
| D79A | srp1-5001 | 0.171 | FALSE |
| H141E | ntf2-h104y | 0.1682 | FALSE |
| D79A | yrb1-51 | 0.1672 | FALSE |
| R108G | yrb2_damp | 0.1669 | FALSE |
| Y148I | yrb2_damp | 0.1654 | FALSE |
| D79S | srm1-g282s | 0.1652 | FALSE |
| R108Y | yrb2_damp | 0.165 | FALSE |
| R108Y | kap95-e126k | 0.1645 | FALSE |
| T34A | rna1-s116f | 0.1637 | FALSE |
| D79A | ntf2-5001 | 0.1621 | FALSE |
| H141R | srm1-ts | 0.1596 | FALSE |
| T34A | srp1-5001 | 0.1557 | FALSE |
| D79A | srm1-g282s | 0.1534 | FALSE |
| T34Q | srp1-5001 | 0.1529 | FALSE |
| H141E | rna1-1 | 0.1529 | FALSE |
| R108L | srm1-g282s | 0.1527 | TRUE |
| Q147E | srm1-g282s | 0.1524 | FALSE |
| Y157A | crm1_damp | 0.1495 | TRUE |
| Q147E | cse1-5002 | 0.148 | FALSE |
| T34G | srm1-g282s | 0.1476 | FALSE |
| R108G | srm1-ts | 0.1462 | TRUE |
| T34G | ntf2-h104y | 0.1453 | FALSE |
| H141E | srm1-ts | 0.1446 | FALSE |
| Q147E | srp1-5001 | 0.1417 | FALSE |
| R108I | srm1-g282s | 0.139 | TRUE |
| H141E | crm1_damp | 0.138 | FALSE |
| Y148I | kap95-e126k | 0.134 | FALSE |
| R112A | kap95-e126k | 0.1302 | TRUE |
| T34E | rna1-1 | 0.1295 | FALSE |
| Q147E | crm1_damp | 0.1269 | FALSE |
| H141I | yrb2_damp | 0.1254 | FALSE |
| D79A | rna1-s116f | 0.1242 | FALSE |
| R108I | crm1_damp | 0.1232 | TRUE |
| T34Q | rna1-s116f | 0.1232 | FALSE |
| R108Q | srm1-ts | 0.1214 | TRUE |
| T34A | rna1-1 | 0.1214 | FALSE |
| T34G | rna1-1 | 0.1199 | FALSE |
| R112S | yrb2_damp | 0.1175 | FALSE |
| R108I | yrb1-51 | 0.1171 | FALSE |
| Y157A | rna1-s116f | 0.1168 | FALSE |
| R108G | kap95-e126k | 0.1162 | FALSE |
| R112S | srm1-g282s | 0.1154 | FALSE |
| H141R | kap95-e126k | 0.115 | FALSE |
| K101R | srp1-5001 | 0.1149 | TRUE |
| Q147E | rna1-s116f | 0.1139 | FALSE |
| H141E | srp1-5001 | 0.1135 | FALSE |
| R112S | kap95-e126k | 0.1112 | TRUE |
| D79S | rna1-s116f | 0.1081 | FALSE |
| G80A | srm1-g282s | 0.1073 | FALSE |
| H141I | srm1-g282s | 0.1062 | FALSE |
| H141I | rna1-s116f | 0.1045 | FALSE |
| R108Y | srm1-g282s | 0.104 | TRUE |
| T34G | rna1-s116f | 0.1031 | FALSE |
| R112A | srm1-g282s | 0.1025 | FALSE |
| H141E | yrb1-51 | 0.1023 | FALSE |
| T34E | rna1-s116f | 0.1013 | FALSE |
| R112A | yrb2_damp | 0.0975 | FALSE |
| T34Q | rna1-1 | 0.0959 | FALSE |
| G80A | rna1-s116f | 0.0947 | FALSE |
| Y148I | srm1-ts | 0.0933 | FALSE |
| T34E | los1 | 0.0932 | FALSE |
| Y157A | kap95-e126k | 0.092 | TRUE |
| R108L | yrb1-51 | 0.0904 | FALSE |
| D79S | rna1-1 | 0.089 | FALSE |
| Y157A | rna1-1 | 0.0878 | FALSE |
| K101R | cse1-5002 | 0.0869 | FALSE |
| Y148I | rna1-s116f | 0.086 | FALSE |
| Y157A | yrb1-51 | 0.0843 | FALSE |
| H141E | yrb2_damp | 0.0828 | FALSE |
| H141I | yrb1-51 | 0.082 | FALSE |
| R108I | msn5 | 0.0815 | TRUE |
| H141I | cse1-5002 | 0.0795 | FALSE |
| D79S | crm1_damp | 0.0789 | TRUE |
| R108I | los1 | 0.075 | TRUE |
| T34Q | crm1_damp | 0.0745 | FALSE |
| G80A | rna1-1 | 0.0735 | FALSE |
| T34E | crm1_damp | 0.0732 | FALSE |
| R108I | cse1-5002 | 0.0721 | FALSE |
| K101R | los1 | 0.0704 | FALSE |
| H141I | rna1-1 | 0.0702 | FALSE |
| R108L | los1 | 0.0699 | TRUE |
| T34A | los1 | 0.0695 | FALSE |
| Q147E | rna1-1 | 0.0666 | FALSE |
| Y148I | yrb1-51 | 0.0651 | FALSE |
| Y157A | cse1-5002 | 0.0637 | FALSE |
| R108G | srm1-g282s | 0.0626 | TRUE |
| T34Q | los1 | 0.0616 | FALSE |
| R108L | msn5 | 0.0612 | TRUE |
| R108G | rna1-1 | 0.0609 | FALSE |
| Y157A | ntf2-5001 | 0.0609 | FALSE |
| R108Q | kap95-e126k | 0.0608 | FALSE |
| R108Q | msn5 | 0.0592 | TRUE |
| H141I | srp1-5001 | 0.059 | FALSE |
| R108G | msn5 | 0.0587 | TRUE |
| R108G | rna1-s116f | 0.0578 | FALSE |
| H141R | srm1-g282s | 0.0573 | FALSE |
| D79A | rna1-1 | 0.0562 | FALSE |
| G80A | yrb2_damp | 0.0522 | FALSE |
| R108I | srp1-5001 | 0.0516 | FALSE |
| D79S | los1 | 0.0504 | FALSE |
| Y148I | rna1-1 | 0.0501 | FALSE |
| Y148I | srm1-g282s | 0.0482 | FALSE |
| D79A | crm1_damp | 0.0482 | TRUE |
| R108L | srp1-5001 | 0.047 | FALSE |

| GSP1 mutant | Partner strain name | Pearson correlation coefficient | Residue in interface core |
|---|---|---|---|
| R108I | rna1-1 | 0.045 | FALSE |
| R112S | rna1-s116f | 0.044 | FALSE |
| Q147E | los1 | 0.0436 | FALSE |
| R112S | rna1-1 | 0.0432 | FALSE |
| R108Y | los1 | -0.0089 | TRUE |
| R78K | srp1-5001 | -0.0124 | FALSE |
| R108G | los1 | -0.0126 | TRUE |
| R78K | los1 | -0.0127 | FALSE |
| R112A | cse1-5002 | -0.013 | TRUE |
| Y157A | msn5 | -0.0139 | FALSE |
| R112S | los1 | -0.0142 | TRUE |
| D79A | msn5 | -0.0197 | TRUE |
| T34G | crm1_damp | -0.0216 | FALSE |
| H141R | rna1-1 | -0.0229 | FALSE |
| R78K | msn5 | -0.023 | FALSE |
| H141I | los1 | -0.0242 | FALSE |
| H141I | msn5 | -0.0244 | FALSE |
| Y148I | msn5 | -0.0253 | FALSE |
| K101R | msn5 | -0.0271 | FALSE |
| T34Q | yrb2_damp | -0.0272 | TRUE |
| R108G | srp1-5001 | -0.0277 | FALSE |
| R112S | cse1-5002 | -0.0303 | TRUE |
| R108Q | cse1-5002 | -0.0317 | FALSE |
| R108L | rna1-s116f | -0.032 | FALSE |
| Q147E | msn5 | -0.0322 | FALSE |
| R78K | ntf2-h104y | -0.0322 | TRUE |
| T34E | yrb2_damp | -0.033 | TRUE |
| H141R | msn5 | -0.0364 | FALSE |
| R108Q | rna1-s116f | -0.0378 | FALSE |
| T34Q | msn5 | -0.0382 | FALSE |
| Y157A | srm1-g282s | -0.039 | FALSE |
| R78K | ntf2-5001 | -0.0426 | TRUE |
| H141E | msn5 | -0.0441 | FALSE |
| D79S | msn5 | -0.0471 | TRUE |
| T34G | msn5 | -0.0479 | FALSE |
| R78K | kap95-e126k | -0.0485 | FALSE |
| H141R | los1 | -0.0521 | FALSE |
| R108Q | los1 | -0.0593 | TRUE |
| Y148I | los1 | -0.0601 | FALSE |
| K101R | yrb2_damp | -0.0651 | FALSE |
| R108Q | srp1-5001 | -0.0685 | FALSE |
| R108Q | yrb1-51 | -0.0696 | FALSE |
| R78K | cse1-5002 | -0.0696 | TRUE |
| R78K | rna1-1 | -0.0711 | FALSE |
| T34E | msn5 | -0.0741 | FALSE |
| R78K | yrb1-51 | -0.0814 | FALSE |
| R78K | crm1_damp | -0.0839 | TRUE |
| G80A | msn5 | -0.0892 | TRUE |

**Supplementary Table 5** Interquartile range (IQR) of $log_2$(fold change) values across all the Gsp1 mutants for each prey protein identified. Ordered by IQR.

| Prey gene name | interquartile range (IQR) | Prey gene name | interquartile range (IQR) | Prey gene name | interquartile range (IQR) |
|---|---|---|---|---|---|
| Spa2 | 14.10 | Hef3 | 2.00 | Mph1 | 1.55 |
| Pup2 | 10.27 | Hsp60 | 1.99 | Rpl26a | 1.55 |
| Cdc3 | 9.65 | Rpa135 | 1.98 | Taf2 | 1.51 |
| Rna1 | 6.86 | Vps1 | 1.98 | Net1 | 1.51 |
| Mae1 | 6.34 | Rvb2 | 1.96 | Msh2 | 1.51 |
| Hrp1 | 6.25 | Yrb30 | 1.96 | Egd1 | 1.49 |
| Spb1 | 6.23 | Dep1 | 1.93 | Rpl29 | 1.47 |
| Adr1 | 5.92 | San1 | 1.92 | Hmo1 | 1.47 |
| Rgr1 | 5.83 | Frs1 | 1.91 | Tdh3 | 1.47 |
| Ecm1 | 5.69 | Rpc31 | 1.90 | Azf1 | 1.46 |
| Swi1 | 5.66 | Oca1 | 1.90 | Nop2 | 1.45 |
| Yar1 | 5.65 | Mtc1 | 1.89 | Rps0b | 1.44 |
| Cmr1 | 5.30 | Tti1 | 1.87 | Rpc82 | 1.43 |
| Acf4 | 5.26 | Yrb1 | 1.87 | Ssa1 | 1.43 |
| Vps71 | 5.25 | Ptc3 | 1.85 | Gbp2 | 1.42 |
| Kri1 | 5.12 | Sdd3 | 1.84 | Lcl2 | 1.42 |
| Lcp5 | 5.09 | Dbp2 | 1.80 | Rpp1a | 1.41 |
| Gcd14 | 4.99 | Tub1 | 1.79 | Mgm101 | 1.40 |
| Srp54 | 4.94 | Rpl39 | 1.78 | Gfa1 | 1.39 |
| Reh1 | 4.79 | Swc5 | 1.78 | Grs1 | 1.38 |
| Gcd10 | 4.79 | Ugp1 | 1.77 | Mcm4 | 1.38 |
| Tdh1 | 4.78 | Tif4631 | 1.77 | Puf6 | 1.38 |
| Srp68 | 4.75 | Aim36 | 1.77 | Rpl10 | 1.37 |
| Srp1 | 4.61 | Dbp3 | 1.76 | Tra1 | 1.37 |
| Rpc37 | 4.04 | Pln1 | 1.73 | Pro3 | 1.37 |
| Kap120 | 3.24 | Rpl37a | 1.73 | Nop4 | 1.35 |
| Pol2 | 3.23 | Svf1 | 1.72 | Tfc3 | 1.35 |
| Kap95 | 3.05 | Thi20 | 1.72 | Spt20 | 1.35 |
| Rix7 | 2.93 | Mcm6 | 1.70 | Rpl3 | 1.34 |
| Yef3 | 2.85 | Npa3 | 1.70 | Rpl33b | 1.33 |
| Rpb8 | 2.62 | Krs1 | 1.68 | Cdc9 | 1.33 |
| Gpn3 | 2.59 | Siw14 | 1.68 | Ubp15 | 1.32 |
| Rea1 | 2.56 | Cbf5 | 1.67 | Rpc11 | 1.31 |
| Paa1 | 2.55 | Zuo1 | 1.67 | Rpo21 | 1.31 |
| Apa1 | 2.53 | Rvb1 | 1.65 | Rlp24 | 1.31 |
| Eno1 | 2.49 | Wtm1 | 1.65 | Skn7 | 1.31 |
| Hpm1 | 2.48 | Vps13 | 1.64 | Hsp42 | 1.31 |
| Srm1 | 2.46 | Cdc14 | 1.64 | Cys4 | 1.30 |
| Scj1 | 2.44 | Dpb4 | 1.64 | Orc2 | 1.30 |
| Rtp1 | 2.44 | Yku70 | 1.63 | Hca4 | 1.29 |
| Tub3 | 2.34 | Fun12 | 1.62 | Ree1 | 1.29 |
| Idh2 | 2.30 | Pwp1 | 1.61 | Ssz1 | 1.27 |
| Idh1 | 2.20 | Rpc34 | 1.61 | Yta7 | 1.27 |
| Tub2 | 2.19 | Aco1 | 1.60 | Pre6 | 1.26 |
| Aro9 | 2.17 | Spt8 | 1.59 | Gtr2 | 1.26 |
| Krr1 | 2.17 | Orc1 | 1.58 | Hal5 | 1.25 |
| Cia2 | 2.14 | Pse1 | 1.58 | Rpb4 | 1.24 |
| Ade5,7 | 2.13 | Pdi1 | 1.57 | Nop6 | 1.24 |
| Ura7 | 2.08 | Rpa190 | 1.57 | Rpc25 | 1.23 |
| Afg2 | 2.02 | Sum1 | 1.56 | Muk1 | 1.23 |
| Yap1 | 2.01 | Yku80 | 1.56 | Caf40 | 1.23 |

| Prey gene name | interquartile range (IQR) | Prey gene name | interquartile range (IQR) | Prey gene name | interquartile range (IQR) |
|---|---|---|---|---|---|
| Aat1 | 1.23 | Stm1 | 1.00 | Rpb11 | 0.74 |
| Msh3 | 1.23 | Reb1 | 1.00 | Arp8 | 0.73 |
| Spt5 | 1.22 | Ioc4 | 0.99 | Rtt102 | 0.73 |
| Rok1 | 1.22 | Asg1 | 0.99 | Cdc1 | 0.72 |
| Swr1 | 1.21 | Ioc3 | 0.99 | Pob3 | 0.69 |
| Irc20 | 1.20 | Msn1 | 0.99 | Htz1 | 0.68 |
| Rpp2a | 1.20 | Adh6 | 0.99 | Spt15 | 0.68 |
| Rim1 | 1.20 | Rpc19 | 0.99 | Rsc58 | 0.67 |
| Hpc2 | 1.19 | Rpc53 | 0.98 | Hir1 | 0.64 |
| Mcm5 | 1.19 | Adh3 | 0.98 | Rfc3 | 0.62 |
| Rpl15a | 1.19 | Rbg1 | 0.98 | Hos3 | 0.61 |
| Rpa49 | 1.19 | Raf1 | 0.98 | Mot1 | 0.61 |
| Ret1 | 1.18 | Orc3 | 0.98 | Rsc8 | 0.61 |
| Tkl2 | 1.17 | Rfc1 | 0.96 | Arp4 | 0.60 |
| Hst1 | 1.16 | Srl2 | 0.96 | Pre2 | 0.60 |
| Rpl9a | 1.16 | Rpl24a | 0.96 | Ies2 | 0.60 |
| Elo1 | 1.16 | Top1 | 0.95 | Arp7 | 0.55 |
| Rtg3 | 1.16 | Rpl6b | 0.95 | Rpc10 | 0.54 |
| Rfm1 | 1.15 | Isw1 | 0.95 | Ant1 | 0.54 |
| Gdh1 | 1.14 | Sth1 | 0.94 | Abf1 | 0.54 |
| Sry1 | 1.13 | Nhp2 | 0.94 | Thi7 | 0.52 |
| Chd1 | 1.12 | Egd2 | 0.93 | Lsm6 | 0.51 |
| Top2 | 1.11 | Npl6 | 0.90 | Rfc5 | 0.49 |
| Rpl31b | 1.11 | Rps29a | 0.90 | Hir3 | 0.49 |
| Cst6 | 1.11 | Taf9 | 0.89 | Srp14 | 0.48 |
| Rpl36a | 1.10 | Gar1 | 0.89 | Rco1 | 0.45 |
| Rpl4a | 1.10 | Rsc4 | 0.89 | Rfc4 | 0.43 |
| Abf2 | 1.09 | Snf2 | 0.88 | Aim14 | 0.37 |
| Rpb5 | 1.09 | Taf14 | 0.88 | Sis1 | 0.33 |
| Spt7 | 1.08 | Grx1 | 0.87 | Aah1 | 0.00 |
| Orc4 | 1.08 | Rpl8b | 0.87 | Aim41 | 0.00 |
| Sin3 | 1.08 | Rsc6 | 0.87 | Arl3 | 0.00 |
| Rpo31 | 1.07 | Mog1 | 0.86 | Cfd1 | 0.00 |
| Nur1 | 1.07 | Rsc9 | 0.84 | Gcn3 | 0.00 |
| Rpb10 | 1.06 | Rsc3 | 0.84 | Lrs4 | 0.00 |
| Sko1 | 1.06 | Gcd1 | 0.83 | Opi1 | 0.00 |
| Rpp2b | 1.06 | Rfc2 | 0.83 | Rad5 | 0.00 |
| Rpo26 | 1.05 | Swi3 | 0.83 | Rpl21b | 0.00 |
| Vps72 | 1.05 | Ies5 | 0.82 | Rrp8 | 0.00 |
| Rpl30 | 1.04 | Ioc2 | 0.82 | Rrs1 | 0.00 |
| Hri1 | 1.04 | Imh1 | 0.81 | Sen1 | 0.00 |
| Nop10 | 1.03 | Oye2 | 0.80 | Slx9 | 0.00 |
| Pdr1 | 1.03 | Ies1 | 0.80 | Smc2 | 0.00 |
| Ald4 | 1.03 | Nhp10 | 0.79 | Snf12 | 0.00 |
| Yra1 | 1.03 | Arp9 | 0.79 | Snf5 | 0.00 |
| Nip7 | 1.02 | Spt16 | 0.77 | Spp41 | 0.00 |
| Prp43 | 1.02 | Sfh1 | 0.76 | Stb4 | 0.00 |
| Rtt106 | 1.02 | Htb2 | 0.76 | Sti1 | 0.00 |
| Hir2 | 1.02 | Enp2 | 0.76 | Sub1 | 0.00 |
| Arp5 | 1.02 | Taf10 | 0.76 | Tif4632 | 0.00 |
| Itc1 | 1.02 | Bur6 | 0.76 | | |
| Rpc40 | 1.01 | Isw2 | 0.75 | | |
| Ade3 | 1.01 | Rsc2 | 0.75 | | |
| Hho1 | 1.01 | Taf5 | 0.74 | | |
| Rpl5 | 1.00 | Ies3 | 0.74 | | |

**Supplementary Table 6** - Michaelis -Menten parameters of GAP-mediated GTP hydrolysis. The two Michaelis-Menten parameters and their ratio (enzymatic efficiency) are determined by an integrated Michaelis-Menten fit for each individual experiment. Standard error is based on the three or more replicates.

| Gsp1 mutant | $k_{cat}$ [$s^{-1}$] | std.error $k_{cat}$ [$s^{-1}$] | $K_m$ [μM] | std.error $K_m$ [μM] | $k_{cat}/K_m$ [$s^{-1}$ $\mu M^{-1}$] | std.error $k_{cat}/K_m$ [$s^{-1}$ $\mu M^{-1}$] |
|---|---|---|---|---|---|---|
| WT | 9.2 | 0.66 | 0.4 | 0.04 | 26.0 | 2.57 |
| T34A | 9.8 | 3.65 | 2.3 | 0.63 | 4.0 | 0.56 |
| T34E | 8.9 | 0.23 | 1.4 | 0.09 | 6.5 | 0.36 |
| T34G | 5.0 | 0.81 | 0.8 | 0.12 | 7.1 | 0.99 |
| T34L | 15.2 | 1.27 | 2.0 | 0.10 | 7.5 | 0.88 |
| T34Q | 5.4 | 0.20 | 2.2 | 0.26 | 2.5 | 0.23 |
| F58A | 8.6 | 0.57 | 0.2 | 0.03 | 35.8 | 2.97 |
| R78K | 4.3 | 0.73 | 2.1 | 0.59 | 2.4 | 0.35 |
| D79A | 11.9 | 2.21 | 3.6 | 1.11 | 3.8 | 0.62 |
| D79S | 4.1 | 0.32 | 1.7 | 0.23 | 3.0 | 0.59 |
| G80A | 8.8 | 0.14 | 0.3 | 0.01 | 28.8 | 1.56 |
| K101R | 8.2 | 1.22 | 0.2 | 0.01 | 44.7 | 9.20 |
| R108A | 7.8 | 0.32 | 0.2 | 0.01 | 42.0 | 4.14 |
| R108G | 9.2 | 0.16 | 0.1 | 0.01 | 82.3 | 5.74 |
| R108I | 13.2 | 2.24 | 3.1 | 0.66 | 4.3 | 0.15 |
| R108L | 5.2 | 0.63 | 0.3 | 0.07 | 19.3 | 2.87 |
| R108Q | 9.2 | 0.03 | 0.2 | 0.00 | 61.2 | 1.18 |
| R108Y | 7.8 | 1.39 | 0.2 | 0.07 | 40.1 | 6.34 |
| R112S | 4.9 | 1.28 | 3.0 | 1.01 | 1.7 | 0.20 |
| K132H | 6.7 | 0.45 | 5.6 | 0.13 | 1.2 | 0.06 |
| H141R | 7.2 | 1.19 | 0.1 | 0.02 | 56.3 | 3.04 |
| K143W | 9.5 | 0.86 | 0.1 | 0.02 | 71.8 | 3.48 |
| Q147E | 7.6 | 0.65 | 0.7 | 0.04 | 11.6 | 1.58 |
| Y157A | 8.8 | 1.89 | 0.2 | 0.03 | 57.7 | 4.87 |
| A180T | 4.0 | 0.49 | 0.4 | 0.04 | 11.1 | 0.29 |

**Supplementary Table 7** - Michaelis-Menten parameters of GEF-mediated nucleotide exchange. Standard error is based on the error of the Michaelis-Menten fit to the data.

| Gsp1 mutant | $k_{cat}$ [$s^{-1}$] | std.error $k_{cat}$ [$s^{-1}$] | $K_m$ [µM] | std.error $K_m$ [µM] | $k_{cat}/K_m$ [$s^{-1}$ $\mu M^{-1}$] | std.error $k_{cat}/K_m$ [$s^{-1}$ $\mu M^{-1}$] |
|---|---|---|---|---|---|---|
| WT | 3.0 | 0.08 | 0.9 | 0.12 | 3.3 | 0.44 |
| T34A | 1.8 | 0.10 | 0.9 | 0.22 | 2.1 | 0.55 |
| T34E | 1.7 | 0.07 | 1.0 | 0.17 | 1.7 | 0.29 |
| T34G | 2.5 | 0.14 | 1.4 | 0.28 | 1.8 | 0.39 |
| T34L | 2.0 | 0.11 | 1.6 | 0.35 | 1.2 | 0.27 |
| T34Q | 1.3 | 0.05 | 1.0 | 0.14 | 1.3 | 0.20 |
| F58A | 1.9 | 0.06 | 1.6 | 0.16 | 1.2 | 0.13 |
| R78K | 3.5 | 0.19 | 10.2 | 1.43 | 0.3 | 0.05 |
| D79A | 3.2 | 0.14 | 2.6 | 0.31 | 1.2 | 0.15 |
| D79S | 2.2 | 0.12 | 0.9 | 0.21 | 2.6 | 0.64 |
| G80A | 1.2 | 0.10 | 1.0 | 0.33 | 1.2 | 0.39 |
| K101R | 4.0 | 0.42 | 304.9 | 50.52 | 0.0 | 0.00 |
| R108A | 3.0 | 0.13 | 0.9 | 0.16 | 3.2 | 0.56 |
| R108G | 5.4 | 0.12 | 8.5 | 0.55 | 0.6 | 0.04 |
| R108I | 8.1 | 0.55 | 149.2 | 15.73 | 0.1 | 0.01 |
| R108L | 3.4 | 0.08 | 49.2 | 2.95 | 0.1 | 0.00 |
| R108Q | 3.8 | 0.10 | 8.7 | 0.64 | 0.4 | 0.03 |
| R108Y | 4.5 | 0.14 | 19.3 | 1.59 | 0.2 | 0.02 |
| R112S | 0.8 | 0.12 | 4.1 | 1.28 | 0.2 | 0.07 |
| K132H | 1.9 | 0.17 | 1.6 | 0.49 | 1.1 | 0.35 |
| H141R | 0.6 | 0.03 | 0.5 | 0.13 | 1.2 | 0.30 |
| K143W | 1.2 | 0.08 | 0.6 | 0.20 | 1.8 | 0.57 |
| Q147E | 1.9 | 0.07 | 1.4 | 0.18 | 1.4 | 0.19 |
| Y157A | 1.0 | 0.06 | 1.0 | 0.24 | 0.9 | 0.22 |
| A180T | 2.3 | 0.05 | 1.2 | 0.09 | 2.0 | 0.16 |

**Supplementary Table 8** Intrinsic GTP hydrolysis rate of wild type and mutant Gsp1. Standard deviation is based on the data from 3 or more replicates.

| Gsp1 mutant | intrinsic GTP hydrolysis rate [$s^{-1}$] | std.error of intrinsic GTP hydrolysis rate [$s^{-1}$] |
|---|---|---|
| WT | 2.5E-05 | 1.2E-06 |
| T34A | 7.4E-06 | 3.0E-06 |
| T34E | 8.7E-06 | 1.1E-06 |
| T34G | 2.0E-05 | 1.9E-06 |
| T34L | 1.8E-05 | 3.7E-07 |
| T34Q | 6.6E-06 | 3.0E-06 |
| F58A | 2.1E-05 | 2.7E-07 |
| R78K | 8.0E-06 | 3.9E-06 |
| D79A | 4.3E-05 | 1.2E-05 |
| D79S | 1.8E-05 | 2.9E-06 |
| G80A | 1.5E-05 | 7.3E-07 |
| K101R | 2.7E-05 | 2.1E-06 |
| R108A | 1.4E-05 | 4.9E-07 |
| R108G | 1.9E-05 | 1.2E-06 |
| R108I | 3.4E-05 | 8.8E-06 |
| R108L | 1.9E-05 | 9.4E-07 |
| R108Q | 1.9E-05 | 5.0E-07 |
| R108Y | 2.0E-05 | 2.4E-06 |
| R112S | 1.6E-05 | 5.9E-06 |
| K132H | 3.3E-05 | 4.9E-06 |
| H141R | 3.1E-05 | 8.8E-07 |
| K143W | 2.9E-05 | 7.6E-07 |
| Q147E | 1.6E-05 | 9.6E-08 |
| Y157A | 3.9E-05 | 5.5E-06 |
| A180T | 2.7E-05 | 1.4E-06 |

**Supplementary Table 9** Apparent Tm values estimated from the circular dichroism (CD) thermal melts. Mutants are ordered by apparent Tm.

| Gsp1 mutant | Apparent Tm / °C |
|---|---|
| R78K | 79 |
| G80A | 77 |
| T34G | 77 |
| R108Y | 77 |
| N105L | 77 |
| R108G | 77 |
| WT | 76 |
| T34L | 76 |
| K101R | 76 |
| R108Q | 75 |
| R108I | 74 |
| A180T | 74 |
| K132H | 74 |
| Q147E | 73 |
| R108L | 73 |
| K143W | 73 |
| D79S | 72 |
| R112S | 71 |
| H141I | 66 |
| H141V | 63 |
| H141R | 63 |
| Y157A | 63 |
